# Transcriptomic profiling reveals multiple mechanisms of insecticide resistance in *Aedes aegypti* from Angola

**DOI:** 10.64898/2026.05.21.723794

**Authors:** Henry A. Youd, Jocelyn M. F. Ooi, Abdullahi Muhammad, Mark J. I. Paine, Eric R. Lucas, Xavier Grau-Bové, Linda Grigoraki, Arlete Dina Troco, Ricardo Parreira, Carla A. Sousa, João Pinto, David Weetman

**Affiliations:** Department of Vector Biology, Liverpool School of Tropical Medicine, Pembroke Place, Liverpool, L3 5QA; Centre de Regulació Genòmica, Barcelona, Spain; Institute of Molecular Biology and Biotechnology, Foundation for Research and Technology Hellas (IMBB-FORTH), Crete, Greece; Programa Nacional de Controlo da Malária, Ministério da Saúde de Angola, Luanda, Angola; Global Health and Tropical Medicine, GHTM, Associate Laboratory in Translation and Innovation Towards Global Health, LA-REAL, Instituto de Higiene e Medicina Tropical, IHMT, Universidade NOVA de Lisboa, Lisbon, Portugal

## Abstract

Control of arboviruses remains heavily reliant on insecticide-based vector control targeting adult *Aedes aegypti*, especially during outbreaks, but the effectiveness of these tools can be compromised by insecticide resistance. While the mechanisms underlying resistance have been widely studied in Latin American and South East Asian *Ae. aegypti*, knowledge from African populations is limited, particularly regarding metabolic resistance. To address this knowledge gap, we sequenced the transcriptomes of *Ae. aegypti* collected in Angola, from both unexposed individuals and survivors of exposure to the organophosphate fenitrothion, alongside two insecticide-susceptible laboratory reference strains. Many overexpressed genes belonged to the major detoxification enzyme families, including 96 cytochrome P450 monooxygenases (CYP450s), 18 glutathione S-transferases (GSTs), and 35 carboxylesterases, with multiple genes previously detected as upregulated in Latin American and Asian populations. These included frequently reported, functionally-validated, metabolic resistance genes such as *CYP9J24*, *CYP9J26*, and *CYP6BB2*. However, expression of auxiliary resistance families including hexamerins, heat shock proteins, and odorant binding proteins were linked to the insecticide resistance phenotype, whilst numerous cuticular genes differentiated the Angolan population from both susceptible laboratory strains. A novel candidate, *CYP6AG7*, that was overexpressed after fenitrothion exposure was experimentally validated, and surprisingly metabolised fenitrothion into its toxic oxon form, which it did not subsequently break down. The antioxidant response element (ARE) motif, to which the transcription factor *Maf-S* binds, was detected in all CYP450 overexpressed in the fenitrothion treatment suggesting their potential coordinated induction. Analysis of genetic differentiation revealed several resistance-linked genes under potential selection, and SNP screening identified both known and novel non-synonymous mutations in the voltage-gated sodium channel (*VGSC*) gene, the target for pyrethroid insecticides. This is the first RNAseq dataset for *Ae. aegypti* from Africa in the context of insecticide resistance, providing insight into the complexity of resistance mechanisms, including some shared, and others potentially novel, compared to better studied populations from other geographical regions.

**Author summary:** Dengue, chikungunya, yellow fever, and Zika are diseases that exert an increasing public health burden across Africa, primarily transmitted by the mosquito *Aedes aegypti*. We rely heavily on insecticides to control these mosquitoes, but populations are increasingly developing resistance, making control efforts less effective. While resistance mechanisms have been well-studied in the Americas and Asia, comparatively little is known about how African *Ae. aegypti* resist insecticides. We collected *Ae. aegypti* mosquitoes from Angola and compared the genes expressed in fenitrothion resistant versus susceptible mosquitoes using RNA sequencing. We identified overexpressed candidate insecticide resistance genes from the detoxification enzyme families that mosquitoes use to break down insecticides, including several genes previously linked to resistance in other regions. One novel enzyme identified, CYP6AG7, was experimentally validated and found to convert the pro-insecticide fenitrothion into its harmful form but interestingly not break down this harmful metabolite further. Mutations potentially linked to insecticide resistance were also detected in detoxification genes and insecticide target site genes. Our study provides the first comprehensive molecular characterization of insecticide resistance mechanisms in African *Aedes aegypti,* offering crucial data to inform vector control strategies and insecticide resistance management across the continent as dengue and related diseases continue to spread.

## Introduction

The major *Aedes*-borne diseases, dengue, chikungunya, yellow fever and Zika, exert a huge public health burden across the tropics. The impact of dengue is of particular concern, with an estimated 390 million cases every year [1] and approximately half the world’s population now at risk of transmission [2]. *Aedes aegypti* is usually the principal vector of these arboviral diseases [3,4]. With currently available arboviral vaccines not fully effective and limited to certain epidemiological settings, or supply-limited, suppressing populations of *Ae. aegypti* represents the main disease control measure [5]. Alongside larval source management [6], chemical interventions against adults constitute the majority of *Ae. aegypti* control [7–9].

Insecticide use has long been the cornerstone of *Ae. aegypti* control [10], particularly in arboviral outbreak scenarios [11,12]. While mosquito bednets are commonly used against many mosquito species, they are relatively ineffective against *Ae. aegypti* due to their crepuscular peak biting times [13]. Space spraying/fogging of peridomestic areas [14], and especially indoor residual spraying (IRS) [15], have been shown to be effective in some settings. IRS exploits *Ae. aegypti*’s endophilic, indoor resting behaviour which is common in many New World populations [16–19], but efficacy in Africa is uncertain owing to more exophilic resting behaviour [13,20]. The insecticides used for control of *Ae. aegypti* adults are primarily carbamates, pyrethroids, and organophosphates (OPs), particularly the latter two for space spraying [21–23]. Unfortunately, resistance to each class has developed in *Ae. aegypti* in the Americas, Asia, and Africa [24] with notable effects on control [25]. Insecticide resistance in *Ae. aegypti* is thought to operate mainly through two mechanisms - metabolic detoxification and target site mutations, which can occur either independently or in tandem [26–28]. Other mechanisms such as reduced cuticular penetration and/or sequestration of insecticides may also play a role but are far less well studied [29–31].

Metabolic detoxification involves a series of different gene families. The best known are the cytochrome P450 monooxygenases (CYP450), which are a large and diverse family of enzymes that metabolise harmful xenobiotics via oxidation in the presence of NADP+-CYP reductase (CPR) and often cytochrome b5 [32,33]. Upregulation of CYP450s, alongside mutations and copy number variants in these genes, have been implicated in resistance to DDT, pyrethroids, carbamates, and OPs, including cross-resistance [34–39]. Several *Ae. aegypti* CYP450s in the CYP6 and CYP9 families have been shown experimentally to metabolise insecticides including *CYP9J24*, *CYP9J26*, and *CYP9J28* against deltamethrin and permethrin, *CYP9J32* against chlorfenapyr, and *CYP6Z8* against pyriproxyfen [40–42]. Glutathione S-transferases (GST) are another large family of multifunctional enzymes which have been reported to both directly conjugate xenobiotics with glutathione to produce excretable water-soluble compounds and assist with metabolism of secondary products produced by other detoxifying enzymes such as CYP450s [43,44]. GSTs have similarly been associated with resistance to DDT, pyrethroids, carbamates, and OPs, with *GSTE2* and *GSTE5* specifically shown to metabolise DDT [24,43,45–47]. Moreover, upregulation and copy number variation of carboxylesterases (COE) has been linked to carbamate, OP, and pyrethroid resistance through catalysing the hydrolysis of ester bonds present in insecticides [24,48–51]. COEs are also capable of sequestering insecticides, followed by slow metabolism. In *Ae. aegypti*, one such enzyme, *COEAE3A*, has been shown to metabolise the oxon form of the larvicide temephos, and confer OP, carbamate, and pyrethroid resistance when transgenically over- expressed in *Anopheles gambiae* [50,52]. Expression of genes in these detoxification families is thought to be regulated in insects by the antioxidant response element (ARE) motif, through which the transcription factors *CncC* and *Maf-S* bind upon insecticide exposure [53]. For example, in *An. gambiae*, *CYP6M2* and *GSTD1* have been demonstrated to be regulated by *Maf-S*, modulating DDT and pyrethroid resistance [54]. Other *Ae. aegypti* detoxification candidate genes, such as UDP-glycosyltransferases (UGT), ATP-binding cassette (ABC) transporters, alcohol dehydrogenases, short-chain reductase/dehydrogenases (SDR), and heme peroxidase have also been associated with resistance phenotypes [55–59], though experimental validation is lacking.

Odorant binding proteins (OBPs), which transport odorants to sensory receptors [60,61], have also been implicated as a potential resistance mechanism in mosquitoes [31,62–64], through sequestration of the insecticide preventing it from reaching the intended binding site.

Target site mutations are most commonly point mutations within the primary insecticide target, which reduce binding affinity and can confer resistance. In mosquitoes this is known to occur within the voltage gated sodium channel (*VGSC*), acetylcholinesterase 1 (*ACE-1*), and GABA receptor (*RDL*), though *VGSC*-based resistance to pyrethroids, appears by far the most ubiquitous of these in *Ae. aegypti* [65,66]. Numerous amino acid substitutions (*kdr* mutations) in the four domains of the *VGSC* that induce sodium channel insensitivity have been recorded in *Ae. aegypti*, including V410L, S723T, G923V, L982W, S989P, I1011M/V, V1016G/I, T1520I, F1534S/L/C, D1763Y [24,67]. Some of these *kdr* mutations can co-occur in haplotypes which confer an elevated level of phenotypic resistance, for example, 989P/1016G/1534C in Asia and 410L/1016I/1534C in the Americas [68,69].

In African *Ae. aegypti,* beyond studies of known *VGSC* target site mutations [70–75], there is very limited knowledge of resistance mechanisms and associated genes, which reduces capacity for design of insecticide resistance management (IRM) programmes [76]. Discovery studies for non-target site mechanisms typically involve transcriptomic analyses, but to date there have been no published RNAseq datasets for African *Ae. aegypti* in the context of insecticide resistance. Within Africa, Angolan populations are of particular interest, being noteworthy for their markedly admixed ancestry between sylvatic African forms (*Ae. aegypti formosus*) and the globally invasive domestic subspecies (*Ae. aegypti aegypti*), which shows elevated capacity for transmission of arboviral disease [77–84] Recent genomic research has also revealed the presence of high frequency insecticide resistance mutations in Angola, with admixture-aided introduction through recent gene flow from the Americas back to Angola and other African urban populations [84]. Collectively the spread of *Ae. aegypti aegypti* ancestry and resistance-associated alleles through African cities has likely contributed to the current emergence of dengue and other *Ae. aegypti*-transmitted diseases across the continent [85]. Here we use RNAseq for discovery of genes associated with resistance in *Ae. aegypti* from Angola, followed by *in silico* and *in vitro* validation approaches.

## Methods

### Mosquito sampling and resistance phenotyping

An entomological survey was carried out in Luanda Province, Angola, between October and November 2016 in which *Ae. aegypti* mosquitoes were collected by larval sampling and ovitraps. Mosquitoes were reared to adults, with mixing of the emerged mosquitoes from the same cage to reduce relatedness in the subsequent tests.

Bioassays were performed on 3-5-day old non-blood fed female mosquitoes using fenitrothion (1%), malathion (5.0%), bendiocarb (0.1%), DDT (4.0%), permethrin (0.75%) and deltamethrin (0.05%) impregnated papers, with 60-minute exposure time, according to WHO kits and protocols [86]. For each assay, controls consisted of mosquitoes unexposed to insecticides but to papers impregnated with the insecticide excipient. Phenotyped *Ae. aegypti* individuals were preserved in 0.5 ml microtubes with RNALater (ThermoFisher Scientific), transported on dry ice and stored at −20°C until further analysis. The overall study methodology is shown in Fig 1.

**Fig 1.**
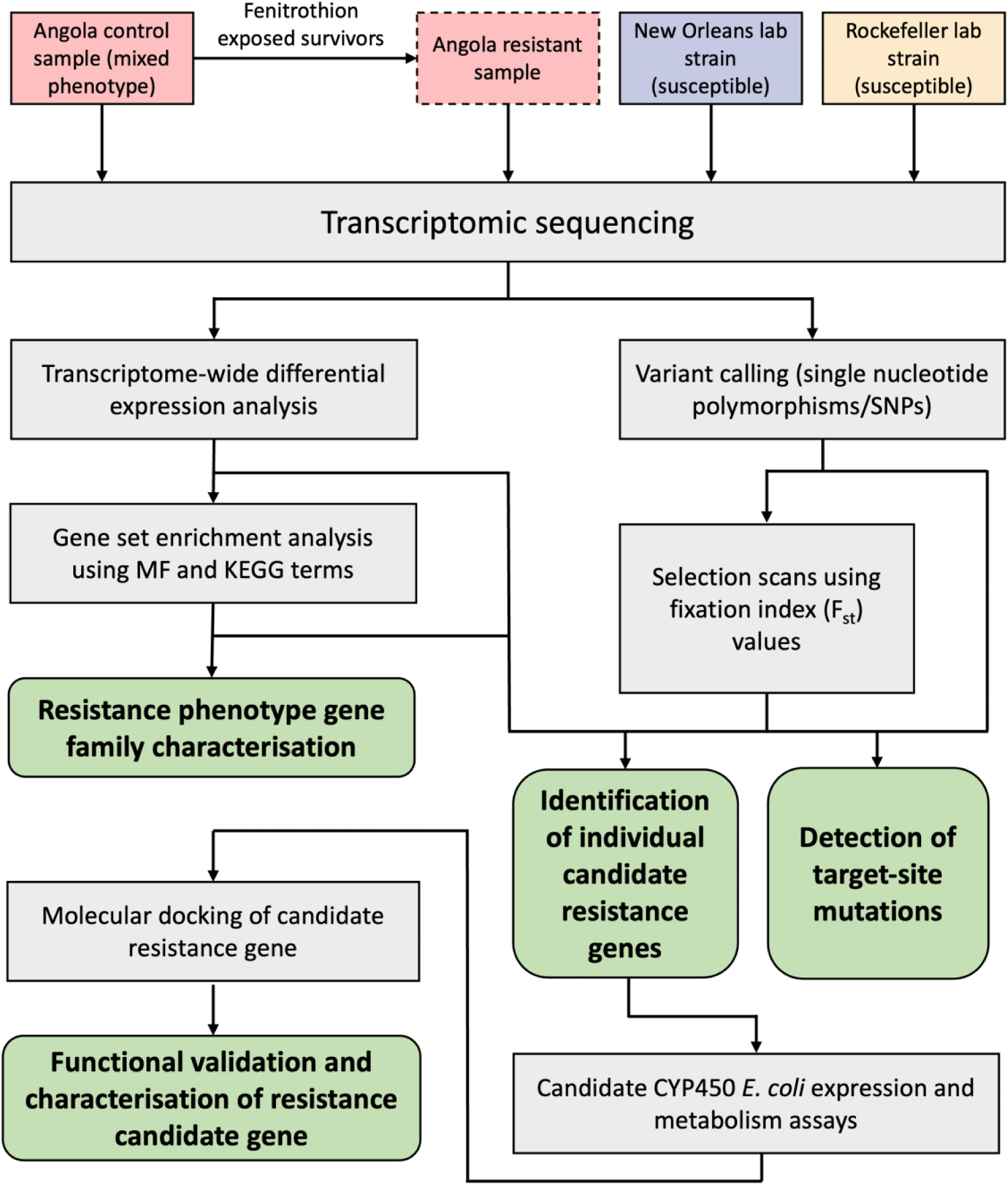
The methodology and workflow of the study.

### RNA isolation and sequencing

RNA extraction and isolation was performed using the PicoPure RNA Isolation Kit (ThermoFisher Scientific) according to manufacturer instructions. RNA was sampled from four groups: Angola control (i.e. insecticide unexposed), Angola fenitrothion resistant (fenitrothion exposed survivors), and two insecticide-susceptible lab strains, New Orleans and Rockefeller held at the Liverpool School of Tropical Medicine. Henceforth, these groups are referred to as CON, FEN, NWO, and ROC, respectively. Three pools of N = 5 each were sequenced for each group. Sequencing was conducted at the Centre for Genomic Research (CGR) at the University of Liverpool, following poly-A enrichment to reduce rRNA present in the pools, to produce 2x150bp paired-end reads.

### RNAseq differential expression analysis

Raw pair-end FASTQ reads were inputted into the Snakemake pipeline RNA-Seq-Pop [87] for analysis, which analyses read counts and genetic polymorphisms. All software and steps used in this study (excluding phylogenetic analysis, gene set enrichment analysis, protein modelling, and other data manipulation and visualisation using Microsoft Excel v16.103.1, Python v3.1.1, and R v4.2.1) as mentioned in the below sections were included as part of this workflow.

Raw RNAseq reads were first subjected to sequence quality control using FastQC v0.11.9 (http://www.bioinformatics.babraham.ac.uk/projects/fastqc/). High quality reads were then mapped to the annotated reference *Ae. aegypti* transcriptome AaegL5.3 (.fa) taken from VectorBase [88,89] using Kallisto v0.46.1 [90], with a k-mer length of 31, using 100 bootstrap samples, and transcript abundance quantified. Genome-wide analysis was then performed to determine differentially expressed genes (DEGs), following mapping of transcripts to genes, by DESeq2 v1.30.1 [91] with size factor normalised read counts. Differential expression was calculated for three comparisons. Comparisons between NWO vs CON and ROC vs CON were used to identify upregulated genes that comprised part of the natural variation in the Angolan *Ae. aegypti* population and were constitutively upregulated, as well as to enable comparison to other studies. Because this Angola population exhibits multiple and often strong resistance phenotypes (in the bioassays) to several insecticides, and because both NWO and ROC have been held in colony for many years, many of the DEGs will be unrelated to resistance phenotypes. Thus, the CON vs FEN comparison was used as an additional filter to identify differentially expressed genes more confidently involved in phenotypic fenitrothion-specific resistance. A gene was considered up or down regulated if the log2 fold change (FC) was > 1 or < −1, respectively, and if the adjusted p value was < 0.05. P-values were obtained using a Wald test and adjusted for multiple testing to control the false discovery rate at 5%. As additional quality control steps, a principal component analysis (PCA) plot and a sample-to-sample correlation heatmap were produced using normalised read count results from DESeq2 to check clustering of biological replicates.

Significantly upregulated genes were taken forward for gene set enrichment analysis (GSEA) using g:Profiler [92] to provide insight into functionally-related groups of upregulated genes. GSEA was performed using molecular function (MF) and KEGG terms [93,94]. Multiple testing correction was performed using the g:SCS threshold from g:Profiler.

DEGs were also specifically filtered for those in detoxification-related gene families (CYP450s, GSTs, COEs, UGTs, ABCs etc.), and also those within gene families previously implicated in auxiliary resistance mechanisms such as cuticular-related genes, hexamerins, heat shock proteins, and odorant binding proteins [29,95]. Because many genes were annotated as “unspecified product” according to VectorBase description, InterPro domains were instead used to examine these gene families. This was performed according to InterPro domains which contained the terms: Cytochrome P450", "GlutathioneS-transferase", "Carboxylesterase", "UDP-glucuronosyl/UDP-glucosyltransferase", and "ABC transporter”, for detoxification-related genes, "AMP-dependent synthetase/ligase", "Thioesterase;Phosphopantetheine binding ACP domain", "Acyl-CoA desaturase", "ELO family", "Fatty acyl-CoA reductase", and "cuticle protein" for cuticular-related genes, along with "hexamerin", "heat shock protein", "odorant binding/odorant-binding protein".

### Phylogenetic analysis of CYP450s

CYP450s that were significantly overexpressed in any comparison were compared using phylogenetic analysis to visualise potential clustering of DEGs within subfamilies, and relationships with known insecticide metabolisers. The amino acid sequence of each CYP450 was extracted from VectorBase and then aligned using MUSCLE [96] within MEGA v11.0.10 [97] using default options. A phylogenetic tree was then built in IQ-TREE3 v3.0.1 [98] from the aligned sequences according to maximum likelihood, using the best-fit model according to BIC (LG+I+R5 substitution model), with 1000 bootstraps.

### Single nucleotide polymorphism and selection analyses

Raw reads also were aligned to the reference *Ae. aegypti* genome assembly AaegL5.3 (.fa), again taken from VectorBase, using HISAT2 v2.2.1 [99], with duplicate reads marked by samblaster v0.1.26 [100], to produce binary alignment files (BAM). The resulting BAMs were then indexed using SAMtools v1.19 [101] and single nucleotide polymorphisms (SNPs) called using freebayes v1.3.2 [102] for Bayesian, haplotype-based, variant detection, with the options --pooled-discrete --ploidy 10 (2 per individual in each sample pool) with default quality filters to produce variant call format (VCF) files. SNPs were called with each of the four different groups as separate populations. Called SNPs were then annotated, and the functional effect of each variant predicted using SnpEff v5.0 [103].

SNPs were filtered for selection analysis by applying a quality score of 30 and using a missingness proportion of 1, resulting in no missing allele calls (reference or alternative allele must be present). Filtered SNPs were used to compute the median fixation index Fst [104,105] for each protein-coding gene using scikit-allele v1.2.1 [106]. The difference between the modal per gene Fst and smallest binned Fst value (bins = 50) in each comparison was then calculated, and genes were considered of interest if their Fst value was greater than two interquartile ranges of this difference above the mode (Fig S2).

Genes that were both upregulated according to differential expression analysis and exhibited significant Fst values in the CON vs FEN comparison were taken forward for SNP analysis. Allele frequencies of every called non-synonymous variant in each group were calculated from VCF files following splitting of multiallelic sites, left-normalisation, and filtering using a quality score of 30 and a missingness threshold of 0.5. Genes exhibiting upregulation and significant Fst in the NWO vs CON or ROC vs CON comparisons were also reported, but analysis was not taken further in these groups. Nicotinic acetylcholine receptor subunits (nAChR) were also checked for SNPs, due to their role in neonicotinoid resistance, if they exhibited significant Fst in the CON vs FEN comparison, regardless of expression. Variants in these genes were reported only if they were present in the CON or FEN groups, and absent in the susceptible groups. Amino acid positions were reported according to the first transcript annotated with “missense” by SnpEff for each gene. In addition, the focal target site genes *VGSC*, *ACE-1*, and *RDL* were each checked for SNPs, regardless of missingness. Called SNPs within VGSC were checked against the standard *Musca domestica* notation by aligning all 13 *Ae. aegypti VGSC* transcripts (AAEL023266) to the *M. domestica* VGSC transcript (MDOA002080-RB) taken from VectorBase and aligned using Clustal Omega v1.2.4 [107]. *VGSC* SNP location is numbered according to *M. domestica*, with *Ae. aegypti* amino acids retained. The location according to the *Ae. aegypti VGSC* (AAEL023266-RL) is also provided to aid comparison in studies using *Ae. aegypti* codon numbering. Approximate domain location was determined using Davies *et al.* [108] (File S1).

### Recombinant CYP450 expression

A previously unstudied resistance gene candidate (*CYP6AG7*) identified in the analysis was experimentally validated via recombinant expression and subsequent metabolism assays. Additionally, the known *Anopheles gambiae* pyrethroid and pyriproxyfen metaboliser *CYP9J5* was used as a positive control, following prior successful recombinant expression and metabolism [109,110]. The nucleotide sequence of *CYP6AG7* was taken from the reference genome AaegL5.3 in VectorBase and optimised to include restriction sites for NdeI and NaeI at the 5’ end and SacI at the 3’ end, followed by insertion into the modified *Escherichia coli* expression vector pCW-OmpA2 [111] by Eurofins Genomics (Ebersberg, Germany), termed pCW:P450. A synonymous mutation (TCA to TCG) was introduced for serine coding at position 1014 in *CYP6AG7* to remove an NdeI binding site. DNA sequencing was performed by Eurofins to check for errors. The *An. gambiae* cytochrome P450 reductase plasmid (pACYC:AgCPR) was also produced as described in McLaughlin et al [111]. pCW:P450 and pACYC:AgCPR contain ampicillin and chloramphenicol resistance genes for selection, respectively. Expression for both constructs were under control of a tactac promoter for IPTG induction in cell cultures. Further, *An. gambiae* cytochrome b5 was produced to supplement enzyme reactions as described previously [112].

Competent *E. coli* DH5α cells were co-transformed with pCW:P450 and pACYC:AgCPR plasmids via heat shock and selected on Luria-Bertani (LB) agar plates with 20 μg/ml chloramphenicol and 100 μg/ml ampicillin and incubated at 37°C overnight. Colonies were then picked to undergo PCR and gel electrophoresis for additional plasmid confirmation. Positive colony samples were used to produce 3ml of LB broth starter culture in replicate for each gene, cultured at 37°C overnight with 200 rpm orbital shaking. 2 ml of starter culture were used to inoculate 200 ml of Terrific Broth in duplicate, again with chloramphenicol and ampicillin at stated concentrations. Cultures were incubated at 37°C with 200 rpm orbital shaking and monitored until early log-phase growth (A595 = 0.8–1.0) was reached. At this point the cultures were cooled to 25°C and 1 mM IPTG and 0.5 mM ALA (final concentrations) were added to induce expression, and incubation continued at 25°C with 150 rpm orbital shaking. CYP450 expression levels were monitored every two hours for up to approximately 20 hours. Expression was estimated by resuspending whole cells in Spectrum Buffer (0.1 M potassium phosphate at pH 7.4, 20% (v/v) glycerol, 6 mM magnesium acetate, 0.1 mM dithiothreitol) [113], adding approximately 1 mg/ml of sodium dithionite as a reducing agent and exposing to carbon monoxide for 1 min. Recording the absorption spectra (400-500 nm) change and examining the peak heights at 450 nm and 490 nm was used to determine CYP450 concentration [114]. Recombinant CYP450s were harvested ∼20h post induction and membranes prepared as described by Pritchard *et al.* [113]. Membranes were then isolated using ultracentrifugation for 1 h at 180,000 g and homogenized using a Dounce homogenizer in approximately 1 ml TSE buffer (50 mM Tris-acetate, pH 7.6, 250 mM sucrose, 0.25 mM EDTA) per 200 ml culture processed. Membrane samples were tested for CYP450 content and quality by 100-fold dilution as previously [114] Samples were stored as aliquots at below −70°C.

### Metabolism assays

A 2 mM stock concentration of insecticide was prepared in acetonitrile. Standard reactions contained 10 µM insecticide, 0.1 µM CYP450, and 0.8 µM cytochrome b5 in 0.1 M potassium phosphate at pH 7.4 with NADPH generating components (1 mM glucose-6-phosphate, 1U/ml glucose-6-phosphate-dehydrogenase, 0.25 mM MgCl2, and 0.1 mM NADP+), or without NADPH generating components for negative control reactions. Samples were incubated for 2 h at 30°C with 1200 rpm orbital shaking. Samples were quenched by adding 200 µl of acetonitrile and incubated for 5 min before being centrifuged at 13000 rpm for 15 min. 130µl of supernatant was transferred to HPLC vials and analyzed within 24 h. The retention times and absorbance of fenitrothion and deltamethrin were 5.40 min and 268 nm, and 9.14 min and 226nm, respectively. The difference in HPLC peak area between +NADP+ and -NADP+ samples was used to estimate insecticide depletion (2 technical replicates, 1 biological replicate). A metabolite peak, M1, was confirmed as the fenitrothion-oxon with retention time (3.7 min) identical to a fenitrothion-oxon standard. Standard reactions were used to assess the metabolism of the fenitrothion-oxon with differences as follows; 7.66 µM metabolite was used with 1 µM CYP6AG7 and 0.1 µM CYP9J5.

### Time-course assays

Time-course assays were performed to assess M1 production and to determine the period of linearity for the kinetics study. For CYP9J5, 0.1 µM enzyme was incubated with 20 µM fenitrothion with cytochrome b5 and NADP+, and samples were collected at 0, 2, 5, 10, 20, 30, and 60 min (n = 1), including a -NADP+ control at 60 min. For *CYP6AG7*, 0.2 µM enzyme was used with 100 µM fenitrothion and sampled at the same timepoints with a 60 min -NADP+ control (n = 1).

### Enzyme kinetics

Kinetic assays were conducted to determine the Km, Kcat, and Vmax values for fenitrothion-oxon production. For CYP9J5, 0.1 µM enzyme was incubated for 20 min with a concentration range of fenitrothion (0.8 µM to 100 µM) and -NADP+ controls at each concentration (2 technical replicates, 2 biological replicates). For *CYP6AG7*, 0.2 µM of enzyme was used with the same substrate range and incubated for 60 min, with -NADP+ controls at the highest and lowest concentrations (2 technical replicates, 2 biological replicates).

### Molecular docking

Molecular docking was conducted to examine the interaction between *CYP6AG7* (and *CYP9J5*) and fenitrothion. Docking was also performed against deltamethrin for comparison of binding affinities and with the metabolism assays, but the residue specific interactions was not characterised further. The translated amino acid sequence for each CYP450 was inputted into AlphaFold 3 [115], along with a heme group as a ligand, to produce protein models in CIF format. Per-atom confidence estimate was generally very high across the predicted models with > 90 predicted local distance difference test (pLDDT) [115] across the models. The Cys residues at position 442 in *CYP6AG7* and position 485 in *CYP9J5* were manually converted to Cym to indicate a deprotonated cysteine thiolate, which is bound to the heme iron atom, with a negatively charged thiol group. Similarly, models for deltamethrin, fenitrothion and the fenitrothion-oxon were taken from PubChem in SDF format [116]. Models were then accordingly passed to the mk_prepare_receptor.py and mk_prepare_ligand.py scripts from Meeko [117], to add partial charges. AutoDock Vina v1.2.0 [118] was then used to dock each insecticide with each receptor with exhaustiveness and number of modes set at 256 and 40, respectively. The search space was centred for both receptors and the grid dimensions were set at 25 angstroms across the X, Y, and Z axes. Considering the metabolic pathway from fenitrothion to the fenitrothion-oxon through oxidative desulphurisation in CYP450s [119], the best scored poses in which the sulphur atom was facing toward the heme iron was reported, and visualised using Mol* [120]. Binding pocket residues exhibiting hydrophobic contacts with fenitrothion were determined by LigPlot+ v2.3 [121] with default parameters. Interactions between these residues and fenitrothion were also calculated according to PLIP [122] with maximum hydrophobic interaction distance set at 5Å.

### Maf-S Antioxidant Response Element (ARE) motif scan

The presence of the antioxidant response element (ARE) motif to which *Maf-S* binds was checked across six CYP450s upregulated across all comparisons (and one GST) by examining the flanking region 3000 bp either side of each gene. This was performed using the R package MotifDb v1.40.0 [123], to check for the cnc-Maf-S *Drosophila melanogaster* motif using the frequency matrix taken from JASPAR (MA0530.1) [124]. Returned motifs with at least an 80% identity match were manually checked for conformation to the motif 5’-nnTGnnnnnnCnnnn-3’ as observed in *An. Gambiae* mosquitoes by Ingham *et al.* [54].

### Data availability

Raw RNA reads were deposited in the Sequence Read Archive under the BioProject number PRJNA1460739.

## Results

### Bioassays

Bioassays showed a 72.5% (N = 91) mortality rate to fenitrothion indicating presence of resistance in the population (Fig S1). Although fenitrothion resistance was the focus of the subsequent analyses, the Angolan population also exhibited strong resistance to deltamethrin (mortality = 7.3%, N = 109), permethrin (mortality = 2.6%, N = 116), bendiocarb (mortality = 10.4%, N = 116) and DDT (mortality = 0%, N = 92), but susceptibility to malathion (mortality = 100%, N = 99) (Fig S1).

### Differential expression of resistance-associated genes

Fenitrothion resistant mosquitoes were selected for RNA sequencing. This was firstly due to the fact that organophosphate insecticides have widespread uses in vector control programs, through IRS and ULV spraying applications. Secondly, organophosphate resistance is a rarely observed phenotype in African *Ae. aegypti* [13,125], and thus fenitrothion represents a model compound to investigate organophosphate resistance.

Of the 19,764 genes with detected transcripts, 7,925 (40.10%) genes were differentially expressed in the CON group compared to the NWO group, and 7,619 (38.55%) compared to the ROC group; with 4,602 (23.28%) differentially expressed in the FEN group compared to the CON group. Clustering of biological replicates in each group according to PCA, and expression correlation, was seen as expected (Fig S3). There was a large degree of overlap in significantly differentially expressed genes (DEGs) between the three comparisons; however, when considering commonality in the direction of expression, the number of shared genes dropped substantially (Fig 2).

**Fig 2.**
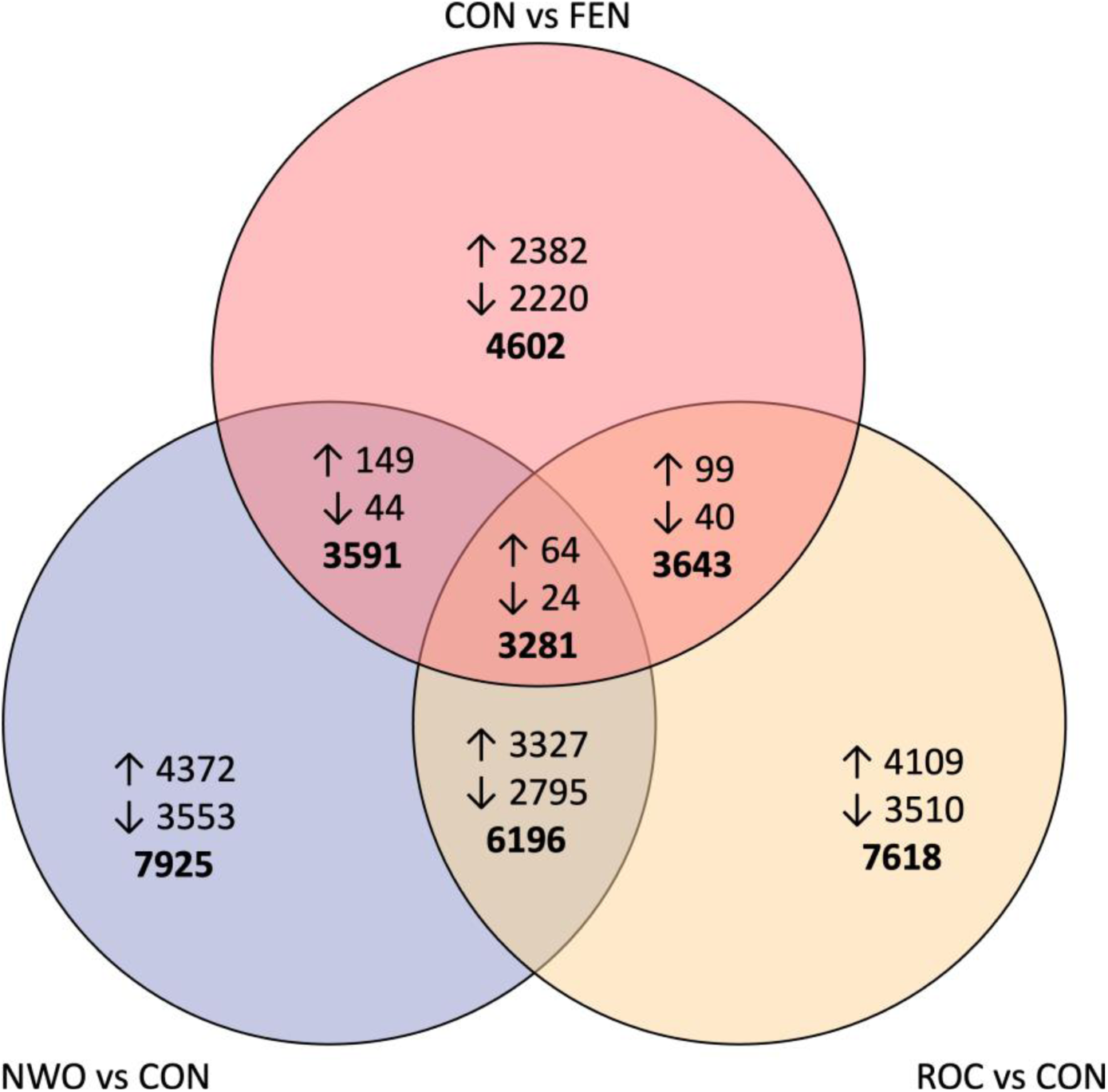
Venn diagram showing DEGs, according to log2 fold change (> 1 or < −1) and adjusted p value (< 0.05), in each of the three comparisons individually and shared between comparisons. Numbers next to arrows indicate the number of up- or downregulated genes, in which the directionality of expression had to be the same. Bold numbers indicate the number of DEGs whilst not considering such directionality

Gene set enrichment analysis using significantly upregulated genes revealed over representation of several cuticular-related terms in both the NWO and ROC vs CON comparisons (File S2). These included structural constituent of cuticle (GO:0042302, 94 genes vs NWO, 99 genes vs ROC), and chitin binding vs NWO (GO:0008061, 51 genes), as well as fatty acid elongase activity (GO:0009922, 15 genes in both comparisons). Several terms potentially linked to insecticide metabolism were also enriched including oxidoreductase activity (GO:0016491, 251 genes in both comparisons), heme binding (GO:0020037, 79 genes vs NWO, 78 genes vs ROC), monooxygenase activity (GO:0004497, 76 genes vs NWO, 77 genes vs ROC), serine hydrolase activity (GO:0017171, 133 genes vs NWO only), and glucuronosyltransferase activity (GO:0015020, 17 genes in both comparisons). Terms relating to transmembrane transporter activity and respiration were also seen in both susceptible strain comparisons. For the CON vs FEN comparison, all enriched terms were exclusively related to cell damage and DNA repair. All enriched terms can be found in File S2.

Among genes significantly over-expressed in all any comparison, 212 were members of detoxification families according to VectorBase or InterPro domains (96 CYP450s, 18 GSTs, 35 COEs, 35 ABCs, 28 UGTs; Fig 3 and File S3), alongside an additional 11 hexamerins, nine heat shock proteins, and 41 OBPs. Furthermore, 177 genes involved in cuticular modification were upregulated in any comparison; 66 cuticular hydrocarbon synthesis genes and 111 cuticle protein encoding genes (Fig 3 and File S3) consistent with results from the GSEA above. Genes that were significantly upregulated across all comparisons included CYP450s *CYP304B2*, *CYP6AG7*, *CYP6BB2*, *CYP9J24*, *CYP9J26*, AAEL029064 (putative *CYP9J27 [126]*), an ABC transporter (AAEL005043), a UGT (AAEL003098), a hexamerin (AAEL011169), and and *OBP56e* (Table 1). Phylogenetic analysis of the translated protein sequences of all significantly overexpressed CYP450s demonstrated clustering according to subfamilies, with patterns of expression observed between these groups (Fig S4). For example, upregulation of the CYP4H family appeared characteristic of the CON vs FEN comparison, whilst the CYP325 family was characteristic of the NWO vs CON and RO vs CON comparisons, and the CYP6 and CYP9J families broadly showed similar patterns of expression across comparisons.

**Fig 3.**
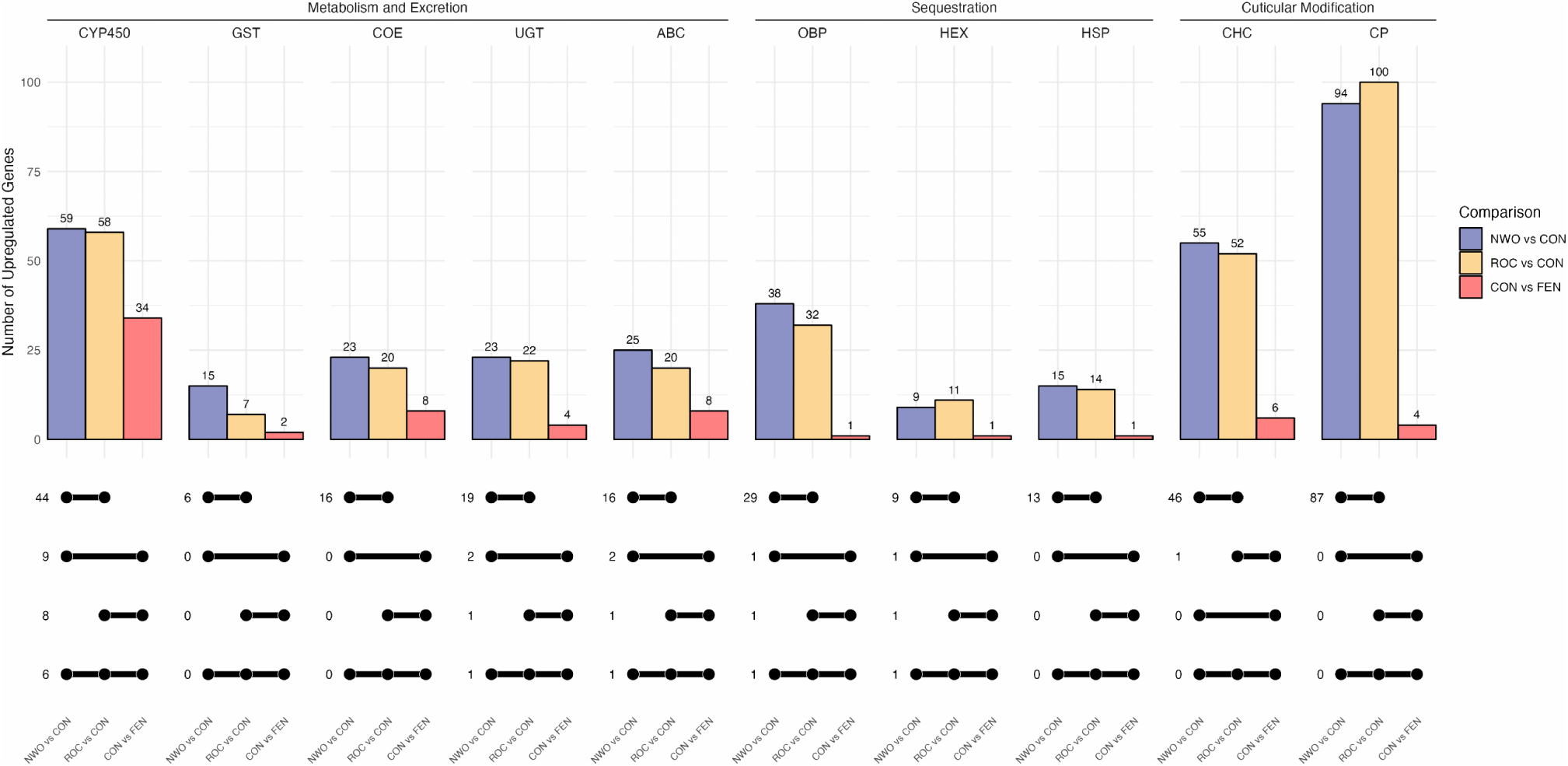
Number of upregulated insecticide resistance candidate genes in each gene family previously implicated with insecticide resistance in each comparison, and number of upregulated genes shared across comparisons.

**Table 1.** log2 fold change of insecticide resistance candidate genes significantly overexpressed in all three comparisons (log2 fold change > 1 and adjusted p value < 0.05). Genes in families previously implicated with insecticide resistance are shown. Values are rounded to two decimal places. Gene descriptions are taken from VectorBase according to community annotation.

| Gene Family | Gene ID | Gene Description | NWO vs CON | ROC vs CON | CON vs FEN |
| --- | --- | --- | --- | --- | --- |
| CYP450 | AAEL014412 | CYP304B2 | 3.95 | 2.86 | 2.27 |
|  | AAEL006989 | CYP6AG7 | 2.38 | 2.57 | 1.59 |
|  | AAEL014893 | CYP6BB2 | 3.38 | 2.44 | 1.49 |
|  | AAEL014613 | CYP9J24 | 7.66 | 5.03 | 2.55 |
|  | AAEL014609 | CYP9J26 | 5.17 | 2.26 | 1.44 |
|  | AAEL029064 | Cytochrome P450 | 5.15 | 1.87 | 1.05 |
| ABC | AAEL005043 | ABC Transporter | 1.78 | 1.00 | 1.44 |
| UGT | AAEL003098 | UDP-glycosyltransferase | 1.39 | 1.19 | 1.03 |
| Hexamerin | AAEL011169 | Hexamerin 2 beta | 5.04 | 3.85 | 1.71 |
| Chemosensory | AAEL002626 | OBP56e | 1.20 | 1.75 | 1.60 |

### Significantly differentiated genomic regions and genes

Fst analysis revealed 1,210 genes significantly differentiated between the different groups (File S4), seven of which were also overexpressed in all three comparisons, including the two CYP450s; *CYP6AG7* and *CYP9J26*. Insecticide resistance-associated genes with significant Fst and overexpression within each comparison can be seen in Fig S5. Owing to their more specific link to a resistant phenotype, the CON vs FEN comparison is of particular interest and detected six genes with significant Fst values on chromosomes 2 and 3 which were also overexpressed (Fig 4). This included *GSTE6*, *CYP4AR2*, *CYP4H28*, and AAEL023509 which were overexpressed only in the CON vs FEN comparison, and *CYP6AG7* and *CYP9J26* which were overexpressed in all comparisons. Alongside this, a nicotinic acetylcholine receptor subunit (AAEL004935) was found to have significant Fst in this comparison.

**Fig 4.**
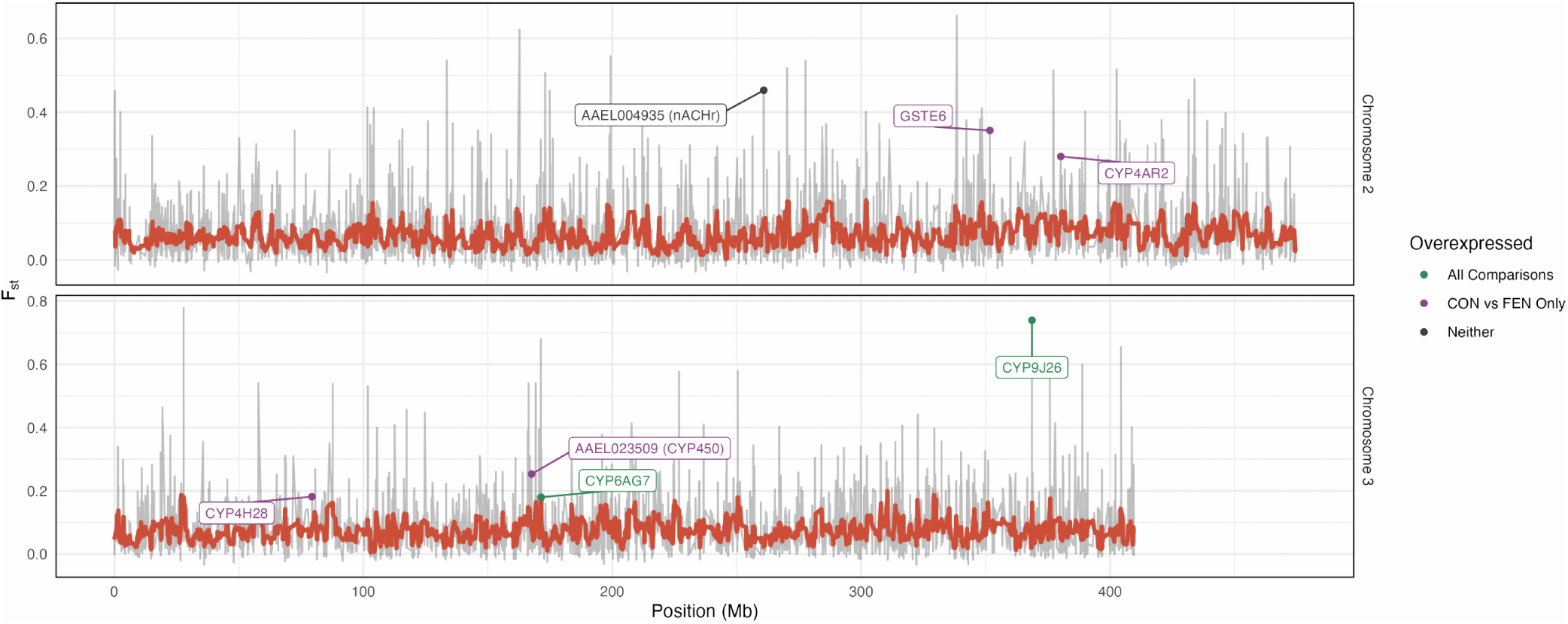
Per gene Fst for CON vs FEN. The grey line shows each per-gene Fst value, and the red line shows the rolling average over every 10 genes. Labelled points indicate genes with significant Fst values two ranges above the mode in gene families associated with resistance, and are coloured according to whether they were significantly overexpressed either in the CON vs FEN comparison, or across all comparisons. Only chromosomes 2 and 3 are shown due to no genes of interest on chromosome 1.

### Identification of target site mutations

Four previously identified non-synonymous *VGSC* (AAEL023266) mutations were detected in the Angolan population (Fig 5A). V410L (C to A at 3:315939224), V1016I (C to T at 3:315983763), and F1534C (A to C at 3:315939224) were all fixed, along with the less well-known mutation S723T (A to T at 3:316014588). Five mutations, previously unreported in *Ae. aegypti*, were found in different *VGSC* domains (Fig 5B) at variable frequencies. K427R (T to C at 3:316080670) was detected at high frequencies in the Angolan samples, whilst also present in the susceptible lab populations so, at least alone, this mutation is unlikely to be resistance-associated. V1365G (A to C at 3:315948012)\ was detected at high frequency in one of the CON sample pools, although only two sample pools contained data for this position. S749I (C to A at 3:316014446) was observed at low frequency in one ROC and CON sample pool each, whilst L879S (AG to GA at 3:316000485) and L963P (A to G at 3:315984154) were identified in the CON and FEN samples, respectively.

**Fig 5.**
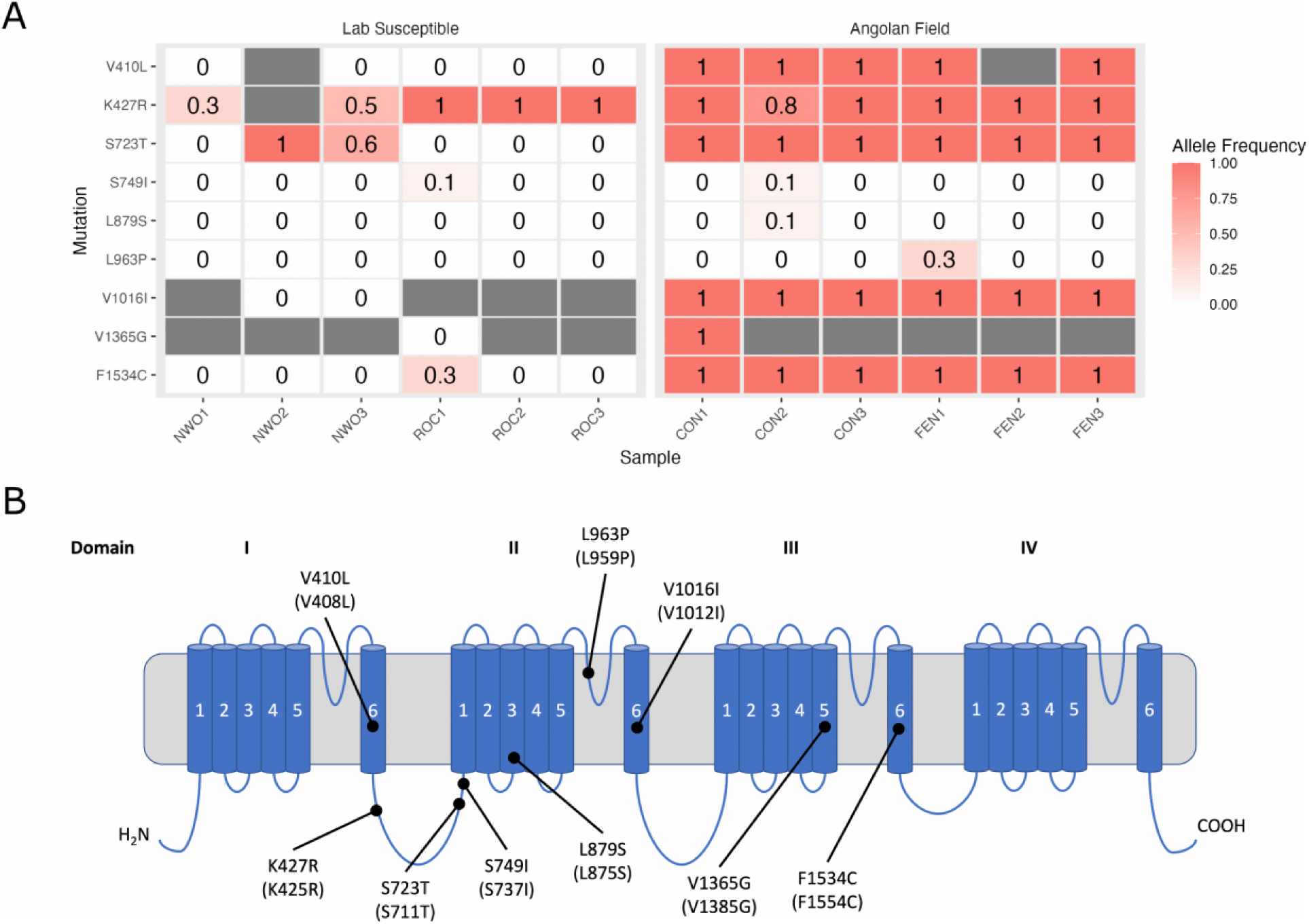
A) Allele frequency of various *VGSC* SNPs called across all samples in each sample pool for each group. Blank cells indicate no sequenced reads at that locus. B) Schematic showing the approximate location of each called SNP observed in the Angolan populations, within the VGSC protein. Amino acid positions of mutations are numbered based on M. domestica *VGSC* (VectorBase ID: MDOA002080-RB). The numbers of the corresponding positions in *Ae. aegypti VGSC* (VectorBase ID: AAEL023266-RL) are shown in brackets

*ACE-1* (AAEL000511) and *RDL* (AAEL008354) were also investigated due to their known role in resistance in multiple mosquito species. Eight mutations were detected in *ACE-1*, with four absent in the susceptible colonies and at low frequency in the Angolan samples (File S5) including L696V (G to C at 3:161487451) at 0.1 in FEN, D80N (CTGAA to TTGAG at 3:161510788) at 0.07 in CON, L47V (T to G at 3:161694909) at 0.07 in CON and 0.03 in FEN, and N152D (A to G at 3:161695287) at 0.03 in FEN. Meanwhile, 14 mutations were identified in *RDL*, with three mutations being absent in the susceptible colonies (File S5), comprising G64V (G to T at 2:41742575) and A296S (G to T at 2:41847790) at low frequencies of 0.07 and 0.03 in the CON and FEN samples, respectively, alongside M97I (G to T at 2:41755160), which was detected only in the FEN samples at high frequency (0.7) with no corresponding data in susceptible sample pools.

The six genes in families associated with resistance which exhibited both significant Fst and significant overexpression in the CON vs FEN comparison, revealed several SNPs of interest, following filtering for absence in the susceptible (NWO and ROC) groups (Table 2 and File S5). This included two SNPs in *GSTE6*, 14 in *CYP4AR2*, five in *CYP4H28*, three in AAEL023509 (CYP450), and nine in *CYP6AG7*. AAEL004935 (*nAChRb2*) also appeared to have two SNPs of interest (K214N and V393L). *CYP9J26* contained one non-synonymous SNP, but this mutation was not absent in the susceptible colonies.

**Table 2.** Allele frequencies for the identified SNPs in the CON and FEN groups within genes of interest.

| Gene ID | Gene Description | SNP Location | Nucleotide Mutation | Amino Acid Substitution | CON | FEN |
| --- | --- | --- | --- | --- | --- | --- |
| AAEL004935 | nAChRb2 | 2:260851442 | G>T | K214N | 0.17 | 0 |
|  |  | 2:260852154 | G>T | V393L | 0.27 | 0 |
| AAEL007946 | GSTE6 | 2:351707645 | CAGGTCC>GAGCTCT | D171E | 0 | 0.15 |
|  |  | 2:351708094 | T>C | T45A | 0.03 | 0 |
| AAEL010154 | CYP4AR2 | 2:380179404 | T>G | L29R | 0.5 | 0.67 |
|  |  | 2:380179566 | C>G | S83C | 0 | 0.77 |
|  |  | 2:380179758 | T>A | V147E | 1 | 0.67 |
|  |  | 2:380179796 | T>C | S160P | 0.35 | 0 |
|  |  | 2:380179806 | AGGTA>TGGTG | E163V | 1 | 0 |
|  |  | 2:380180100 | G>A | G261E | 0.13 | 0 |
|  |  | 2:380180124 | C>G | S269C | 0 | 0.17 |
|  |  | 2:380180133 | T>C | L272S | 0 | 0.57 |
|  |  | 2:380180180 | G>A | D288N | 0 | 0.23 |
|  |  | 2:380180575 | T>C | S378P | 0.07 | 0 |
|  |  | 2:380180637 | CACTG>AACCA | V400I | 0 | 0.33 |
|  |  | 2:380180822 | C>T | T460I | 0 | 0.03 |
|  |  | 2:380180928 | AAG>GAA | R496K | 0.15 | 0.63 |
|  |  | 2:380180930 | G>C | R496T | 0 | 0.07 |
| AAEL003380 | CYP4H28 | 3:79340421 | G>A | L507F | 0.5 | 0.07 |
|  |  | 3:79340461 | A>C | I493M | 0.5 | 0.07 |
|  |  | 3:79340514 | A>C | L476V | 1 | 0.03 |
|  |  | 3:79340634 | G>A | P436S | 0 | 0.03 |
|  |  | 3:79353142 | A>C | I103S | 0 | 0.33 |
| AAEL023509 | CYP450 | 3:167617984 | A>T | K18N | 0.5 | 0.33 |
|  |  | 3:167627879 | C>G | H141Q | 0 | 0.23 |
|  |  | 3:167627943 | G>A | V163I | 0 | 0.33 |
| AAEL006989 | CYP6AG7 | 3:171418122 | G>T | A71S | 0 | 0.03 |
|  |  | 3:171418176 | G>A | A89T | 0.07 | 0.1 |
|  |  | 3:171418192 | T>C | V94A | 0.13 | 0 |
|  |  | 3:171418551 | CGC>TGG | A194G | 0.07 | 0 |
|  |  | 3:171418575 | A>T | K201N | 0.03 | 0 |
|  |  | 3:171418760 | A>G | E263G | 0.13 | 0.23 |
|  |  | 3:171419086 | C>T | P372S | 0.13 | 0 |
|  |  | 3:171419113 | CGAAG>AGAAA | S382N | 0 | 0.03 |
|  |  | 3:171419272 | C>T | P434S | 0.07 | 0 |

### Metabolism assays and kinetics studies

Owing to its consistent overexpression across all comparisons, significant Fst, and lack of prior functional characterisation, CYP6AG7 was selected for functional validation. Its activity was compared to *An. gambiae* CYP9J5, a CYP450 known to metabolise pyrethroids, pyriproxyfen, and chlorfenapyr [42,109]. Both CYP450s were recombinantly expressed, and their CO-reduced spectra were examined for peaks at 420 nm. The absence of such peaks in the final membranes suggested low levels of inactive enzyme (Fig S6). Metabolism assays were first conducted for CYP9J5 and CYP6AG7 against fenitrothion to assess insecticide depletion. Additional assays were performed with deltamethrin to evaluate the activity of CYP6AG7 against another class of insecticide to which the Angolan population exhibited resistance (Fig S1).

Metabolism assays with deltamethrin enzyme showed moderate depletion when incubated with CYP6AG7 (14.64%), and greater depletion with CYP9J5 (34.33%) (Fig 6A). Metabolism assays with fenitrothion produced a small depletion with CYP6AG7 (7.66%), and a substantial depletion (26.78%) with CYP9J5 (Fig 6B). In all fenitrothion metabolism assays, a metabolite with retention time of 3.7 minutes was detected and designated M1 (Fig S7), which was hypothesised to be the fenitrothion-oxon. Consistent with this, the peak area of M1 increased in proportion to fenitrothion depletion (Fig 6B and Fig 6C). Identification of M1 as the fenitrothion-oxon was supported by a standard curve generated using authentic fenitrothion-oxon (Fig S8), which showed a linear relationship between concentration and peak area at the same retention time (R2 = 1.00).

**Fig 6.**
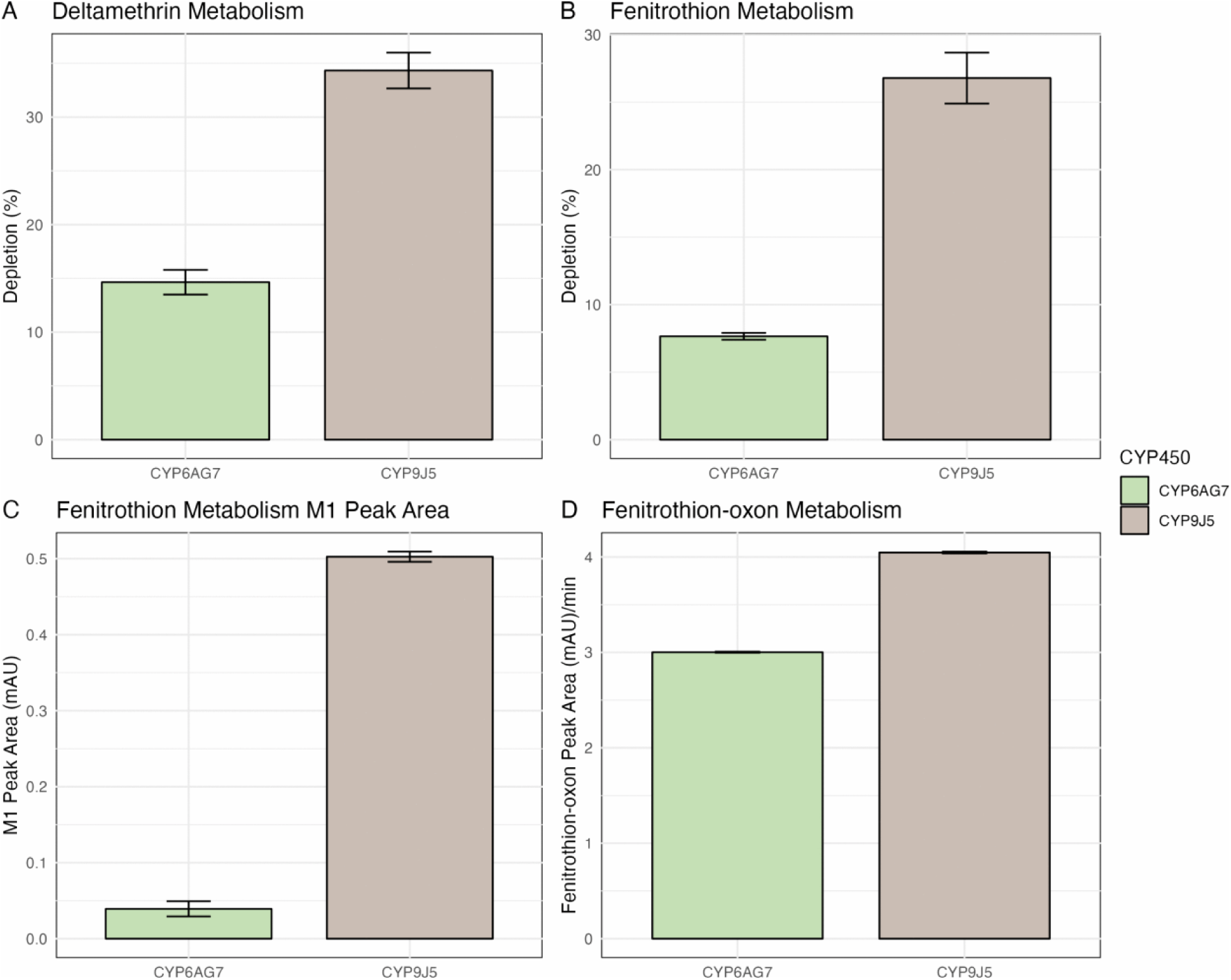
Metabolism of deltamethrin and fenitrothion by recombinant P450 enzymes CYP6AG7 and CYP9J5 as a positive control. A) Percentage depletion of 10µM deltamethrin after incubation with 0.1µM of CYP6AG7 or CYP9J5. B) Percentage depletion of 10µM fenitrothion after incubation with 0.1µM CYP6AG7 and CYP9J5. C) Peak area of M1 metabolite following fenitrothion metabolism in reactions corresponding to panel B, indicating formation of the primary metabolite. D) Peak area of fenitrothion-oxon after incubation with either 1µM CYP6AG7 or 0.1µM CYP9J5. Bars show mean ± standard deviation.

Metabolism assays using fenitrothion-oxon as the substrate showed no difference in peak area between +NADP+ and -NADP+ conditions for CYP6AG7 or CYP9J5, indicating that they do not further metabolise the fenitrothion-oxon (Fig 6D). Time-course assays for both CYP450s showed a linear production of the fenitrothion-oxon over time (Fig S9A and Fig S9B). Kinetic analysis demonstrated increasing fenitrothion-oxon formation with increasing fenitrothion concentration for both CYP450s, before reaching saturation (Fig S9C and Fig S9D). CYP6AG7 exhibited a fenitrothion-oxon turnover rate (*Kc*at) of 0.08 min-1, while CYP9J5 showed a substantially higher rate of *K*cat at 0.65 min-1; approximately eight times greater turnover. In contrast, the Michaelis constants were similar for both CYP450s (*K*m = 31.90µM for CYP6AG7 and *K*m = 30.55µM for CYP9J5), indicating similar fenitrothion binding affinities. Overall, these results demonstrate that although both CYP450s can bioactivate fenitrothion to its oxon form, CYP9J5 shows markedly greater turnover and catalytic efficiency than CYP6AG7, and neither enzyme appears to further detoxify the toxic oxon metabolite.

### Molecular docking

The best docked pose between deltamethrin and CYP6AG7, and deltamethrin and CYP9J5, had predicted binding free energies of −9.429 kcal/mol and −8.483 kcal/mol, respectively. The best scored pose for which the sulphur atom in the fenitrothion was orientated towards the heme group was ranked at 8 (RMSD from best mode of 1.926) for CYP6AG7 with a binding free energy of −5.869 kcal/mol, and at 9 (RMSD from best mode of 2.192) for CYP9J5, with a similar binding free energy of −5.349 kcal/mol. Residues comprising the surrounding of the binding pocket for CYP6AG7 included Phe 105, Phe 122, Asp 297, Thr 301, Ala 362, Phe 364, Met 365, and Leu 483 with bonding shown in Phe 122, Asp 297, and Met 365, and a distance of 3.31Å between the fenitrothion sulphur and heme iron (Fig 7 A and C). For CYP9J5, residues in the surrounding the binding pocket included His 105, Ile 126, Leu 340, Ile 337, Ala 341, Thr 345, Ala 408, Ala 410, Thr 411, alongside bonding shown in Ile 126, Ala 408, and Thr 411, and a distance of 3.61Å between the fenitrothion sulphur and heme iron (Fig 7 B and D).

**Fig 7.**
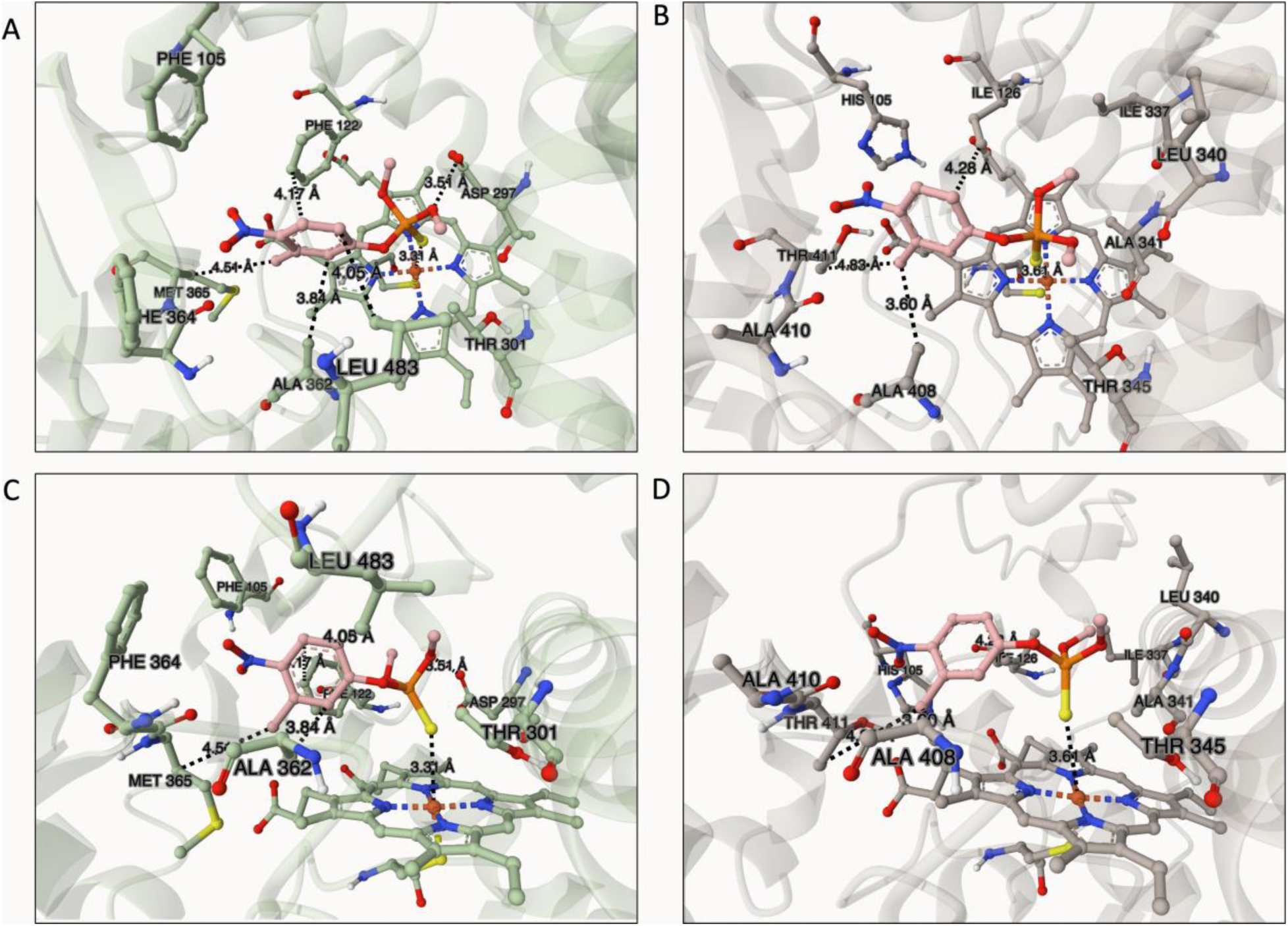
Visualisation following docking of CYP6AG7 (A, top view; C, side view) and CYP9J5 (B, top view; D, side view) with fenitrothion. Amino acid residues within the surrounding binding pocket which have predicted hydrophobic interactions with fenitrothion are shown. Distances labelled show protein-ligand interactions highlighted by PLIP, along with the distance between the fenitrothion sulphur atom and the heme iron.

## Discussion

This work represents a first study of the range of resistance mechanisms present in an African *Ae. aegypti* population; a priority given the current lack of data, and the continent’s rapidly escalating dengue burden and the heightened need for effective vector control [127]. Several of the candidate genes identified in this Angolan *Ae. aegypti* population are from key detoxification families such as CYP450s. Generally, upregulated CYP450s were from the CYP304, CYP325, CYP4, CYP6, and CYP9 subfamilies as is commonly seen in resistant populations of this species [10,12,24]. Many of these mirror those identified outside of Africa (Table 1 and Fig S4), for example, *CYP9J24* and *CYP9J26* in permethrin resistant populations from Thailand and Mexico [40,128], AAEL029064 (*CYP9J27*) in pyrethroid resistant populations from Malaysia [126], and *CYP6BB2* in several pyrethroid resistant wild populations and resistant lab colonies [129–132]. Many further instances of upregulation of these genes in a resistant phenotype can be found in literature [24]. It appears that the genes most commonly implicated in pyrethroid resistance are also associated with fenitrothion resistance in Angola. Several of the lesser-reported CYP325 family genes were overexpressed in the Angolan samples via NWO and ROC vs CON comparisons (Fig S4). Upregulation of genes in this subfamily has been observed in pyrethroid resistant *Ae. aegypti* from Mexico and Malaysia [126,133,134]. Candidate gene (qPCR) studies within Africa on resistant *Ae. aegypti* from Burkina Faso and Senegal have reported upregulation of several of the same CYP450s as seen in the present dataset including *CYP9J26*, *CYP9J28*, *CYP6BB2*, and *CYP6Z8*, compared to susceptible lab strains [71,135]. More recently, a genomic duplication spanning the CYP6 cluster (*CYP6BB2*, AAEL026582, AAEL017061, *CYP6P12,* and *CYP6CC1*) was shown to be involved in deltamethrin resistance [39]. Indeed, almost all of these genes were overexpressed in this study (File S3), but were slightly under the threshold of reporting according to log2 fold change threshold applied across all comparisons.

There are also two novel CYP450s identified here with limited presence in the literature outside of Africa: *CYP304B2* and *CYP6AG7*. *CYP304B2* was noted to have increased copy number in multi-resistant *Ae. aegypti* from French Guiana [136], has recently been shown to exhibit progressively higher expression across *Ae. aegypti* lines from Colombia selected for increasing resistance to lambda-cyhalothrin [137], whilst *CYP6AG7* was seen to be upregulated in a Mexican population from Mérida compared to the NWO strain [138], as well as in deltamethrin resistant *Ae. aegypti* from Thailand [57]. Interestingly, *CYP6AG7* has also been observed to be upregulated in malathion and lambda-cyhalothrin resistant *Ae. aegypti* in Puerto Rico [132], and in a temephos-resistant *Ae. aegypti* laboratory colony originally from Brazil [139]. Generally upregulated CYP450s, either constitutively or following fenitrothion exposure, in this Angolan population were from the CYP304, CYP325, CYP4, CYP6, and CYP9 subfamilies as is commonly seen in resistant populations of this species [10,12,24].

Metabolism assays revealed that the insecticide resistance candidate CYP6AG7 was able to deplete both deltamethrin and fenitrothion to varying extents, although to a lesser extent than the CYP9J5 positive control (Fig 6 and S9). CYP6AG7 also catalysed the turnover of fenitrothion, producing the metabolite M1 identified as the fenitrothion-oxon (Fig 6, S7, and S8). Production was at a much lower rate (∼8 fold) than CYP9J5, although both enzymes displayed similar substrate binding affinities, as measured by comparable *K*m values (Fig S9).

Consistent with the metabolism studies, molecular docking analysis predicted higher binding energies of both P450s for deltamethrin than for fenitrothion. Furthermore, they predicted similar binding energies for fenitrothion, in agreement with the calculated Km values. The approximately eight-fold higher rate of metabolism of fenitrothion by CYP9J5 compared to CYP6AG7 may be explained by difference in active-site architecture rather than substrate binding. Docking analysis revealed that the residues lining the binding pocket differed between the two enzymes. For CYP9J5, predicted non-covalent interactions with fenitrothion were primarily mediated by Ile 126, Ala 408, and Thr 411, whilst CYP6AG7 interactions involved Phe 122, Asp 297, Ala 362, and Met 365 (Fig 7), as identified using PLIP. These differences in interacting residues may produce a more catalytically efficient and metabolically productive orientation of fenitrothion in the CYP9J5 active site, thereby contributing to its higher metabolic activity.

It is unclear, and apparently paradoxical, why *CYP6AG7* would be upregulated in the CON vs FEN comparison given that, according to our experiments, it is metabolising the pro-insecticide fenitrothion into the harmful oxon form which acts upon the acetylcholinesterase receptor leading to death [140]. However, it should be noted that the Angolan population was resistant to several insecticide classes, including pyrethroids (Fig S1), with *CYP6AG7* shown here to also metabolise deltamethrin, and fenitrothion resistant females might also be more strongly pyrethroid resistant. It is thus plausible that upregulation of this gene is favourable to detoxifying other insecticides but it is under the same regulatory control pathway as other fenitrothion resistance conferring enzymes that metabolise the oxon. For example, differential expression and silencing of the transcription factor *Maf-S* in *An. gambiae* (AGAP01045), which binds to antioxidant response elements (ARE), has been shown to modulate expression of several detoxification genes including *CYP6M2* and *GSTD1*, conferring insecticide resistance [54]. The homologous *Maf-S* encoding gene in *Ae. aegypti* (AAEL026468) was found to be overexpressed in the CON vs FEN comparison (File S3: log2 fold change = 0.96, adj p value < 0.00055). Examination of the immediate up and downstream region for *CYP6AG7, CYP6BB2, CYP9J24, CYP9J26*, AAEL029064 (*CYP9J27*), and *CYP304B2* revealed the presence of CnC-Maf-S binding sites (Table S1). The CYP6 family in *D. melanogaster* has also been shown to be under the control of this transcription factor, along with GSTD family genes conferring malathion resistance [141,142], likely through hydrolysis of any produced fenitrothion-oxon. Thus, *GSTD6* which was upregulated in the CON vs FEN comparison (File S3) was also checked and found to possess such binding sites (Table S1). Furthermore, the Km value for *CYP6AG7* against fenitrothion is relatively weak compared to previous metabolism studies [40], indicating that *CYP6AG7* might be an auxiliary metaboliser of fenitrothion, acting on other compounds in the fenitrothion exposed survivors but, due to the promiscuous nature of CYP450s, fenitrothion is able to weakly bind and produce the oxon.

Whilst GSTs and COEs were individually upregulated between comparisons (Fig 3), there were no genes in these families with concurrent upregulation across all comparisons (Fig 3 and File S3). Conversely, an ABC transporter (AAEL005043) and a UGT (AAEL003098) were upregulated in all comparisons. AAEL005043 has not been previously associated with adulticide detoxification, but has been seen to be upregulated in larvae guts and suggested to be involved in resistance to *Bacillus thuringiensis* Cry toxins [143]. Upregulation of AAEL003098 in relation to insecticide resistance has yet to be reported. Differential expression of several short-chain dehydrogenases/reductases (SDRs) and haem peroxidases (HPXs) was also observed (File S3), but with no concurrent upregulation across all comparisons. A number of lesser-known enzyme families potentially associated with resistance were identified as upregulated in this study, including hexamerins, HSPs, and OBPs (Fig 3 and File S3). The hexamerin AAEL011169, which was upregulated in all comparisons, was also seen to be upregulated in resistant *Ae. aegypti* from Madeira Island [144]. Several OBPs, a family associated with resistance through insecticide sequestration [31], were among the most upregulated genes in the CON and FEN groups, including *OBP56e* which was upregulated across all comparisons and has not been reported in the context of resistance before. Overall, whilst there is some fluctuation in the less upregulated genes, it appears that the most strongly upregulated detoxification genes, and gene families, are often consistent in insecticide resistant *Ae. aegypti* worldwide.

Many cuticular-associated genes were upregulated across the NWO vs CON and ROC vs CON comparisons, such as those involved in fatty acid synthesis or cuticle proteins (Fig 3 and File S3), suggesting that cuticular resistance may be present in Angolan *Ae. aegypti*, as in implicated permethrin resistant *Ae. aegypti* from Singapore and in pyrethroid selected lab strains from Colombia [26,137]. Previous work by Grigoraki *et al.* [145] delineated the genes involved in hydrocarbon biosynthesis pathway leading to cuticle formation in *An. gambiae* using oenocytes. Following assessment of expression of the homologous genes in *Ae. aegypti* (File S6), we observed intense upregulation in all but 4 genes out of the 28 examined in the NWO vs CON and ROC vs CON comparisons (Fig S10). This includes *CYP4G36*, the orthologue of *CYP4G16* which is known to promote cuticular resistance through hydrocarbon deposition [29]. Furthermore, *CYP301A1*, which was upregulated in the NWO and ROC vs CON comparisons, and AAEL023509 (putative *CYP314A1-1*), which was upregulated in the CON vs FEN comparison and exhibited significant Fst (File S3), have both been shown to be involved in cuticle formation, as well as involved in regulating growth hormones and eclosion, potentially relating to cuticle development [146,147]. However, whether overexpression of these cuticular-associated genes is related to environmental factors such an adaptation to desiccation [29,148,149] in comparison the laboratory strains, or whether they do indeed confer a resistant phenotype is unclear. Indeed, the strong downregulation of many of these same genes in the CON vs FEN comparison complicates inference, but a similar expression pattern in these genes has been seen in organophosphate and pyrethroid resistant *Ae. aegypti* from Puerto Rico [132]. Further investigation is required to validate the role of these genes in African *Ae. aegypti*. Of the many insect cuticle proteins seen to be upregulated here, enhanced expression of AAEL021930, AAEL0256608, and AAEL011444 has been associated with increasing permethrin resistance in a selected line from Colombia [137], whilst the remaining cuticle protein-encoding genes remain largely undescribed in relation to insecticide resistance, but may be involved in cuticular modification through chitin binding, helping to form the organised structure [29,150,151]. ABC transporters have also been implicated in cuticular deposition and assembly and tissue localisation [152–154], perhaps providing a mechanistic explanation for upregulation of certain ABC transporter encoding genes (Fig 3 and File S3).

Investigation of target site variants revealed the presence of several mutations in *ACE-1* which were absent in the susceptible colonies sequenced. Indeed, all *ACE-1* mutations reported have not been observed before and may represent candidates for OP resistance; although, considering their low frequency, such markers would require further investigation. Interestingly, C699S, which has previously been found in West and Central Africa, and the Americas [155], was seen in the ROC samples (File S5), and may indicate that the mutation is not involved in resistance. Of the reported *RDL* mutations, only A296S has been seen before, being present in Africa, the Americas, and Asia [155–157], and likely represents historic use of organochlorines.

In *VGSC* (Fig. 5), the known variants F1534C [158–161], V1016I [72,74,162,163], and V410L [74,164–167], which are associated with pyrethroid resistance, were all found to be fixed in the Angolan populations. The V1016G mutation often seen in Asia was not detected, in line with previous findings in West and Central Africa [13,24,84]. However, this mutation has recently been observed in Senegal and Benin [168–170], representing the first reports of this mutation in Africa, and its presence should still be monitored in future surveillance efforts. S723T, previously reported across the Americas and in West Africa [84,134,171,172], was again detected here at high frequency. Five additional VGSC mutations (K427R, S749I, L879S, L963P, and V1365G) were also identified, with K427R previously detected in *Ae. albopictus* from Italy and Vietnam [173], but their patterns across sample pools limit any conclusions about relevance to phenotypic resistance. Interestingly, whilst individual haplotypes were not determined, given that many detected *VGSC* mutations were fixed in the Angolan samples means they existed as a haplotype in the individuals sequenced. Such combination mutants have been shown in *Ae. aegypti* to increase phenotypic resistance in a synergistic manner, for example 989P/1016G/1534C [68] and as V410L/V1016I/F1534C [67,165].

SNP analysis of upregulated genes with significant Fst revealed several variants in metabolic resistance genes (Table 2). Interestingly, in a previous study examining variants in permethrin-resistant *Ae. aegypti,* mutations in *CYP4AR2* were shown to be among the most highly associated with resistance, across all detoxification genes [134], although none of those SNPs were identified here. This gene was also upregulated in resistant *Ae. aegypti* from French Guyana [38]. In the same variant association study, one mutation in *CYP6AG7 was* noted [134], but was not among those found in this study. No resistance-associated mutations have previously been reported in *CYP4H28*, but it was overexpressed in temephos and deltamethrin resistant populations from Thailand [49,57]. The presence of these non-synonymous mutations may enhance the binding affinity of upregulated CYP450s against current public health insecticides, leading to their selection, although further validation to elucidate their impact is required. The lack of non-synonymous mutations in *CYP9J26*, despite exhibiting significant Fst, suggests this differentiation is driven by other SNPs. The nicotinic acetylcholine receptor subunit AAEL004935 demonstrating significant Fst, is notable because its *An. gambiae* orthologue (AGAP010057), has previously shown a strong Fst signal associated with pirimiphos-methyl resistance [174]. Although neonicotinoid resistance through mutations in these subunits is well characterised in agricultural pests [175–180], its molecular basis in mosquitoes, and its possible role in organophosphate resistance, remains poorly understood, and warrants further investigation.

## Conclusion

This work has consolidated the association of known key gene families with insecticide resistance in *Ae. aegypti* in this Angolan population and identified a significant number of resistance gene candidates and mutations using a genome- and transcriptome-wide approach. Also highlighted is the presence of several auxiliary mechanisms of resistance seen in this study, such as upregulation of hexamerins, odorant binding proteins, and cuticular-associated genes, that demonstrate the complex multi-layered resistance mechanisms evolutionarily employed by this mosquito population. We provide a list of candidate genes and mutations for cross-reference for future studies of African *Ae. aegypti* populations. A greater emphasis on ‘omics-based research should be employed, to disentangle resistance mechanisms present in the highly genetically and bionomically variable *Ae. aegypti* populations of Africa, where surveillance is currently poor, to inform rational insecticide choices and facilitate molecular surveillance.

## Supplementary Figures and Tables

**Fig S1.**
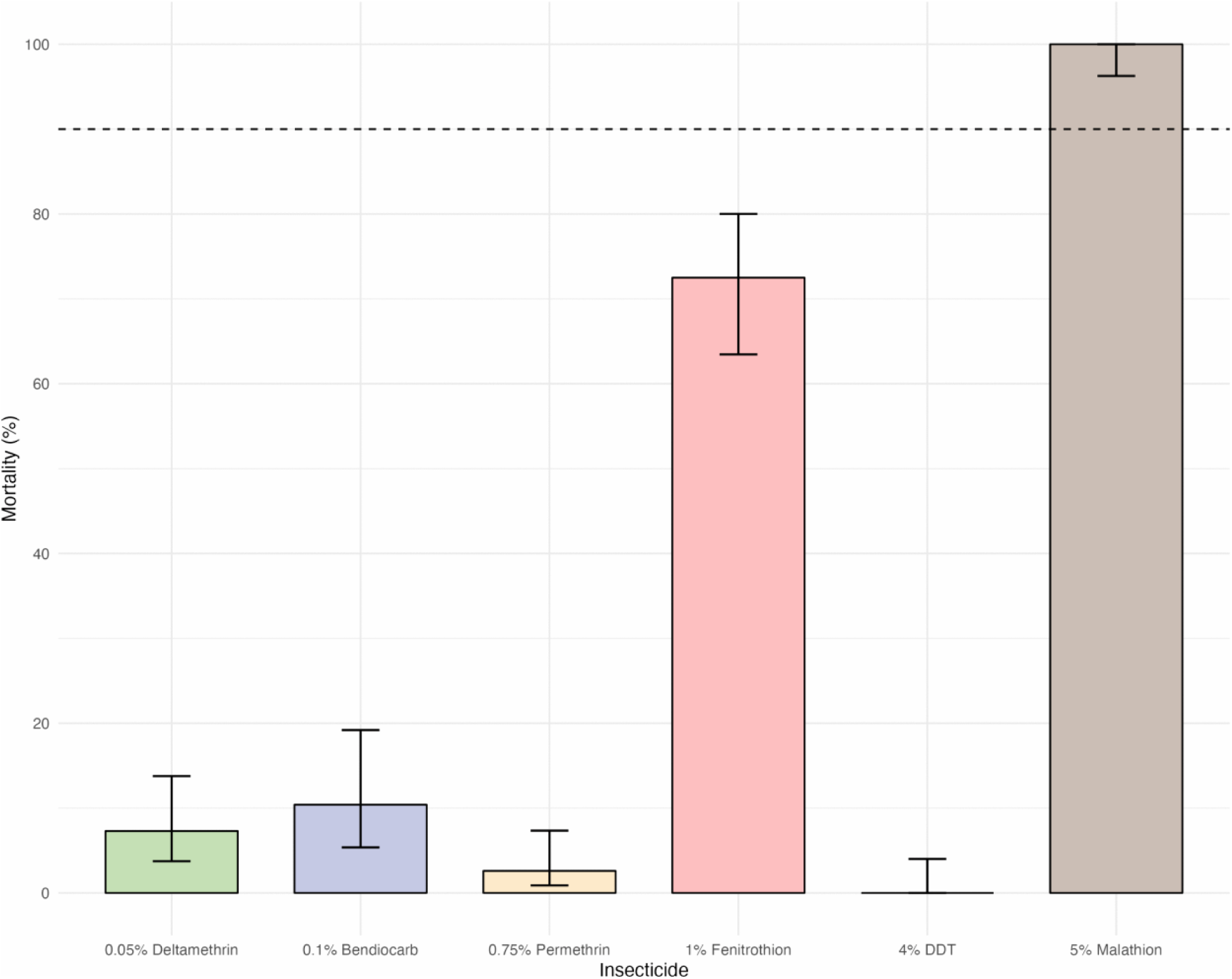
Bioassay results for the five insecticides tested at given concentrations. Error bars show 95% binomial confidence intervals. Dashed line indicates threshold for confirmed resistance (mortality < 90%) according to WHO guidelines (WHO, 2013).

**Fig S2.**
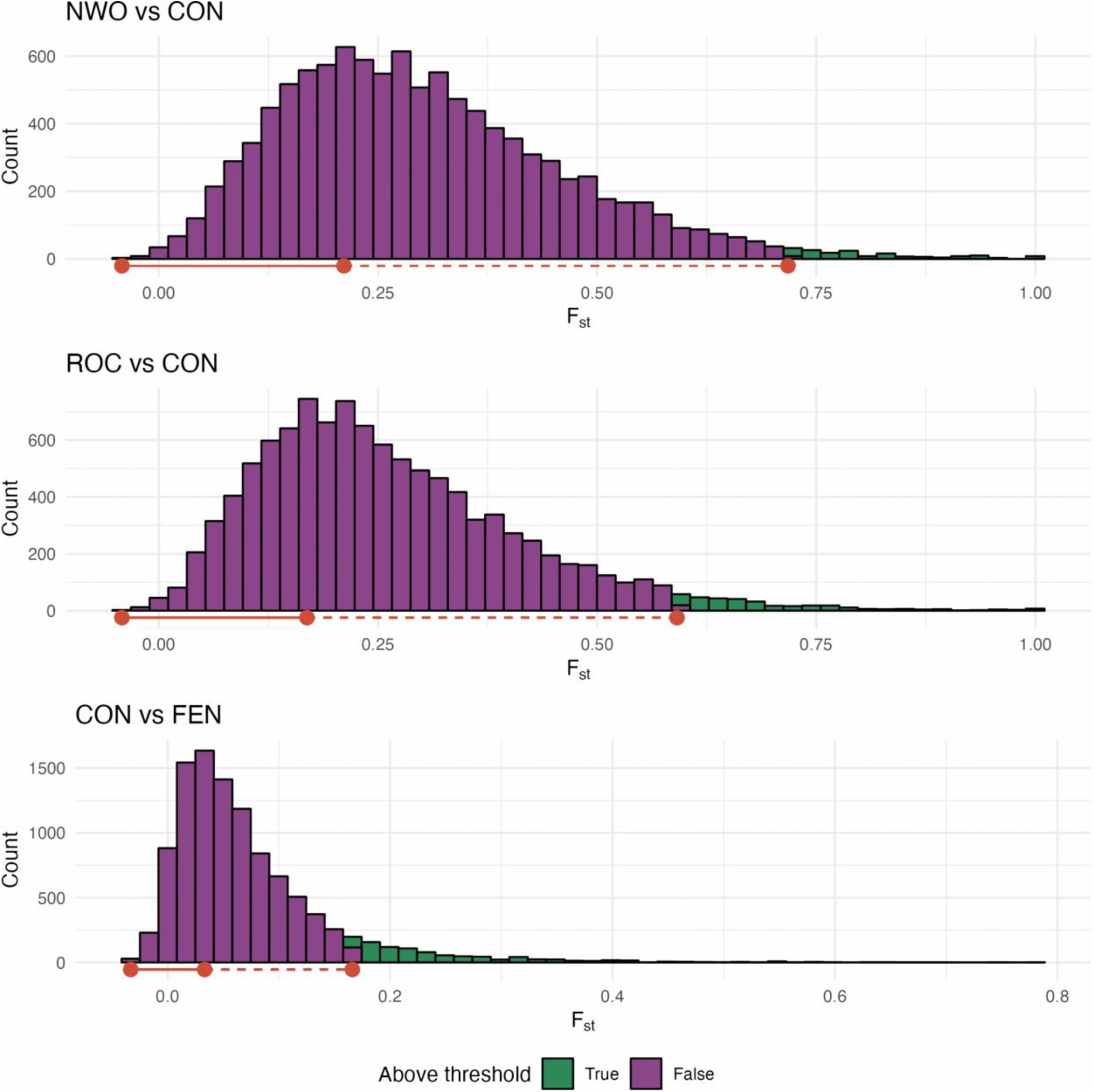
Distribution of per-gene Fst with a bin size of 50. Smallest binned Fst, modal Fst, and Fst two ranges of this difference above the modal Fst are shown in red. Genes for which their Fst was above this threshold are coloured as green bars.

**Fig S3.**
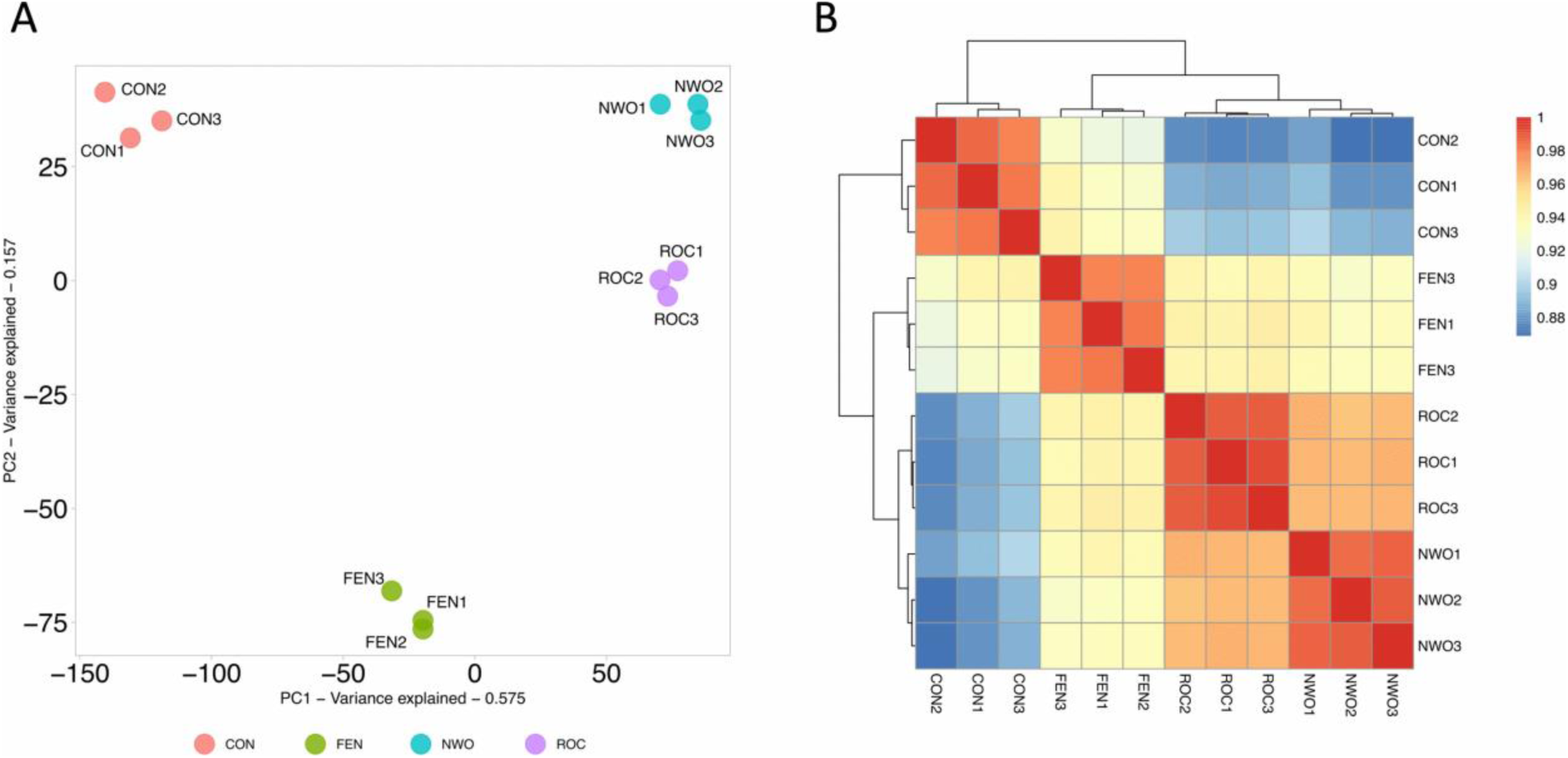
Principal component analysis (A) and Pearson’s correlation heatmap (B) for each sample pool across the four groups according to normalised read count data. Dendrograms show hierarchical clustering of Pearson’s correlations.

**Fig S4.**
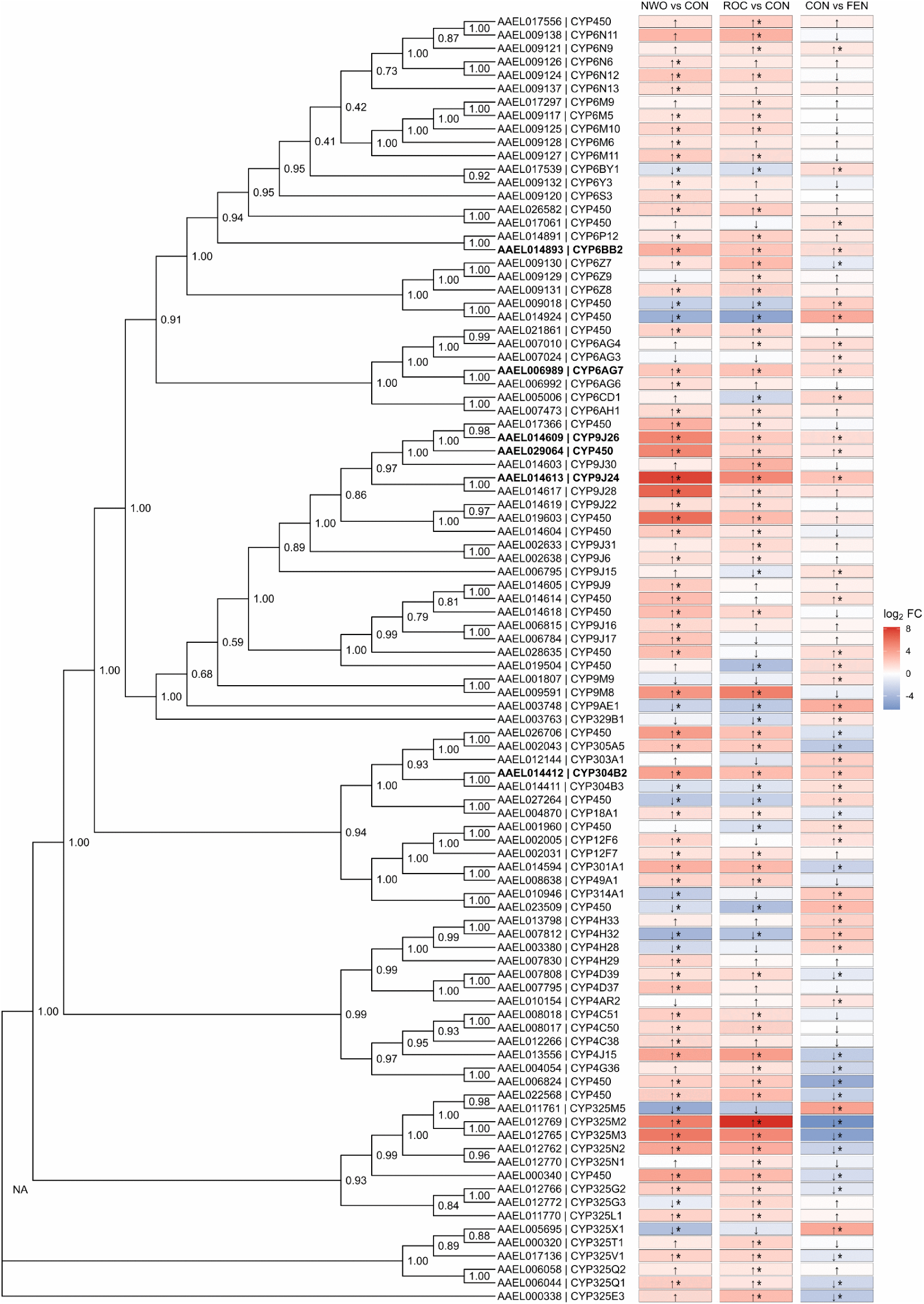
Phylogenetic analysis of all CYP450s which were upregulated across any of the three comparisons. Arrows indicate direction of expression, and asterisks indicate significance. Bold genes were upregulated in all comparisons. Phylogeny constructed using maximum likelihood with bootstrap (1000 replicates). Numbers indicate the frequency of replicates which agree with that node

**Fig S5.**
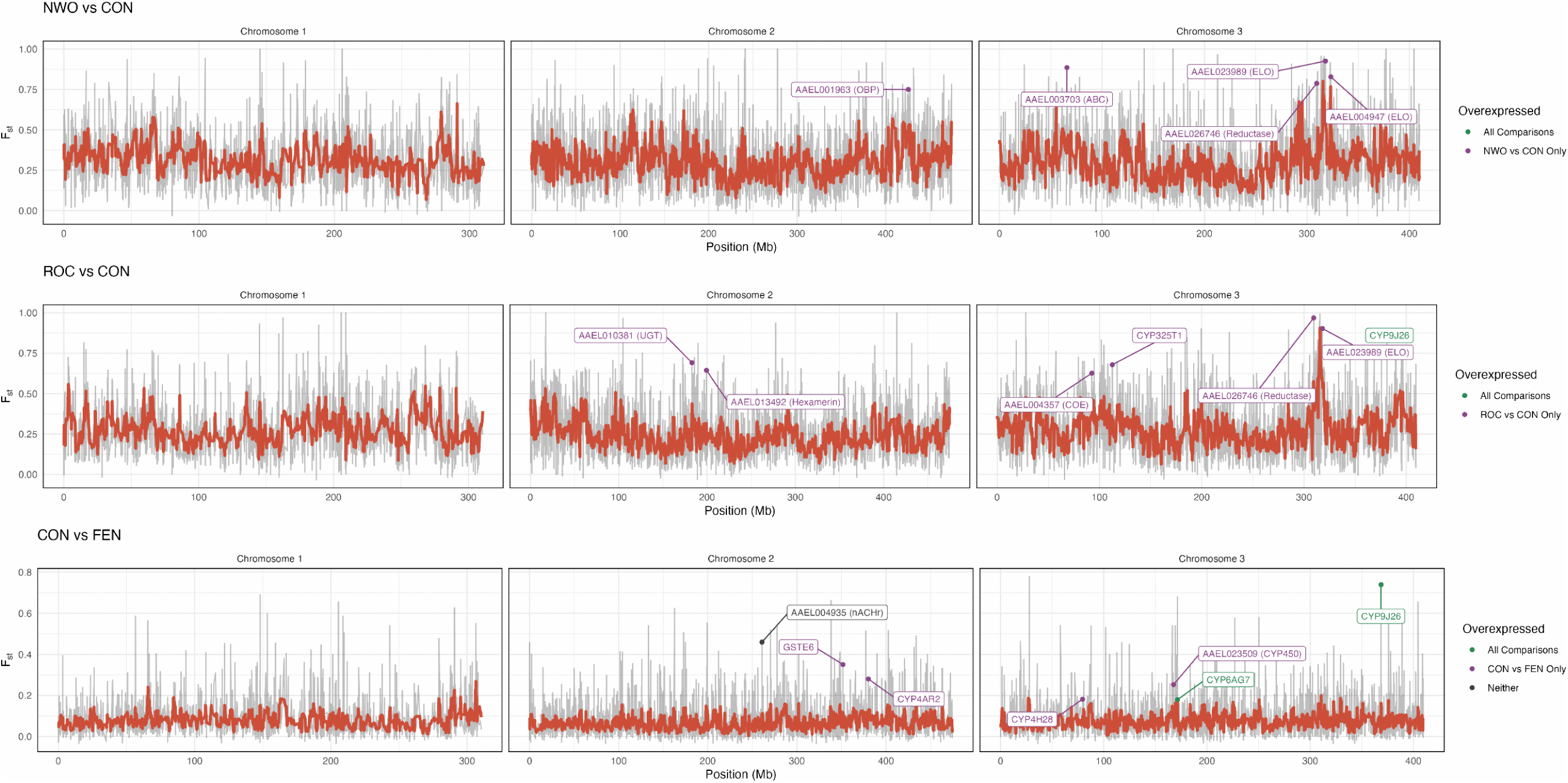
Per gene Fst for all comparisons. The grey line shows each per-gene Fst value, and the red line shows the rolling average over every 10 genes. Labelled points indicate genes with significant Fst values two ranges above the mode in gene families associated with resistance, and are coloured according to whether they were significantly overexpressed either in the comparison shown in the panel, or across all comparisons.

**Fig S6.**
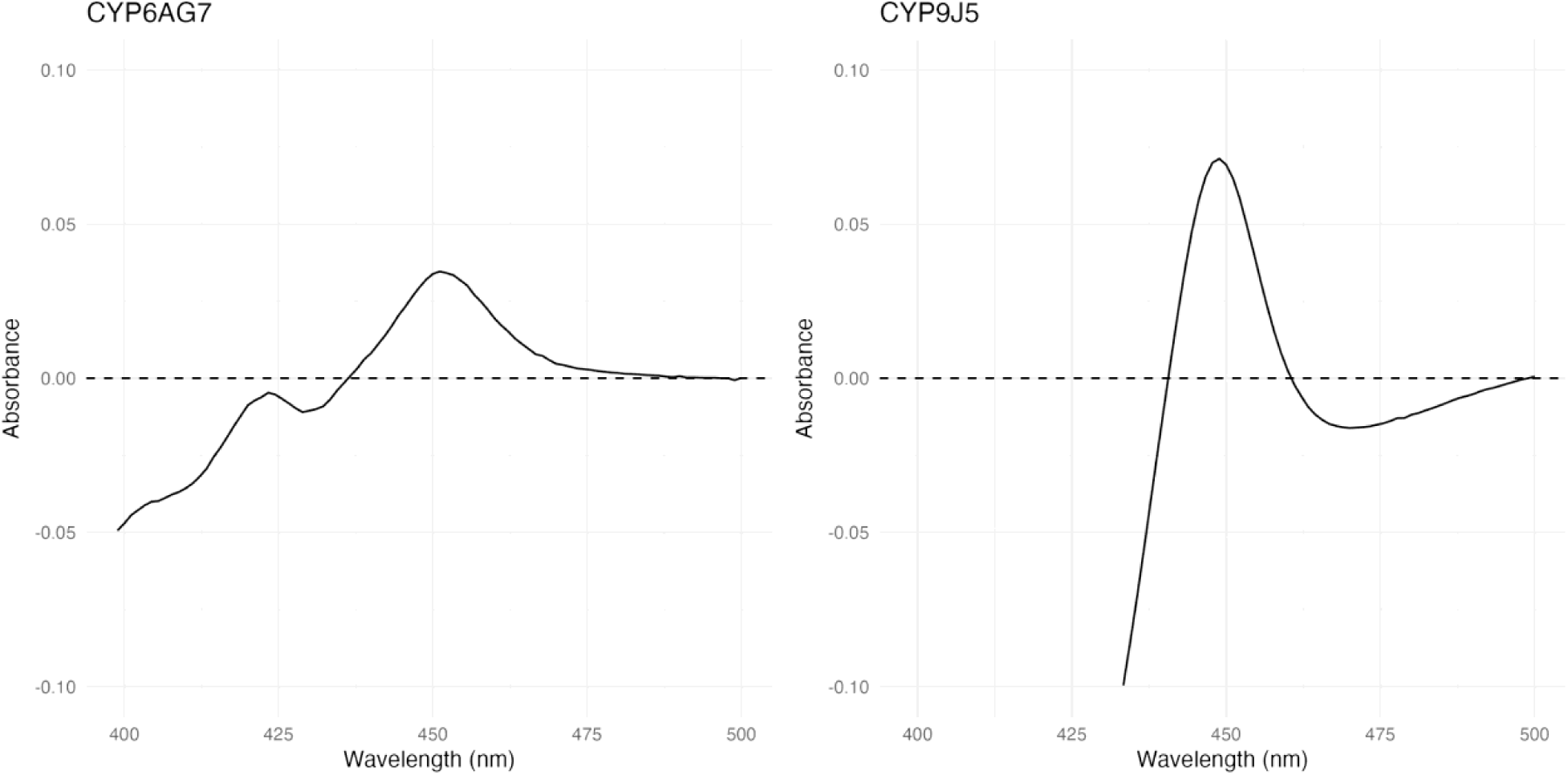
Carbon monoxide difference spectra of final bacterial membranes expressing the *Ae. aegypti* resistance candidate CYP6AG7 and the positive control *An. gambiae* CYP9J5.

**Fig S7.**
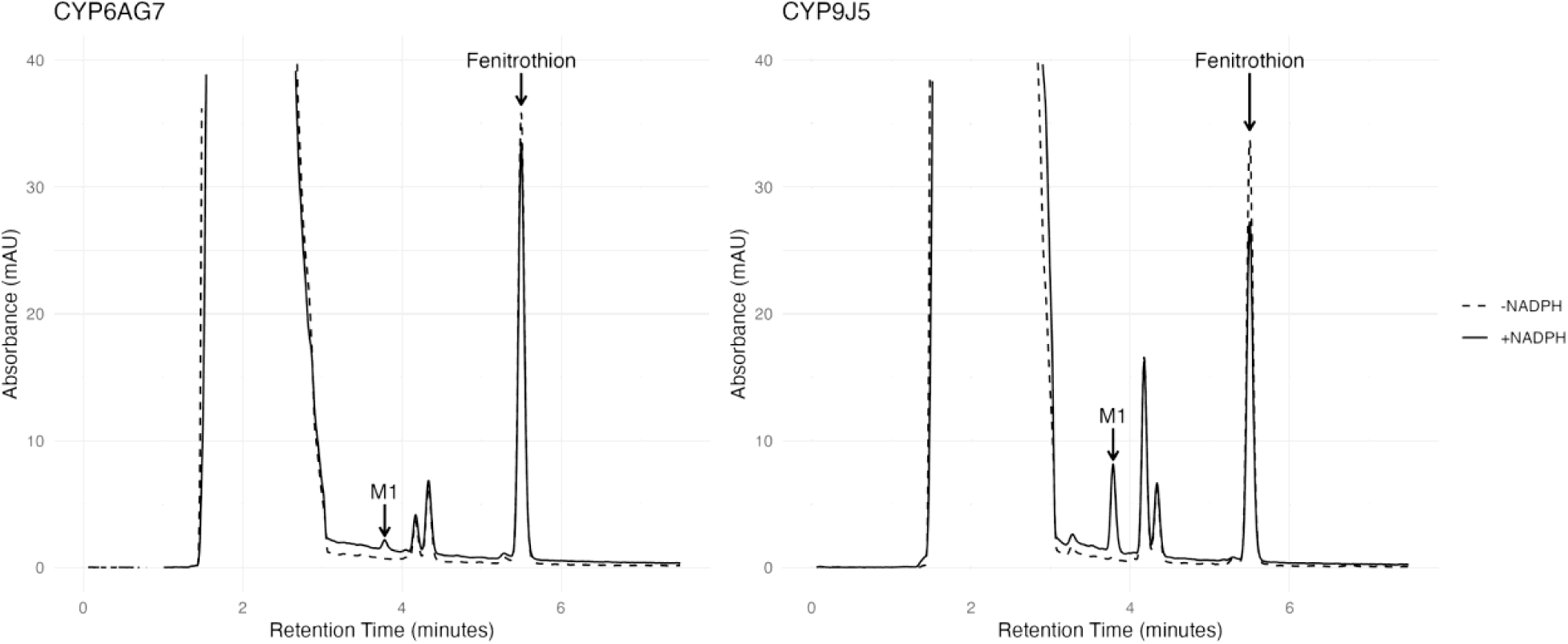
Representative chromatogram for reactions between CYP6AG7 and CYP9J5 with fenitrothion. Labelled are the peaks for fenitrothion and the metabolite M1 (fenitrothion-oxon).

**Fig S8.**
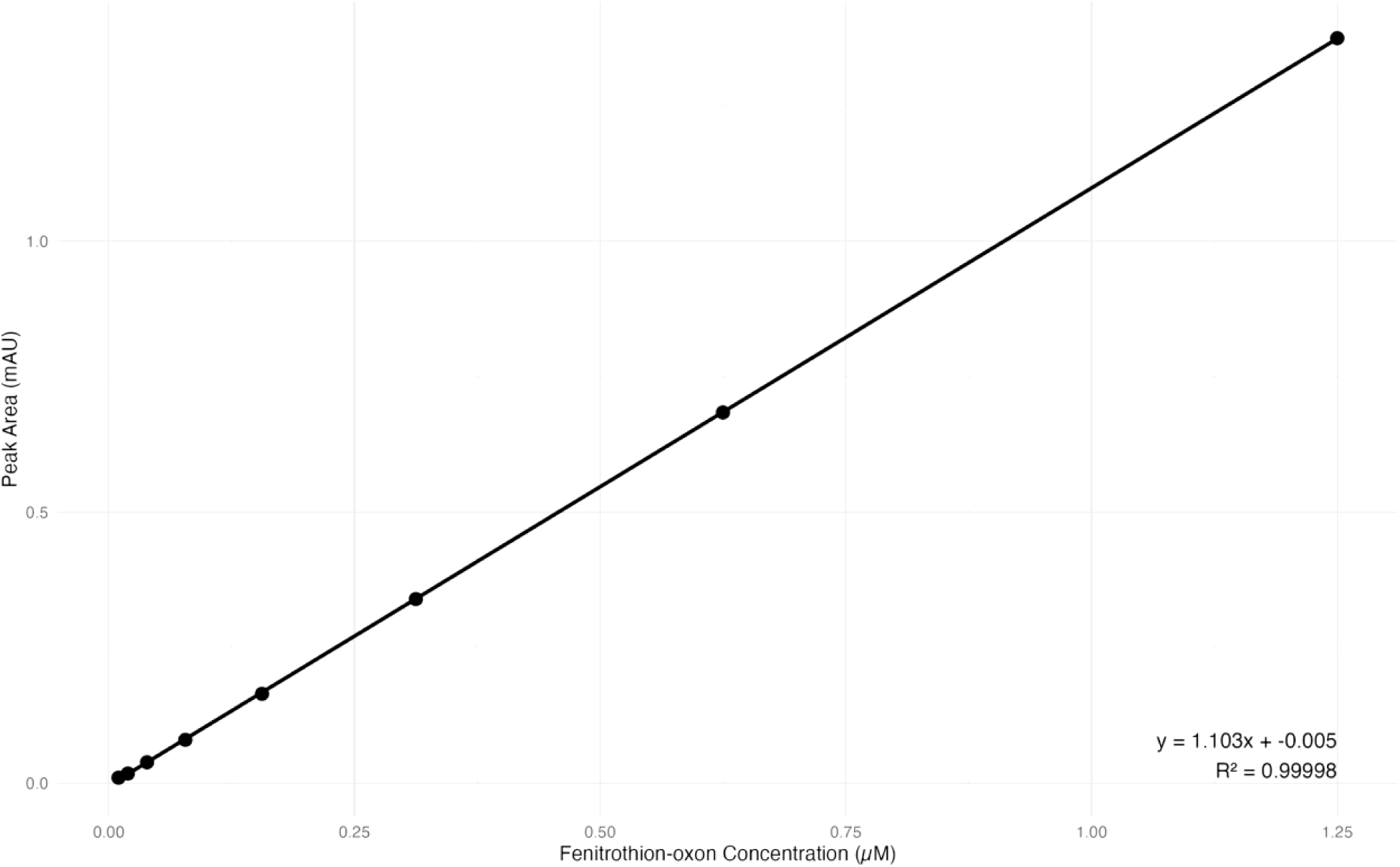
Standard curve of increasing concentrations of fenitrothion-oxon coinciding with increasing peak area with the same retention time as observed metabolite M1.

**Fig S9.**
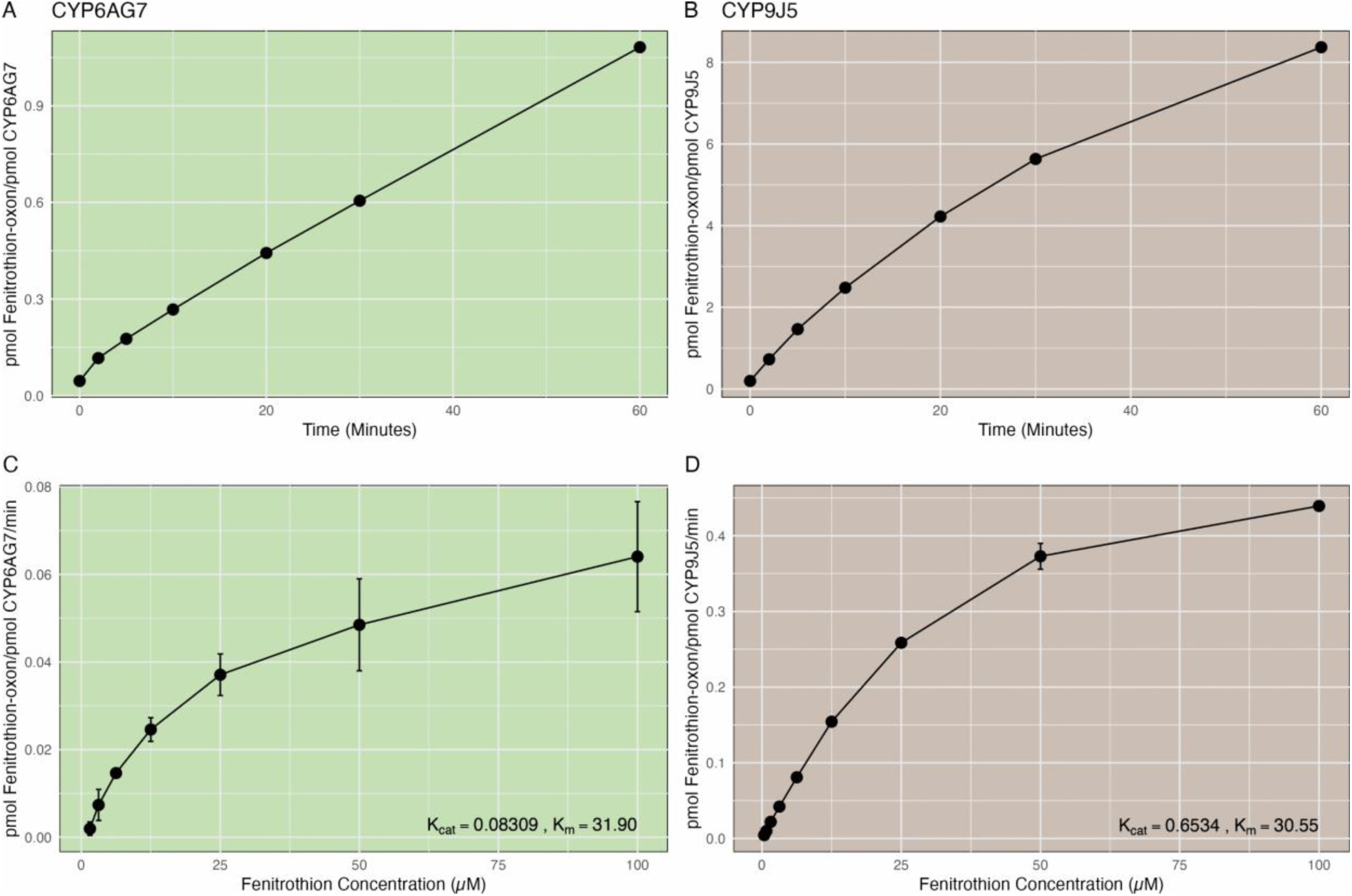
Time course and kinetics analysis of fenitrothion metabolism by CYP6AG7 and CYP9J5 as a positive control. Time course of fenitrothion-oxon formation by 0.2µM CYP6AG7 (A) and 0.1µM CYP9J5 (B) in the presence of 100 µM fenitrothion and NADPH. Michaelis-Menten kinetics of fenitrothion metabolism by 0.2µM CYP6AG7 (C) and 0.1µM CYP9J5 (D), showing rate of oxon formation (pmol/min/pmol CYP450) over a range of fenitrothion concentrations (0.39µM to 100 µM). Bars show mean ± standard deviation. Calculated kinetic parameters, binding affinity (Km) and substrate turnover (Kcat) are shown on each panel.

**Fig S10.**
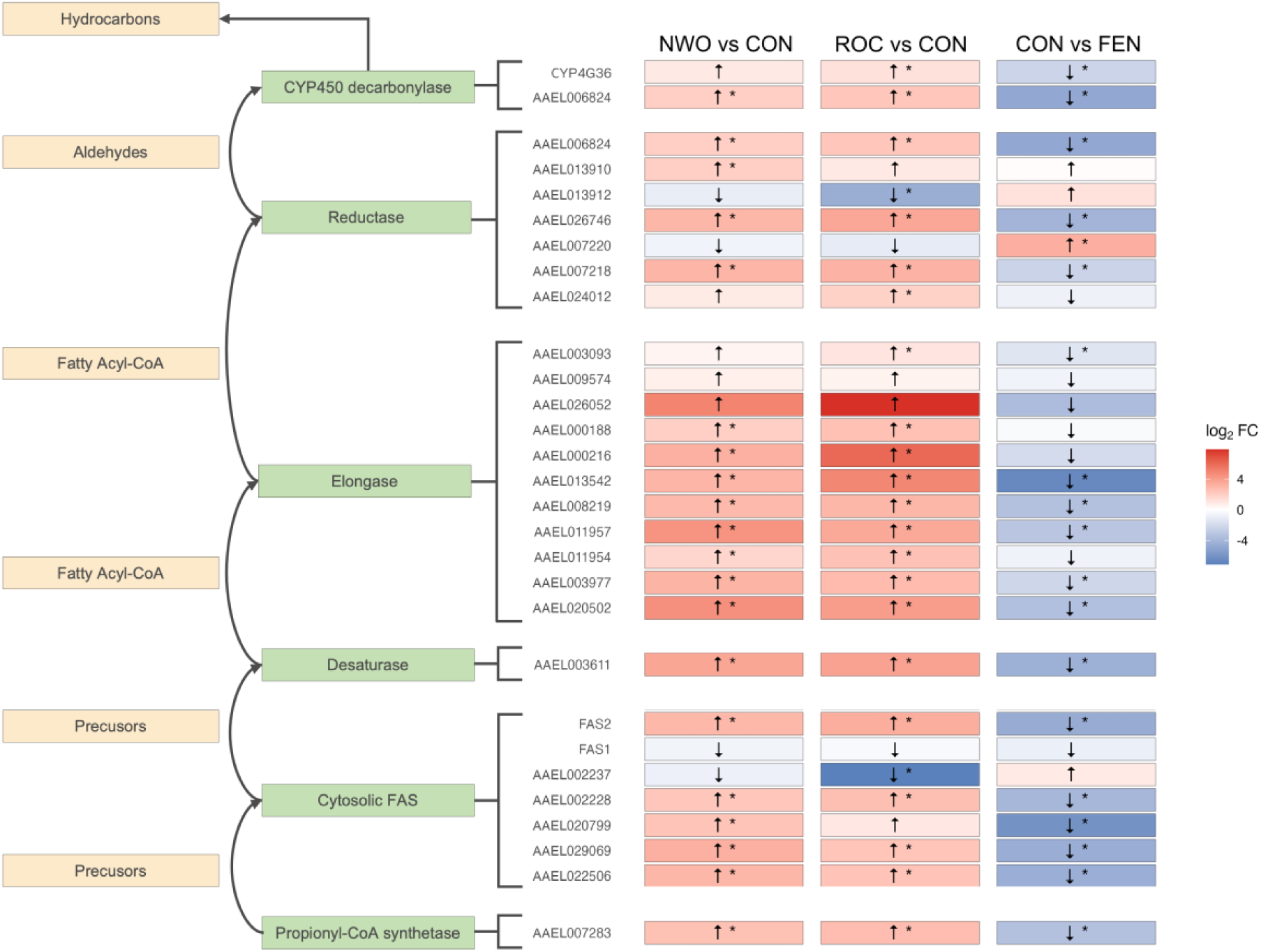
Expression data across the comparisons for cuticular-related genes involved in hydrocarbon synthesis, along with compounds produced at each step of the pathway. Adapted from Grigoraki *et al.* [145]. Direction of expression is shown by the arrows, and significance according to adjusted p value and log2 fold change indicated by an asterisk. The 28 *Ae. aegypti* genes shown are homologous for the genes in *An. gambiae* implicated in the pathway from the same study.

**Table S1.**
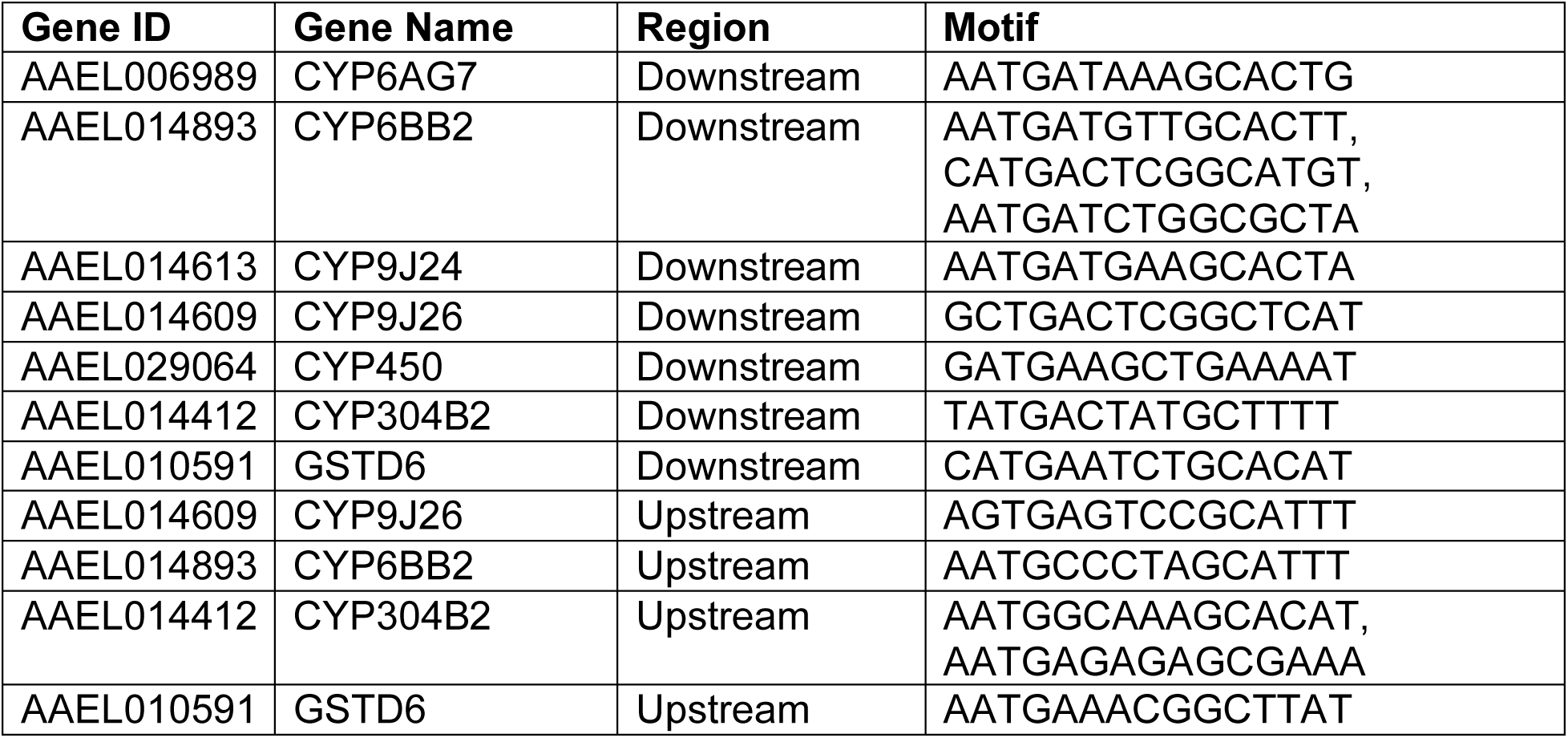
cnc-Maf-S *Drosophila melanogaster* binding motifs identified within the 6 CYP450s found to be upregulated across all comparisons, and *GSTD1*.

## Supplementary Data

File S1. Alignment of the 13 *Ae. aegypti VGSC* transcripts (AAEL023266) and *M. domestica VGSC* transcript (MDOA002080-RB) using Clustal Omega with default options. Locations of SNPs in the Angolan populations are highlighted in purple.

AAEL023266-RL was used to report *Ae. aegypti* amino acid substitution locations.

File S2. Gene set enrichment analysis according to molecular function GO terms of upregulated genes across the three comparisons.

File S3. Results of differential expression analysis across the three comparisons in this study. Grouping of genes according to families referred to in-text is shown. Genes which were significantly differentially expressed across all comparisons are also listed.

File S4. Per-gene Fst values between all three comparisons.

File S5. All detected non-synonymous mutations in the six upregulated genes with significant Fst in the CON vs FEN comparison. Also shown are detected non-synonymous mutations in the nAChR (AAEL004935) with significant Fst in the CON vs FEN comparison, alongside *VGSC*, *ACE-1*, and *RDL*. Blank cells indicate missing data. Absence of each variant in the two susceptible colonies (NWO and ROC) is displayed.

File S6. Homologous genes in Ae. aegypti for genes in An. gambiae implicated in the hydrocarbon synthesis pathway leading to cuticular deposition by Grigoraki *et al.* [145]. Genes with synetity are labelled. Where there were instances of 1-to-many homologous, all genes are listed.

## Author contributions

Conceptualisation: DW, JP Data Curation: HAY, XGB, JP Formal Analysis: HAY, XGB Funding Acquisition: JP, DW

Investigation: HAY, JMFO, AM, LG, JP, CAS, RP, ADT Methodology: JP, DW, CAS, HAY, ERL, MJIP

Project Administration: JP, DW Resources: JP, CAS, MJIP, DW Software: HAY, ERL Supervision: DW, MJIP, ERL Validation: HAY, ERL, JMFO, AM Visualisation: HAY

Writing - Original Draft Preparation: HAY Writing - Review & Editing: ALL

