## Supplementary material for "Transcriptomic profiling reveals multiple mechanisms of insecticide resistance in *Aedes aegypti* from Angola": FileS1

S1 File. Alignment of the 13 *Ae. aegypti VGSC* transcripts (AAEL023266) and *M. domestica VGSC* transcript (MDOA002080-RB) using Clustal Omega with default options. Location of called SNPs in the Angolan populations are highlighted in purple. AAEL023266-RL was used to report *Ae. aegypti* amino acid substitution locations.

```
CLUSTAL O(1.2.4) multiple sequence alignment


MDOA002080-RB      MTEDSDSISEEERSLFRPFTRESLLQIEQRIAEHEKQ-KELERKRA---------AEGEQ	50
AAEL023266-RG      MTEDSDSISEEERSLFRPFTRESLAAIERRIADAEAKQRELEKKRAEGETGFGRKKKKKE	60
AAEL023266-RE      MTEDSDSISEEERSLFRPFTRESLAAIERRIADAEAKQRELEKKRAEG-----------E	49
AAEL023266-RH      MTEDSDSISEEERSLFRPFTRESLAAIERRIADAEAKQRELEKKRAEG-----------E	49
AAEL023266-RF      MTEDSDSISEEERSLFRPFTRESLAAIERRIADAEAKQRELEKKRAEG-----------E	49
AAEL023266-RC      MTEDSDSISEEERSLFRPFTRESLAAIERRIADAEAKQRELEKKRAEG-----------E	49
AAEL023266-RD      MTEDSDSISEEERSLFRPFTRESLAAIERRIADAEAKQRELEKKRAEG-----------E	49
AAEL023266-RI      MTEDSDSISEEERSLFRPFTRESLAAIERRIADAEAKQRELEKKRAEG-----------E	49
AAEL023266-RM      MTEDSDSISEEERSLFRPFTRESLAAIERRIADAEAKQRELEKKRAEG-----------E	49
AAEL023266-RA      MTEDSDSISEEERSLFRPFTRESLAAIERRIADAEAKQRELEKKRAEG-----------E	49
AAEL023266-RB      MTEDSDSISEEERSLFRPFTRESLAAIERRIADAEAKQRELEKKRAEG-----------E	49
AAEL023266-RL      MTEDSDSISEEERSLFRPFTRESLAAIERRIADAEAKQRELEKKRAEG-----------E	49
AAEL023266-RJ      MTEDSDSISEEERSLFRPFTRESLAAIERRIADAEAKQRELEKKRAEG-----------E	49
AAEL023266-RK      MTEDSDSISEEERSLFRPFTRESLAAIERRIADAEAKQRELEKKRAEG-----------E	49
                   ************************  **:***: * : :***:***             :

MDOA002080-RB      IRYDDEDEDEGPQPDPTLEQGVPIPVRMQGSFPPELASTPLEDIDPFYSNVLTFVVISKG	110
AAEL023266-RG      IRYDDEDEDEGPQPDSTLEQGVPIPVRMQGSFPPELASTPLEDIDSYYANQRTFVVVSKG	120
AAEL023266-RE      IRYDDEDEDEGPQPDSTLEQGVPIPVRMQGSFPPELASTPLEDIDSYYANQRTFVVVSKG	109
AAEL023266-RH      IRYDDEDEDEGPQPDSTLEQGVPIPVRMQGSFPPELASTPLEDIDSYYANQRTFVVVSKG	109
AAEL023266-RF      IRYDDEDEDEGPQPDSTLEQGVPIPVRMQGSFPPELASTPLEDIDSYYANQRTFVVVSKG	109
AAEL023266-RC      IRYDDEDEDEGPQPDSTLEQGVPIPVRMQGSFPPELASTPLEDIDSYYANQRTFVVVSKG	109
AAEL023266-RD      IRYDDEDEDEGPQPDSTLEQGVPIPVRMQGSFPPELASTPLEDIDSYYANQRTFVVVSKG	109
AAEL023266-RI      IRYDDEDEDEGPQPDSTLEQGVPIPVRMQGSFPPELASTPLEDIDSYYANQRTFVVVSKG	109
AAEL023266-RM      IRYDDEDEDEGPQPDSTLEQGVPIPVRMQGSFPPELASTPLEDIDSYYANQRTFVVVSKG	109
AAEL023266-RA      IRYDDEDEDEGPQPDSTLEQGVPIPVRMQGSFPPELASTPLEDIDSYYANQRTFVVVSKG	109
AAEL023266-RB      IRYDDEDEDEGPQPDSTLEQGVPIPVRMQGSFPPELASTPLEDIDSYYANQRTFVVVSKG	109
AAEL023266-RL      IRYDDEDEDEGPQPDSTLEQGVPIPVRMQGSFPPELASTPLEDIDSYYANQRTFVVVSKG	109
AAEL023266-RJ      IRYDDEDEDEGPQPDSTLEQGVPIPVRMQGSFPPELASTPLEDIDSYYANQRTFVVVSKG	109
AAEL023266-RK      IRYDDEDEDEGPQPDSTLEQGVPIPVRMQGSFPPELASTPLEDIDSYYANQRTFVVVSKG	109
                   *************** ***************************** :*:*  ****:***

MDOA002080-RB      KDIFRFSASKAMWLLDPFNPIRRVAIYILVHPLFSLFIITTILTNCILMIMPTTPTVEST	170
AAEL023266-RG      KDIFRFSATNALYVLDPFNPIRRVAIYILVHPLFSFFIITTILTNCILMIMPSTPTVEST	180
AAEL023266-RE      KDIFRFSATNALYVLDPFNPIRRVAIYILVHPLFSFFIITTILTNCILMIMPSTPTVEST	169
AAEL023266-RH      KDIFRFSATNALYVLDPFNPIRRVAIYILVHPLFSFFIITTILTNCILMIMPSTPTVEST	169
AAEL023266-RF      KDIFRFSATNALYVLDPFNPIRRVAIYILVHPLFSFFIITTILTNCILMIMPSTPTVEST	169
AAEL023266-RC      KDIFRFSATNALYVLDPFNPIRRVAIYILVHPLFSFFIITTILTNCILMIMPSTPTVEST	169
AAEL023266-RD      KDIFRFSATNALYVLDPFNPIRRVAIYILVHPLFSFFIITTILTNCILMIMPSTPTVEST	169
AAEL023266-RI      KDIFRFSATNALYVLDPFNPIRRVAIYILVHPLFSFFIITTILTNCILMIMPSTPTVEST	169
AAEL023266-RM      KDIFRFSATNALYVLDPFNPIRRVAIYILVHPLFSFFIITTILTNCILMIMPSTPTVEST	169
AAEL023266-RA      KDIFRFSATNALYVLDPFNPIRRVAIYILVHPLFSFFIITTILTNCILMIMPSTPTVEST	169
AAEL023266-RB      KDIFRFSATNALYVLDPFNPIRRVAIYILVHPLFSFFIITTILTNCILMIMPSTPTVEST	169
AAEL023266-RL      KDIFRFSATNALYVLDPFNPIRRVAIYILVHPLFSFFIITTILTNCILMIMPSTPTVEST	169
AAEL023266-RJ      KDIFRFSATNALYVLDPFNPIRRVAIYILVHPLFSFFIITTILTNCILMIMPSTPTVEST	169
AAEL023266-RK      KDIFRFSATNALYVLDPFNPIRRVAIYILVHPLFSFFIITTILTNCILMIMPSTPTVEST	169
                   ********::*:::*********************:****************:*******

MDOA002080-RB      EVIFTGIYTFESAVKVMARGFILCPFTYLRDAWNWLDFVVIALAYVTMGIDLGNLAALRT	230
AAEL023266-RG      EVIFTGIYTFESAVKVMARGFILQPFTYLRDAWNWLDFVVIALAYVTMGIDLGNLAALRT	240
AAEL023266-RE      EVIFTGIYTFESAVKVMARGFILQPFTYLRDAWNWLDFVVIALAYVTMGIDLGNLAALRT	229
AAEL023266-RH      EVIFTGIYTFESAVKVMARGFILQPFTYLRDAWNWLDFVVIALAYVTMGIDLGNLAALRT	229
AAEL023266-RF      EVIFTGIYTFESAVKVMARGFILQPFTYLRDAWNWLDFVVIALAYVTMGIDLGNLAALRT	229
AAEL023266-RC      EVIFTGIYTFESAVKVMARGFILQPFTYLRDAWNWLDFVVIALAYVTMGIDLGNLAALRT	229
AAEL023266-RD      EVIFTGIYTFESAVKVMARGFILQPFTYLRDAWNWLDFVVIALAYVTMGIDLGNLAALRT	229
AAEL023266-RI      EVIFTGIYTFESAVKVMARGFILQPFTYLRDAWNWLDFVVIALAYVTMGIDLGNLAALRT	229
AAEL023266-RM      EVIFTGIYTFESAVKVMARGFILQPFTYLRDAWNWLDFVVIALAYVTMGIDLGNLAALRT	229
AAEL023266-RA      EVIFTGIYTFESAVKVMARGFILQPFTYLRDAWNWLDFVVIALAYVTMGIDLGNLAALRT	229
AAEL023266-RB      EVIFTGIYTFESAVKVMARGFILQPFTYLRDAWNWLDFVVIALAYVTMGIDLGNLAALRT	229
AAEL023266-RL      EVIFTGIYTFESAVKVMARGFILQPFTYLRDAWNWLDFVVIALAYVTMGIDLGNLAALRT	229
AAEL023266-RJ      EVIFTGIYTFESAVKVMARGFILQPFTYLRDAWNWLDFVVIALAYVTMGIDLGNLAALRT	229
AAEL023266-RK      EVIFTGIYTFESAVKVMARGFILQPFTYLRDAWNWLDFVVIALAYVTMGIDLGNLAALRT	229
                   *********************** ************************************

MDOA002080-RB      FRVLRALKTVAIVPGLKTIVGAVIESVKNLRDVIILTMFSLSVFALMGLQIYMGVLTQKC	290
AAEL023266-RG      FRVLRALKTVAIVPGLKTIVGAVIESVKNLRDVIILTMFSLSVFALMGLQIYMGVLTQKC	300
AAEL023266-RE      FRVLRALKTVAIVPGLKTIVGAVIESVKNLRDVIILTMFSLSVFALMGLQIYMGVLTQKC	289
AAEL023266-RH      FRVLRALKTVAIVPGLKTIVGAVIESVKNLRDVIILTMFSLSVFALMGLQIYMGVLTQKC	289
AAEL023266-RF      FRVLRALKTVAIVPGLKTIVGAVIESVKNLRDVIILTMFSLSVFALMGLQIYMGVLTQKC	289
AAEL023266-RC      FRVLRALKTVAIVPGLKTIVGAVIESVKNLRDVIILTMFSLSVFALMGLQIYMGVLTQKC	289
AAEL023266-RD      FRVLRALKTVAIVPGLKTIVGAVIESVKNLRDVIILTMFSLSVFALMGLQIYMGVLTQKC	289
AAEL023266-RI      FRVLRALKTVAIVPGLKTIVGAVIESVKNLRDVIILTMFSLSVFALMGLQIYMGVLTQKC	289
AAEL023266-RM      FRVLRALKTVAIVPGLKTIVGAVIESVKNLRDVIILTMFSLSVFALMGLQIYMGVLTQKC	289
AAEL023266-RA      FRVLRALKTVAIVPGLKTIVGAVIESVKNLRDVIILTMFSLSVFALMGLQIYMGVLTQKC	289
AAEL023266-RB      FRVLRALKTVAIVPGLKTIVGAVIESVKNLRDVIILTMFSLSVFALMGLQIYMGVLTQKC	289
AAEL023266-RL      FRVLRALKTVAIVPGLKTIVGAVIESVKNLRDVIILTMFSLSVFALMGLQIYMGVLTQKC	289
AAEL023266-RJ      FRVLRALKTVAIVPGLKTIVGAVIESVKNLRDVIILTMFSLSVFALMGLQIYMGVLTQKC	289
AAEL023266-RK      FRVLRALKTVAIVPGLKTIVGAVIESVKNLRDVIILTMFSLSVFALMGLQIYMGVLTQKC	289
                   ************************************************************

MDOA002080-RB      IKRFPLDGSWGNLTDENWFLHNSNSSNWFTENDGESYPVCGNVSGAGQCGEDYVCLQGFG	350
AAEL023266-RG      IREFPMDGSWGNLSDENWERFNNNDSNWYFSETGD-TPLCGNSSGAGQCEEGYICLQGYG	359
AAEL023266-RE      IREFPMDGSWGNLSDENWERFNNNDSNWYFSETGD-TPLCGNSSGAGQCEEGYICLQGYG	348
AAEL023266-RH      IREFPMDGSWGNLSDENWERFNNNDSNWYFSETGD-TPLCGNSSGAGQCEEGYICLQGYG	348
AAEL023266-RF      IREFPMDGSWGNLSDENWERFNNNDSNWYFSETGD-TPLCGNSSGAGQCEEGYICLQGYG	348
AAEL023266-RC      IREFPMDGSWGNLSDENWERFNNNDSNWYFSETGD-TPLCGNSSGAGQCEEGYICLQGYG	348
AAEL023266-RD      IREFPMDGSWGNLSDENWERFNNNDSNWYFSETGD-TPLCGNSSGAGQCEEGYICLQGYG	348
AAEL023266-RI      IREFPMDGSWGNLSDENWERFNNNDSNWYFSETGD-TPLCGNSSGAGQCEEGYICLQGYG	348
AAEL023266-RM      IREFPMDGSWGNLSDENWERFNNNDSNWYFSETGD-TPLCGNSSGAGQCEEGYICLQGYG	348
AAEL023266-RA      IREFPMDGSWGNLSDENWERFNNNDSNWYFSETGD-TPLCGNSSGAGQCEEGYICLQGYG	348
AAEL023266-RB      IREFPMDGSWGNLSDENWERFNNNDSNWYFSETGD-TPLCGNSSGAGQCEEGYICLQGYG	348
AAEL023266-RL      IREFPMDGSWGNLSDENWERFNNNDSNWYFSETGD-TPLCGNSSGAGQCEEGYICLQGYG	348
AAEL023266-RJ      IREFPMDGSWGNLSDENWERFNNNDSNWYFSETGD-TPLCGNSSGAGQCEEGYICLQGYG	348
AAEL023266-RK      IREFPMDGSWGNLSDENWERFNNNDSNWYFSETGD-TPLCGNSSGAGQCEEGYICLQGYG	348
                   *:.**:*******:****  .*.*.***: .: *:  *:*** ****** *.*:****:*

MDOA002080-RB      PNPNYDYTSFDSFGWAFLSAFRLMTQDFWEDLYQHVLQAAGPWHMLFFIVIIFLGSFYLV	410
AAEL023266-RG      DNPNYGYTSFDTFGWAFLSAFRLMTQDYWENLYQLVLRSAGPWHMLFFIVIIFLGSFYLV	419
AAEL023266-RE      DNPNYGYTSFDTFGWAFLSAFRLMTQDYWENLYQLVLRSAGPWHMLFFIVIIFLGSFYLV	408
AAEL023266-RH      DNPNYGYTSFDTFGWAFLSAFRLMTQDYWENLYQLVLRSAGPWHMLFFIVIIFLGSFYLV	408
AAEL023266-RF      DNPNYGYTSFDTFGWAFLSAFRLMTQDYWENLYQLVLRSAGPWHMLFFIVIIFLGSFYLV	408
AAEL023266-RC      DNPNYGYTSFDTFGWAFLSAFRLMTQDYWENLYQLVLRSAGPWHMLFFIVIIFLGSFYLV	408
AAEL023266-RD      DNPNYGYTSFDTFGWAFLSAFRLMTQDYWENLYQLVLRSAGPWHMLFFIVIIFLGSFYLV	408
AAEL023266-RI      DNPNYGYTSFDTFGWAFLSAFRLMTQDYWENLYQLVLRSAGPWHMLFFIVIIFLGSFYLV	408
AAEL023266-RM      DNPNYGYTSFDTFGWAFLSAFRLMTQDYWENLYQLVLRSAGPWHMLFFIVIIFLGSFYLV	408
AAEL023266-RA      DNPNYGYTSFDTFGWAFLSAFRLMTQDYWENLYQLVLRSAGPWHMLFFIVIIFLGSFYLV	408
AAEL023266-RB      DNPNYGYTSFDTFGWAFLSAFRLMTQDYWENLYQLVLRSAGPWHMLFFIVIIFLGSFYLV	408
AAEL023266-RL      DNPNYGYTSFDTFGWAFLSAFRLMTQDYWENLYQLVLRSAGPWHMLFFIVIIFLGSFYLV	408
AAEL023266-RJ      DNPNYGYTSFDTFGWAFLSAFRLMTQDYWENLYQLVLRSAGPWHMLFFIVIIFLGSFYLV	408
AAEL023266-RK      DNPNYGYTSFDTFGWAFLSAFRLMTQDYWENLYQLVLRSAGPWHMLFFIVIIFLGSFYLV	408
                    ****.*****:***************:**:*** **::*********************

MDOA002080-RB      NLILAIVAMSYDELQKKAEEEEAAEEEAIREAEEAAAAKAAKLEERANVAAQAAQDAADA	470
AAEL023266-RG      NLILAIVAMSYDELQKKAEEEEAAEEEALREAEEAAAAKAAKLEAQAAA-----------	468
AAEL023266-RE      NLILAIVAMSYDELQKKAEEEEAAEEEALREAEEAAAAKAAKLEAQAAA-----------	457
AAEL023266-RH      NLILAIVAMSYDELQKKAEEEEAAEEEALREAEEAAAAKAAKLEAQAAA-----------	457
AAEL023266-RF      NLILAIVAMSYDELQKKAEEEEAAEEEALREAEEAAAAKAAKLEAQAAA-----------	457
AAEL023266-RC      NLILAIVAMSYDELQKKAEEEEAAEEEALREAEEAAAAKAAKLEAQAAA-----------	457
AAEL023266-RD      NLILAIVAMSYDELQKKAEEEEAAEEEALREAEEAAAAKAAKLEAQAAA-----------	457
AAEL023266-RI      NLILAIVAMSYDELQKKAEEEEAAEEEALREAEEAAAAKAAKLEAQAAA-----------	457
AAEL023266-RM      NLILAIVAMSYDELQKKAEEEEAAEEEALREAEEAAAAKAAKLEAQAAA-----------	457
AAEL023266-RA      NLILAIVAMSYDELQKKAEEEEAAEEEALREAEEAAAAKAAKLEAQAAA-----------	457
AAEL023266-RB      NLILAIVAMSYDELQKKAEEEEAAEEEALREAEEAAAAKAAKLEAQAAA-----------	457
AAEL023266-RL      NLILAIVAMSYDELQKKAEEEEAAEEEALREAEEAAAAKAAKLEAQAAA-----------	457
AAEL023266-RJ      NLILAIVAMSYDELQKKAEEEEAAEEEALREAEEAAAAKAAKLEAQAAA-----------	457
AAEL023266-RK      NLILAIVAMSYDELQKKAEEEEAAEEEALREAEEAAAAKAAKLEAQAAA-----------	457
                   ****************************:*************** :* .

MDOA002080-RB      AAAALHPEMAKSP-TYSCISYELFVGGEKGNDDNNKEKMSIRSVEVESESVSVIQRQPAP	529
AAEL023266-RG      AAAAANPEIAKSPSDFSCHSYELFVNQEKGNDDNNKEKMSIRSEGLESVSEITRTTAPTA	528
AAEL023266-RE      AAAAANPEIAKSPSDFSCHSYELFVNQEKGNDDNNKEKMSIRSEGLESVSEITRTTAPTA	517
AAEL023266-RH      AAAAANPEIAKSPSDFSCHSYELFVNQEKGNDDNNKEKMSIRSEGLESVSEITRTTAPTA	517
AAEL023266-RF      AAAAANPEIAKSPSDFSCHSYELFVNQEKGNDDNNKEKMSIRSEGLESVSEITRTTAPTA	517
AAEL023266-RC      AAAAANPEIAKSPSDFSCHSYELFVNQEKGNDDNNKEKMSIRSEGLESVSEITRTTAPTA	517
AAEL023266-RD      AAAAANPEIAKSPSDFSCHSYELFVNQEKGNDDNNKEKMSIRSEGLESVSEITRTTAPTA	517
AAEL023266-RI      AAAAANPEIAKSPSDFSCHSYELFVNQEKGNDDNNKEKMSIRSEGLES------------	505
AAEL023266-RM      AAAAANPEIAKSPSDFSCHSYELFVNQEKGNDDNNKEKMSIRSEGLESVSEITRTTAPTA	517
AAEL023266-RA      AAAAANPEIAKSPSDFSCHSYELFVNQEKGNDDNNKEKMSIRSEGLES------------	505
AAEL023266-RB      AAAAANPEIAKSPSDFSCHSYELFVNQEKGNDDNNKEKMSIRSEGLESVSEITRTTAPTA	517
AAEL023266-RL      AAAAANPEIAKSPSDFSCHSYELFVNQEKGNDDNNKEKMSIRSEGLESVSEITRTTAPTA	517
AAEL023266-RJ      AAAAANPEIAKSPSDFSCHSYELFVNQEKGNDDNNKEKMSIRSEGLESVSEITRTTAPTA	517
AAEL023266-RK      AAAAANPEIAKSPSDFSCHSYELFVNQEKGNDDNNKEKMSIRSEGLESVSEITRTTAPTA	517
                   **** :**:****  :** ******. ****************  :**

MDOA002080-RB      TTAPATKVR-------KVSTTSLSLPGSPFNLRRGSRSSHKYTIRNGRGRF-GIPGSDRK	581
AAEL023266-RG      TAAGTAKARKVSAGVAAFQKASLSLPGSPFNLRRGSRGSHQFTIRNGRGRFVGVPGSDRK	588
AAEL023266-RE      TAAGTAKARKVSAGVAAFQKASLSLPGSPFNLRRGSRGSHQFTIRNGRGRFVGVPGSDRK	577
AAEL023266-RH      TAAGTAKARKVSAGVAAFQKASLSLPGSPFNLRRGSRGSHQFTIRNGRGRFVGVPGSDRK	577
AAEL023266-RF      TAAGTAKARKVS----------------------------AFTIRNGRGRFVGVPGSDRK	549
AAEL023266-RC      TAAGTAKARKVSA-------ASLSLPGSPFNLRRGSRGSHQFTIRNGRGRFVGVPGSDRK	570
AAEL023266-RD      TAAGTAKARKVSA-------ASLSLPGSPFNLRRGSRGSHQFTIRNGRGRFVGVPGSDRK	570
AAEL023266-RI      --------------------ASLSLPGSPFNLRRGSRGSHQFTIRNGRGRFVGVPGSDRK	545
AAEL023266-RM      TAAGTAKARKVSA-------ASLSLPGSPFNLRRGSRGSHQFTIRNGRGRFVGVPGSDRK	570
AAEL023266-RA      --------------------ASLSLPGSPFNLRRGSRGSHQFTIRNGRGRFVGVPGSDRK	545
AAEL023266-RB      TAAGTAKARKVSA-------ASLSLPGSPFNLRRGSRGSHQFTIRNGRGRFVGVPGSDRK	570
AAEL023266-RL      TAAGTAKARKVSAGVAAFQKASLSLPGSPFNLRRGSRGSHQFTIRNGRGRFVGVPGSDRK	577
AAEL023266-RJ      TAAGTAKARKVSA-------ASLSLPGSPFNLRRGSRGSHQFTIRNGRGRFVGVPGSDRK	570
AAEL023266-RK      TAAGTAKARKVSA-------ASLSLPGSPFNLRRGSRGSHQFTIRNGRGRFVGVPGSDRK	570
                                                            :********* *:******

MDOA002080-RB      PLVLQTYQDAQQHLPYADDSNAVTPMSEENGAIIVPAYYCNLGSRHSSYTSHQSRISYTS	641
AAEL023266-RG      PLVLSTYLDAQEHLPYADDSNAVTPMSEENGAIIVPVYYANLGSRHSSYTSHQSRISYTS	648
AAEL023266-RE      PLVLSTYLDAQEHLPYADDSNAVTPMSEENGAIIVPVYYANLGSRHSSYTSHQSRISYTS	637
AAEL023266-RH      PLVLSTYLDAQEHLPYADDSNAVTPMSEENGAIIVPVYYANLGSRHSSYTSHQSRISYTS	637
AAEL023266-RF      PLVLSTYLDAQEHLPYADDSNAVTPMSEENGAIIVPVYYANLGSRHSSYTSHQSRISYTS	609
AAEL023266-RC      PLVLSTYLDAQEHLPYADDSNAVTPMSEENGAIIVPVYYANLGSRHSSYTSHQSRISYTS	630
AAEL023266-RD      PLVLSTYLDAQEHLPYADDSNAVTPMSEENGAIIVPVYYANLGSRHSSYTSHQSRISYTS	630
AAEL023266-RI      PLVLSTYLDAQEHLPYADDSNAVTPMSEENGAIIVPVYYANLGSRHSSYTSHQSRISYTS	605
AAEL023266-RM      PLVLSTYLDAQEHLPYADDSNAVTPMSEENGAIIVPVYYANLGSRHSSYTSHQSRISYTS	630
AAEL023266-RA      PLVLSTYLDAQEHLPYADDSNAVTPMSEENGAIIVPVYYANLGSRHSSYTSHQSRISYTS	605
AAEL023266-RB      PLVLSTYLDAQEHLPYADDSNAVTPMSEENGAIIVPVYYANLGSRHSSYTSHQSRISYTS	630
AAEL023266-RL      PLVLSTYLDAQEHLPYADDSNAVTPMSEENGAIIVPVYYANLGSRHSSYTSHQSRISYTS	637
AAEL023266-RJ      PLVLSTYLDAQEHLPYADDSNAVTPMSEENGAIIVPVYYANLGSRHSSYTSHQSRISYTS	630
AAEL023266-RK      PLVLSTYLDAQEHLPYADDSNAVTPMSEENGAIIVPVYYANLGSRHSSYTSHQSRISYTS	630
                   ****.** ***:************************.**.********************

MDOA002080-RB      HGDLLGGMAAMGASTMTKESKLRSRNTRNQSIGAATNGGSSTAGGGYPDANHKEQRDYEM	701
AAEL023266-RG      HGDLLGGMT--------KESRLRNRSARNTNHSIVPPPNMSGPNMSYVDSNHKGQRDFDM	700
AAEL023266-RE      HGDLLGGMT--------KESRLRNRSARNTNHSIVPPPNMSGPNMSYVDSNHKGQRDFDM	689
AAEL023266-RH      HGDLLGGMT--------KESRLRNRSARNTNHSIVPPPNMSGPNMSYVDSNHKGQRDFDM	689
AAEL023266-RF      HGDLLGGMT--------KESRLRNRSARNTNHSIVPPPNMSGPNMSYVDSNHKGQRDFDM	661
AAEL023266-RC      HGDLLGGMT--------KESRLRNRSARNTNHSIVPPPNMSGPNMSYVDSNHKGQRDFDM	682
AAEL023266-RD      HGDLLGGMT--------KESRLRNRSARNTNHSIVPPPNMSGPNMSYVDSNHKGQRDFDM	682
AAEL023266-RI      HGDLLGGMT--------KESRLRNRSARNTNHSIVPPPNMSGPNMSYVDSNHKGQRDFDM	657
AAEL023266-RM      HGDLLGGMT--------KESRLRNRSARNTNHSIVPPPNMSGPNMSYVDSNHKGQRDFDM	682
AAEL023266-RA      HGDLLGGMT--------KESRLRNRSARNTNHSIVPPPNMSGPNMSYVDSNHKGQRDFDM	657
AAEL023266-RB      HGDLLGGMT--------KESRLRNRSARNTNHSIVPPPNMSGPNMSYVDSNHKGQRDFDM	682
AAEL023266-RL      HGDLLGGMT--------KESRLRNRSARNTNHSIVPPPNMSGPNMSYVDSNHKGQRDFDM	689
AAEL023266-RJ      HGDLLGGMT--------KESRLRNRSARNTNHSIVPPPNMSGPNMSYVDSNHKGQRDFDM	682
AAEL023266-RK      HGDLLGGMT--------KESRLRNRSARNTNHSIVPPPNMSGPNMSYVDSNHKGQRDFDM	682
                   ********:        ***:**.*.:** . . .   . *  . .* *:*** ***::*

MDOA002080-RB      GQDYTDEAGKIKHHDNPFIEPVQTQTVVDMKDVMVLNDIIEQAAGRHSRASERG------	755
AAEL023266-RG      SQDCTDEAGKIKHNDNPFIEPSQTQTVVDMKDVMVLNDIIEQAAGRHSRASDHG------	754
AAEL023266-RE      SQDCTDEAGKIKHNDNPFIEPSQTQTVVDMKDVMVLNDIIEQAAGRHSRASDHGVSVYYF	749
AAEL023266-RH      SQDCTDEAGKIKHNDNPFIEPSQTQTVVDMKDVMVLNDIIEQAAGRHSRASDHG------	743
AAEL023266-RF      SQDCTDEAGKIKHNDNPFIEPSQTQTVVDMKDVMVLNDIIEQAAGRHSRASDHG------	715
AAEL023266-RC      SQDCTDEAGKIKHNDNPFIEPSQTQTVVDMKDVMVLNDIIEQAAGRHSRASDHGVSVYYF	742
AAEL023266-RD      SQDCTDEAGKIKHNDNPFIEPSQTQTVVDMKDVMVLNDIIEQAAGRHSRASDHGVSVYYF	742
AAEL023266-RI      SQDCTDEAGKIKHNDNPFIEPSQTQTVVDMKDVMVLNDIIEQAAGRHSRASDHGVSVYYF	717
AAEL023266-RM      SQDCTDEAGKIKHNDNPFIEPSQTQTVVDMKDVMVLNDIIEQAAGRHSRASDHGVSVYYF	742
AAEL023266-RA      SQDCTDEAGKIKHNDNPFIEPSQTQTVVDMKDVMVLNDIIEQAAGRHSRASDHGVSVYYF	717
AAEL023266-RB      SQDCTDEAGKIKHNDNPFIEPSQTQTVVDMKDVMVLNDIIEQAAGRHSRASDHGVSVYYF	742
AAEL023266-RL      SQDCTDEAGKIKHNDNPFIEPSQTQTVVDMKDVMVLNDIIEQAAGRHSRASDHGVSVYYF	749
AAEL023266-RJ      SQDCTDEAGKIKHNDNPFIEPSQTQTVVDMKDVMVLNDIIEQAAGRHSRASDHGVSVYYF	742
AAEL023266-RK      SQDCTDEAGKIKHNDNPFIEPSQTQTVVDMKDVMVLNDIIEQAAGRHSRASDHGVSVYYF	742
                   .** *********:******* *****************************::*

MDOA002080-RB      --EDDDEDGPTFKDIALEYILKGIEIFCVWDCCWVWLKFQEWVSFIVFDPFVELFITLCI	813
AAEL023266-RG      --EDDDEDGPTFKDKALEFTMRMIDVFCVWDCCWVWLKFQEWVAFIVFDPFVELFITLCI	812
AAEL023266-RE      PTEDDDEDGPTFKDKALEFTMRMIDVFCVWDCCWVWLKFQEWVAFIVFDPFVELFITLCI	809
AAEL023266-RH      --EDDDEDGPTFKDKALEFTMRMIDVFCVWDCCWVWLKFQEWVAFIVFDPFVELFITLCI	801
AAEL023266-RF      --EDDDEDGPTFKDKALEFTMRMIDVFCVWDCCWVWLKFQEWVAFIVFDPFVELFITLCI	773
AAEL023266-RC      PTEDDDEDGPTFKDKALEFTMRMIDVFCVWDCCWVWLKFQEWVAFIVFDPFVELFITLCI	802
AAEL023266-RD      PTEDDDEDGPTFKDKALEFTMRMIDVFCVWDCCWVWLKFQEWVAFIVFDPFVELFITLCI	802
AAEL023266-RI      PTEDDDEDGPTFKDKALEFTMRMIDVFCVWDCCWVWLKFQEWVAFIVFDPFVELFITLCI	777
AAEL023266-RM      PTEDDDEDGPTFKDKALEFTMRMIDVFCVWDCCWVWLKFQEWVAFIVFDPFVELFITLCI	802
AAEL023266-RA      PTEDDDEDGPTFKDKALEFTMRMIDVFCVWDCCWVWLKFQEWVAFIVFDPFVELFITLCI	777
AAEL023266-RB      PTEDDDEDGPTFKDKALEFTMRMIDVFCVWDCCWVWLKFQEWVAFIVFDPFVELFITLCI	802
AAEL023266-RL      PTEDDDEDGPTFKDKALEFTMRMIDVFCVWDCCWVWLKFQEWVAFIVFDPFVELFITLCI	809
AAEL023266-RJ      PTEDDDEDGPTFKDKALEFTMRMIDVFCVWDCCWVWLKFQEWVAFIVFDPFVELFITLCI	802
AAEL023266-RK      PTEDDDEDGPTFKDKALEFTMRMIDVFCVWDCCWVWLKFQEWVAFIVFDPFVELFITLCI	802
                     ************ ***: :: *::*****************:****************

MDOA002080-RB      VVNTMFMAMDHHDMNPELEKVLKSGNYFFTATFAIEASMKLMAMSPKYYFQEGWNIFDFI	873
AAEL023266-RG      VVNTLFMALDHHDMDPDMERALKSGNYFFTATFAIEATMKLIAMSPKYYFQEGWNIFDFI	872
AAEL023266-RE      VVNTLFMALDHHDMDPDMERALKSGNYFFTATFAIEATMKLIAMSPKYYFQEGWNIFDFI	869
AAEL023266-RH      VVNTLFMALDHHDMDPDMERALKSGNYFFTATFAIEATMKLIAMSPKYYFQEGWNIFDFI	861
AAEL023266-RF      VVNTLFMALDHHDMDPDMERALKSGNYFFTATFAIEATMKLIAMSPKYYFQEGWNIFDFI	833
AAEL023266-RC      VVNTLFMALDHHDMDPDMERALKSGNYFFTATFAIEATMKLIAMSPKYYFQEGWNIFDFI	862
AAEL023266-RD      VVNTLFMALDHHDMDPDMERALKSGNYFFTATFAIEATMKLIAMSPKYYFQEGWNIFDFI	862
AAEL023266-RI      VVNTLFMALDHHDMDPDMERALKSGNYFFTATFAIEATMKLIAMSPKYYFQEGWNIFDFI	837
AAEL023266-RM      VVNTLFMALDHHDMDPDMERALKSGNYFFTATFAIEATMKLIAMSPKYYFQEGWNIFDFI	862
AAEL023266-RA      VVNTLFMALDHHDMDPDMERALKSGNYFFTATFAIEATMKLIAMSPKYYFQEGWNIFDFI	837
AAEL023266-RB      VVNTLFMALDHHDMDPDMERALKSGNYFFTATFAIEATMKLIAMSPKYYFQEGWNIFDFI	862
AAEL023266-RL      VVNTLFMALDHHDMDPDMERALKSGNYFFTATFAIEATMKLIAMSPKYYFQEGWNIFDFI	869
AAEL023266-RJ      VVNTLFMALDHHDMDPDMERALKSGNYFFTATFAIEATMKLIAMSPKYYFQEGWNIFDFI	862
AAEL023266-RK      VVNTLFMALDHHDMDPDMERALKSGNYFFTATFAIEATMKLIAMSPKYYFQEGWNIFDFI	862
                   ****:***:*****:*::*:.****************:***:******************

MDOA002080-RB      IVALSLLELGLEGVQGLSVLRSFRLLRVFKLAKSWPTLNLLISIMGRTMGALGNLTFVLC	933
AAEL023266-RG      IVALSLLELGLEGVQGLSVLRSFRLLRVFKLAKSWPTLNLLISIMGRTVGALGNLTFVLC	932
AAEL023266-RE      IVALSLLELGLEGVQGLSVLRSFRLLRVFKLAKSWPTLNLLISIMGRTVGALGNLTFVLC	929
AAEL023266-RH      IVALSLLELGLEGVQGLSVLRSFRLLRVFKLAKSWPTLNLLISIMGRTVGALGNLTFVLC	921
AAEL023266-RF      IVALSLLELGLEGVQGLSVLRSFRLLRVFKLAKSWPTLNLLISIMGRTMGALGNLTFVLC	893
AAEL023266-RC      IVALSLLELGLEGVQGLSVLRSFRLLRVFKLAKSWPTLNLLISIMGRTVGALGNLTFVLC	922
AAEL023266-RD      IVALSLLELGLEGVQGLSVLRSFRLLRVFKLAKSWPTLNLLISIMGRTMGALGNLTFVLC	922
AAEL023266-RI      IVALSLLELGLEGVQGLSVLRSFRLLRVFKLAKSWPTLNLLISIMGRTMGALGNLTFVLC	897
AAEL023266-RM      IVALSLLELGLEGVQGLSVLRSFRLLRVFKLAKSWPTLNLLISIMGRTVGALGNLTFVLC	922
AAEL023266-RA      IVALSLLELGLEGVQGLSVLRSFRLLRVFKLAKSWPTLNLLISIMGRTMGALGNLTFVLC	897
AAEL023266-RB      IVALSLLELGLEGVQGLSVLRSFRLLRVFKLAKSWPTLNLLISIMGRTMGALGNLTFVLC	922
AAEL023266-RL      IVALSLLELGLEGVQGLSVLRSFRLLRVFKLAKSWPTLNLLISIMGRTMGALGNLTFVLC	929
AAEL023266-RJ      IVALSLLELGLEGVQGLSVLRSFRLLRVFKLAKSWPTLNLLISIMGRTMGALGNLTFVLC	922
AAEL023266-RK      IVALSLLELGLEGVQGLSVLRSFRLLRVFKLAKSWPTLNLLISIMGRTVGALGNLTFVLC	922
                   ************************************************:***********

MDOA002080-RB      IIIFIFAVMGMQLFGKNYIDHKDRFKDHELPRWNFTDFMHSFMIVFRVLCGEWIESMWDC	993
AAEL023266-RG      IIIFIFAVMGMQLFGKNYTDNVDRFPDKDLPRWNFTDFMHSFMIVFRVLCGEWIESMWDC	992
AAEL023266-RE      IIIFIFAVMGMQLFGKNYTDNVDRFPDKDLPRWNFTDFMHSFMIVFRVLCGEWIESMWDC	989
AAEL023266-RH      IIIFIFAVMGMQLFGKNYTDNVDRFPDKDLPRWNFTDFMHSFMIVFRVLCGEWIESMWDC	981
AAEL023266-RF      IIIFIFAVMGMQLFGKNYIDNVDRFPDKDLPRWNFTDFMHSFMIVFRVLCGEWIESMWDC	953
AAEL023266-RC      IIIFIFAVMGMQLFGKNYTDNVDRFPDKDLPRWNFTDFMHSFMIVFRVLCGEWIESMWDC	982
AAEL023266-RD      IIIFIFAVMGMQLFGKNYIDNVDRFPDKDLPRWNFTDFMHSFMIVFRVLCGEWIESMWDC	982
AAEL023266-RI      IIIFIFAVMGMQLFGKNYIDNVDRFPDKDLPRWNFTDFMHSFMIVFRVLCGEWIESMWDC	957
AAEL023266-RM      IIIFIFAVMGMQLFGKNYTDNVDRFPDKDLPRWNFTDFMHSFMIVFRVLCGEWIESMWDC	982
AAEL023266-RA      IIIFIFAVMGMQLFGKNYIDNVDRFPDKDLPRWNFTDFMHSFMIVFRVLCGEWIESMWDC	957
AAEL023266-RB      IIIFIFAVMGMQLFGKNYIDNVDRFPDKDLPRWNFTDFMHSFMIVFRVLCGEWIESMWDC	982
AAEL023266-RL      IIIFIFAVMGMQLFGKNYIDNVDRFPDKDLPRWNFTDFMHSFMIVFRVLCGEWIESMWDC	989
AAEL023266-RJ      IIIFIFAVMGMQLFGKNYIDNVDRFPDKDLPRWNFTDFMHSFMIVFRVLCGEWIESMWDC	982
AAEL023266-RK      IIIFIFAVMGMQLFGKNYTDNVDRFPDKDLPRWNFTDFMHSFMIVFRVLCGEWIESMWDC	982
                   ****************** *: *** *::*******************************

MDOA002080-RB      MYVGDVSCIPFFLATVVIGNLVVLNLFLALLLSNFGSSSLSAPTADNDTNKIAEAFNRIA	1053
AAEL023266-RG      MLVGDVSCIPFFLATVVIGNLVVLNLFLALLLSNFGSSSLSAPTADNETNKIAEAFNRIS	1052
AAEL023266-RE      MLVGDVSCIPFFLATVVIGNLVVLNLFLALLLSNFGSSSLSAPTADNETNKIAEAFNRIS	1049
AAEL023266-RH      MLVGDVSCIPFFLATVVIGNLVVLNLFLALLLSNFGSSSLSAPTADNETNKIAEAFNRIS	1041
AAEL023266-RF      MLVGDVSCIPFFLATVVIGNLVVLNLFLALLLSNFGSSSLSAPTADNETNKIAEAFNRIS	1013
AAEL023266-RC      MLVGDVSCIPFFLATVVIGNLVVLNLFLALLLSNFGSSSLSAPTADNETNKIAEAFNRIS	1042
AAEL023266-RD      MLVGDVSCIPFFLATVVIGNLVVLNLFLALLLSNFGSSSLSAPTADNETNKIAEAFNRIS	1042
AAEL023266-RI      MLVGDVSCIPFFLATVVIGNLVVLNLFLALLLSNFGSSSLSAPTADNETNKIAEAFNRIS	1017
AAEL023266-RM      MLVGDVSCIPFFLATVVIGNLVVLNLFLALLLSNFGSSSLSAPTADNETNKIAEAFNRIS	1042
AAEL023266-RA      MLVGDVSCIPFFLATVVIGNLVVLNLFLALLLSNFGSSSLSAPTADNETNKIAEAFNRIS	1017
AAEL023266-RB      MLVGDVSCIPFFLATVVIGNLVVLNLFLALLLSNFGSSSLSAPTADNETNKIAEAFNRIS	1042
AAEL023266-RL      MLVGDVSCIPFFLATVVIGNLVVLNLFLALLLSNFGSSSLSAPTADNETNKIAEAFNRIS	1049
AAEL023266-RJ      MLVGDVSCIPFFLATVVIGNLVVLNLFLALLLSNFGSSSLSAPTADNETNKIAEAFNRIS	1042
AAEL023266-RK      MLVGDVSCIPFFLATVVIGNLVVLNLFLALLLSNFGSSSLSAPTADNETNKIAEAFNRIS	1042
                   * *********************************************:***********:

MDOA002080-RB      RFKNWVKRNIADCFKLIRNKLTNQISDQP------------------------SEHGDNE	1089
AAEL023266-RG      RFSNWIKSNIANALKFVKNKLTSQIASVQPA-----------------------EHGENE	1089
AAEL023266-RE      RFSNWIKSNIANALKFVKNKLTSQIASVQPA-----------------------EHGENE	1086
AAEL023266-RH      RFSNWIKSNIANALKFVKNKLTSQIASVQPA-----------------------EHGENE	1078
AAEL023266-RF      RFSNWIKSNIANALKFVKNKLTSQIASVQPA-------------GKGVCPCISAEHGENE	1060
AAEL023266-RC      RFSNWIKSNIANALKFVKNKLTSQIASVQPAGEQHNHLSWIWNEGKGVCPCISAEHGENE	1102
AAEL023266-RD      RFSNWIKSNIANALKFVKNKLTSQIASVQPA-----------------------EHGENE	1079
AAEL023266-RI      RFSNWIKSNIANALKFVKNKLTSQIASVQPA-----------------------EHGENE	1054
AAEL023266-RM      RFSNWIKSNIANALKFVKNKLTSQIASVQPA-----------------------------	1073
AAEL023266-RA      RFSNWIKSNIANALKFVKNKLTSQIASVQPAGEQHNHLSWIWNEGKGVCPCISAEHGENE	1077
AAEL023266-RB      RFSNWIKSNIANALKFVKNKLTSQIASVQPA-----------------------EHGENE	1079
AAEL023266-RL      RFSNWIKSNIANALKFVKNKLTSQIASVQPAGEQHNHLSWIWNEGKGVCPCISAEHGENE	1109
AAEL023266-RJ      RFSNWIKSNIANALKFVKNKLTSQIASVQPAGEQHNHLSWIWNEGKGVCPCISAEHGENE	1102
AAEL023266-RK      RFSNWIKSNIANALKFVKNKLTSQIASVQPAGEQHNHLSWIWNEGKGVCPCISAEHGENE	1102
                   **.**:* ***:.:*:::****.**:.

MDOA002080-RB      LELGHDEIMGDGLIKKGMKGETQLEVAIGDGMEFTIHGDMKNNKPKKSKFINNTTMIGNS	1149
AAEL023266-RG      LELTPDDILADGLLKKGVKEHNQLEVAIGDGMEFTIHGDLKNKGKKNKQLMNNSKVIGNS	1149
AAEL023266-RE      LELTPDDILADGLLKKGVKEHNQLEVAIGDGMEFTIHGDLKNKGKKNKQLMNNSKVIGNS	1146
AAEL023266-RH      LELTPDDILADGLLKKGVKEHNQLEVAIGDGMEFTIHGDLKNKGKKNKQLMNNSKVIGNS	1138
AAEL023266-RF      LELTPDDILADGLLKKGVKEHNQLEVAIGDGMEFTIHGDLKNKGKKNKQLMNNSK-----	1115
AAEL023266-RC      LELTPDDILADGLLKKGVKEHNQLEVAIGDGMEFTIHGDLKNKGKKNKQLMNNSKVIGNS	1162
AAEL023266-RD      LELTPDDILADGLLKKGVKEHNQLEVAIGDGMEFTIHGDLKNKGKKNKQLMNNSK-----	1134
AAEL023266-RI      LELTPDDILADGLLKKGVKEHNQLEVAIGDGMEFTIHGDLKNKGKKNKQLMNNSKVIGNS	1114
AAEL023266-RM      -----DDILADGLLKKGVKEHNQLEVAIGDGMEFTIHGDLKNKGKKNKQLMNNSKVIGNS	1128
AAEL023266-RA      LELTPDDILADGLLKKGVKEHNQLEVAIGDGMEFTIHGDLKNKGKKNKQLMNNSKVIGNS	1137
AAEL023266-RB      LELTPDDILADGLLKKGVKEHNQLEVAIGDGMEFTIHGDLKNKGKKNKQLMNNSKVIGNS	1139
AAEL023266-RL      LELTPDDILADGLLKKGVKEHNQLEVAIGDGMEFTIHGDLKNKGKKNKQLMNNSKVIGNS	1169
AAEL023266-RJ      LELTPDDILADGLLKKGVKEHNQLEVAIGDGMEFTIHGDLKNKGKKNKQLMNNSKVIGNS	1162
AAEL023266-RK      LELTPDDILADGLLKKGVKEHNQLEVAIGDGMEFTIHGDLKNKGKKNKQLMNNSKVIGNS	1162
                        *:*:.***:***:* ..*****************:**:  *:.:::**:.

MDOA002080-RB      I-NHQDNRLEHELNHRGLSIQDDDTASINSYGSHKNRPFKDESHKGSAETIEGEEKRDVS	1208
AAEL023266-RG      ISNHQDNKLEHELNHRGMSLQDDDTASIKSYGSHKNRPFKDESHKGSAETMEGEEKRDVS	1209
AAEL023266-RE      ISNHQDNKLEHELNHRGMSLQDDDTASIKSYGSHKNRPFKDESHKGSAETMEGEEKRDVS	1206
AAEL023266-RH      ISNHQDNKLEHELNHRGMSLQDDDTASIKSYGSHKNRPFKDESHKGSAETMEGEEKRDVS	1198
AAEL023266-RF      ---------------------DDDTASIKSYGSHKNRPFKDESHKGSAETMEGEEKRDVS	1154
AAEL023266-RC      ISNHQDNKLEHELNHRGMSLQDDDTASIKSYGSHKNRPFKDESHKGSAETMEGEEKRDVS	1222
AAEL023266-RD      ---------------------DDDTASIKSYGSHKNRPFKDESHKGSAETMEGEEKRDVS	1173
AAEL023266-RI      ISNHQDNKLEHELNHRGMSLQDDDTASIKSYGSHKNRPFKDESHKGSAETMEGEEKRDVS	1174
AAEL023266-RM      ISNHQDNKLEHELNHRGMSLQDDDTASIKSYGSHKNRPFKDESHKGSAETMEGEEKRDVS	1188
AAEL023266-RA      ISNHQDNKLEHELNHRGMSLQDDDTASIKSYGSHKNRPFKDESHKGSAETMEGEEKRDVS	1197
AAEL023266-RB      ISNHQDNKLEHELNHRGMSLQDDDTASIKSYGSHKNRPFKDESHKGSAETMEGEEKRDVS	1199
AAEL023266-RL      ISNHQDNKLEHELNHRGMSLQDDDTASIKSYGSHKNRPFKDESHKGSAETMEGEEKRDVS	1229
AAEL023266-RJ      ISNHQDNKLEHELNHRGMSLQDDDTASIKSYGSHKNRPFKDESHKGSAETMEGEEKRDVS	1222
AAEL023266-RK      ISNHQDNKLEHELNHRGMSLQDDDTASIKSYGSHKNRPFKDESHKGSAETMEGEEKRDVS	1222
                                        *******:*********************:*********

MDOA002080-RB      KEDLGLDEELDEEAEGDEGQLDGDIIIHAQNDDEIIDDYPADCFPDSYYKKFPILAGDED	1268
AAEL023266-RG      KEDLGIDEELDDECDGEEGPLDGELIIHADE-DEVIEDSPADCCPDNCYKKFPVLAGDDD	1268
AAEL023266-RE      KEDLGIDEELDDECDGEEGPLDGELIIHADE-DEVIEDSPADCCPDNCYKKFPVLAGDDD	1265
AAEL023266-RH      KEDLGIDEELDDECDGEEGPLDGELIIHADE-DEVIEDSPADCCPDNCYKKFPVLAGDDD	1257
AAEL023266-RF      KEDLGIDEELDDECDGEEGPLDGELIIHADE-DEVIEDSPADCCPDNCYKKFPVLAGDDD	1213
AAEL023266-RC      KEDLGIDEELDDECDGEEGPLDGELIIHADE-DEVIEDSPADCCPDNCYKKFPVLAGDDD	1281
AAEL023266-RD      KEDLGIDEELDDECDGEEGPLDGELIIHADE-DEVIEDSPADCCPDNCYKKFPVLAGDDD	1232
AAEL023266-RI      KEDLGIDEELDDECDGEEGPLDGELIIHADE-DEVIEDSPADCCPDNCYKKFPVLAGDDD	1233
AAEL023266-RM      KEDLGIDEELDDECDGEEGPLDGELIIHADE-DEVIEDSPADCCPDNCYKKFPVLAGDDD	1247
AAEL023266-RA      KEDLGIDEELDDECDGEEGPLDGELIIHADE-DEVIEDSPADCCPDNCYKKFPVLAGDDD	1256
AAEL023266-RB      KEDLGIDEELDDECDGEEGPLDGELIIHADE-DEVIEDSPADCCPDNCYKKFPVLAGDDD	1258
AAEL023266-RL      KEDLGIDEELDDECDGEEGPLDGELIIHADE-DEVIEDSPADCCPDNCYKKFPVLAGDDD	1288
AAEL023266-RJ      KEDLGIDEELDDECDGEEGPLDGELIIHADE-DEVIEDSPADCCPDNCYKKFPVLAGDDD	1281
AAEL023266-RK      KEDLGIDEELDDECDGEEGPLDGELIIHADE-DEVIEDSPADCCPDNCYKKFPVLAGDDD	1281
                   *****:*****:*.:*:** ***::****:: **:*:* **** **. *****:****:*

MDOA002080-RB      SPFWQGWGNLRLKTFQLIENKYFETAVITMILMSSLALALEDVHLPDRPVMQDILYYMDR	1328
AAEL023266-RG      APFWQGWANLRLKTFQLIENKYFETAVITMILLSSLALALEDVHLPHRPILQDVLYYMDR	1328
AAEL023266-RE      APFWQGWANLRLKTFQLIENKYFETAVITMILLSSLALALEDVHLPHRPILQDVLYYMDR	1325
AAEL023266-RH      APFWQGWANLRLKTFQLIENKYFETAVITMILLSSLALALEDVHLPHRPILQDVLYYMDR	1317
AAEL023266-RF      APFWQGWANLRLKTFQLIENKYFETAVITMILLSSLALALEDVHLPHRPILQDVLYYMDR	1273
AAEL023266-RC      APFWQGWANLRLKTFQLIENKYFETAVITMILLSSLALALEDVHLPHRPILQDVLYYMDR	1341
AAEL023266-RD      APFWQGWANLRLKTFQLIENKYFETAVITMILLSSLALALEDVHLPHRPILQDVLYYMDR	1292
AAEL023266-RI      APFWQGWANLRLKTFQLIENKYFETAVITMILLSSLALALEDVHLPHRPILQDVLYYMDR	1293
AAEL023266-RM      APFWQGWANLRLKTFQLIENKYFETAVITMILLSSLALALEDVHLPHRPILQDVLYYMDR	1307
AAEL023266-RA      APFWQGWANLRLKTFQLIENKYFETAVITMILLSSLALALEDVHLPHRPILQDVLYYMDR	1316
AAEL023266-RB      APFWQGWANLRLKTFQLIENKYFETAVITMILLSSLALALEDVHLPHRPILQDVLYYMDR	1318
AAEL023266-RL      APFWQGWANLRLKTFQLIENKYFETAVITMILLSSLALALEDVHLPHRPILQDVLYYMDR	1348
AAEL023266-RJ      APFWQGWANLRLKTFQLIENKYFETAVITMILLSSLALALEDVHLPHRPILQDVLYYMDR	1341
AAEL023266-RK      APFWQGWANLRLKTFQLIENKYFETAVITMILLSSLALALEDVHLPHRPILQDVLYYMDR	1341
                   :******.************************:*************.**::**:******

MDOA002080-RB      IFTVIFFLEMLIKWLALGFKVYFTNAWCWLDFVIVMLSLINLVAVWSGLNDIAVFRSMRT	1388
AAEL023266-RG      IFTVIFFLEMLIKWLALGFRVYFTNAWCWLDFIIVMVSLINFVASLCGAGGIQAFKTMRT	1388
AAEL023266-RE      IFTVIFFLEMLIKWLALGFRVYFTNAWCWLDFIIVMVSLINFVASLCGAGGIQAFKTMRT	1385
AAEL023266-RH      IFTVIFFLEMLIKWLALGFRVYFTNAWCWLDFIIVMVSLINFVASLCGAGGIQAFKTMRT	1377
AAEL023266-RF      IFTVIFFLEMLIKWLALGFRVYFTNAWCWLDFIIVMVSLINFVASLCGAGGIQAFKTMRT	1333
AAEL023266-RC      IFTVIFFLEMLIKWLALGFRVYFTNAWCWLDFIIVMLSLINLTAIWVGAADIPAFRSMRT	1401
AAEL023266-RD      IFTVIFFLEMLIKWLALGFRVYFTNAWCWLDFIIVMVSLINFVASLCGAGGIQAFKTMRT	1352
AAEL023266-RI      IFTVIFFLEMLIKWLALGFRVYFTNAWCWLDFIIVMVSLINFVASLCGAGGIQAFKTMRT	1353
AAEL023266-RM      IFTVIFFLEMLIKWLALGFRVYFTNAWCWLDFIIVMVSLINFVASLCGAGGIQAFKTMRT	1367
AAEL023266-RA      IFTVIFFLEMLIKWLALGFRVYFTNAWCWLDFIIVMVSLINFVASLCGAGGIQAFKTMRT	1376
AAEL023266-RB      IFTVIFFLEMLIKWLALGFRVYFTNAWCWLDFIIVMVSLINFVASLCGAGGIQAFKTMRT	1378
AAEL023266-RL      IFTVIFFLEMLIKWLALGFRVYFTNAWCWLDFIIVMVSLINFVASLCGAGGIQAFKTMRT	1408
AAEL023266-RJ      IFTVIFFLEMLIKWLALGFRVYFTNAWCWLDFIIVMVSLINFVASLCGAGGIQAFKTMRT	1401
AAEL023266-RK      IFTVIFFLEMLIKWLALGFRVYFTNAWCWLDFIIVMVSLINFVASLCGAGGIQAFKTMRT	1401
                   *******************:************:***:****:.*   *  .* .*::***

MDOA002080-RB      LRALRPLRAVSRWEGMKVVVNALVQAIPSIFNVLLVCLIFWLIFAIMGVQLFAGKYFKCK	1448
AAEL023266-RG      LRALRPLRAMSRMQGMRVVVNALVQAIPSIFNVLLVCLIFWLIFAIMGVQLFAGKYFKCV	1448
AAEL023266-RE      LRALRPLRAMSRMQGMRVVVNALVQAIPSIFNVLLVCLIFWLIFAIMGVQLFAGKYFKCV	1445
AAEL023266-RH      LRALRPLRAMSRMQGMRVVVNALVQAIPSIFNVLLVCLIFWLIFAIMGVQLFAGKYFKCV	1437
AAEL023266-RF      LRALRPLRAMSRMQGMRVVVNALVQAIPSIFNVLLVCLIFWLIFAIMGVQLFAGKYFKCV	1393
AAEL023266-RC      LRALRPLRAVSRWEGMRVVVNALVQAIPSIFNVLLVCLIFWLIFAIMGVQLFAGKYFKCV	1461
AAEL023266-RD      LRALRPLRAMSRMQGMRVVVNALVQAIPSIFNVLLVCLIFWLIFAIMGVQLFAGKYFKCV	1412
AAEL023266-RI      LRALRPLRAMSRMQGMRVVVNALVQAIPSIFNVLLVCLIFWLIFAIMGVQLFAGKYFKCV	1413
AAEL023266-RM      LRALRPLRAMSRMQGMRVVVNALVQAIPSIFNVLLVCLIFWLIFAIMGVQLFAGKYFKCV	1427
AAEL023266-RA      LRALRPLRAMSRMQGMRVVVNALVQAIPSIFNVLLVCLIFWLIFAIMGVQLFAGKYFKCV	1436
AAEL023266-RB      LRALRPLRAMSRMQGMRVVVNALVQAIPSIFNVLLVCLIFWLIFAIMGVQLFAGKYFKCV	1438
AAEL023266-RL      LRALRPLRAMSRMQGMRVVVNALVQAIPSIFNVLLVCLIFWLIFAIMGVQLFAGKYFKCV	1468
AAEL023266-RJ      LRALRPLRAMSRMQGMRVVVNALVQAIPSIFNVLLVCLIFWLIFAIMGVQLFAGKYFKCV	1461
AAEL023266-RK      LRALRPLRAMSRMQGMRVVVNALVQAIPSIFNVLLVCLIFWLIFAIMGVQLFAGKYFKCV	1461
                   *********:** :**:******************************************

MDOA002080-RB      DGNDTVLSHEIIPNRNACKSENYTWENSAMNFDHVGNAYLCLFQVATFKGWIQIMNDAID	1508
AAEL023266-RG      DKNKTTLSHEIIPDVNACVAENYTWENSPMNFDHVGKAYLCLFQVATFKGWIQIMNDAID	1508
AAEL023266-RE      DKNKTTLSHEIIPDVNACVAENYTWENSPMNFDHVGKAYLCLFQVATFKGWIQIMNDAID	1505
AAEL023266-RH      DKNKTTLSHEIIPDVNACVAENYTWENSPMNFDHVGKAYLCLFQVATFKGWIQIMNDAID	1497
AAEL023266-RF      DKNKTTLSHEIIPDVNACVAENYTWENSPMNFDHVGKAYLCLFQVATFKGWIQIMNDAID	1453
AAEL023266-RC      DKNKTTLSHEIIPDVNACVAENYTWENSPMNFDHVGKAYLCLFQVATFKGWIQIMNDAID	1521
AAEL023266-RD      DKNKTTLSHEIIPDVNACVAENYTWENSPMNFDHVGKAYLCLFQVATFKGWIQIMNDAID	1472
AAEL023266-RI      DKNKTTLSHEIIPDVNACVAENYTWENSPMNFDHVGKAYLCLFQVATFKGWIQIMNDAID	1473
AAEL023266-RM      DKNKTTLSHEIIPDVNACVAENYTWENSPMNFDHVGKAYLCLFQVATFKGWIQIMNDAID	1487
AAEL023266-RA      DKNKTTLSHEIIPDVNACVAENYTWENSPMNFDHVGKAYLCLFQVATFKGWIQIMNDAID	1496
AAEL023266-RB      DKNKTTLSHEIIPDVNACVAENYTWENSPMNFDHVGKAYLCLFQVATFKGWIQIMNDAID	1498
AAEL023266-RL      DKNKTTLSHEIIPDVNACVAENYTWENSPMNFDHVGKAYLCLFQVATFKGWIQIMNDAID	1528
AAEL023266-RJ      DKNKTTLSHEIIPDVNACVAENYTWENSPMNFDHVGKAYLCLFQVATFKGWIQIMNDAID	1521
AAEL023266-RK      DKNKTTLSHEIIPDVNACVAENYTWENSPMNFDHVGKAYLCLFQVATFKGWIQIMNDAID	1521
                   * *.*.*******: *** :******** *******:***********************

MDOA002080-RB      SREVDKQPIRETNIYMYLYFVFFIIFGSFFTLNLFIGVIIDNFNEQKKKAGGSLEMFMTE	1568
AAEL023266-RG      SREVGKQPIRETNIYMYLYFVFFIIFGSFFTLNLFIGVIIDNFNEQKKKAGGSLEMFMTE	1568
AAEL023266-RE      SREVGKQPIRETNIYMYLYFVFFIIFGSFFTLNLFIGVIIDNFNEQKKKAGGSLEMFMTE	1565
AAEL023266-RH      SREVGKQPIRETNIYMYLYFVFFIIFGSFFTLNLFIGVIIDNFNEQKKKAGGSLEMFMTE	1557
AAEL023266-RF      SREVGKQPIRETNIYMYLYFVFFIIFGSFFTLNLFIGVIIDNFNEQKKKAGGSLEMFMTE	1513
AAEL023266-RC      SREVGKQPIRETNIYMYLYFVFFIIFGSFFTLNLFIGVIIDNFNEQKKKAGGSLEMFMTE	1581
AAEL023266-RD      SREVGKQPIRETNIYMYLYFVFFIIFGSFFTLNLFIGVIIDNFNEQKKKAGGSLEMFMTE	1532
AAEL023266-RI      SREVGKQPIRETNIYMYLYFVFFIIFGSFFTLNLFIGVIIDNFNEQKKKAGGSLEMFMTE	1533
AAEL023266-RM      SREVGKQPIRETNIYMYLYFVFFIIFGSFFTLNLFIGVIIDNFNEQKKKAGGSLEMFMTE	1547
AAEL023266-RA      SREVGKQPIRETNIYMYLYFVFFIIFGSFFTLNLFIGVIIDNFNEQKKKAGGSLEMFMTE	1556
AAEL023266-RB      SREVGKQPIRETNIYMYLYFVFFIIFGSFFTLNLFIGVIIDNFNEQKKKAGGSLEMFMTE	1558
AAEL023266-RL      SREVGKQPIRETNIYMYLYFVFFIIFGSFFTLNLFIGVIIDNFNEQKKKAGGSLEMFMTE	1588
AAEL023266-RJ      SREVGKQPIRETNIYMYLYFVFFIIFGSFFTLNLFIGVIIDNFNEQKKKAGGSLEMFMTE	1581
AAEL023266-RK      SREVGKQPIRETNIYMYLYFVFFIIFGSFFTLNLFIGVIIDNFNEQKKKAGGSLEMFMTE	1581
                   ****.*******************************************************

MDOA002080-RB      DQKKYYNAMKKMGSKKPLKAIPRPRWRPQAIVFEIVTDKKFDIIIMLFIGLNMFTMTLDR	1628
AAEL023266-RG      DQKKYYNAMKKMGSKKPLKAIPRPRWRPQAIVFEIVTNKKFDMIIMLFIGFNMLTMTLDH	1628
AAEL023266-RE      DQKKYYNAMKKMGSKKPLKAIPRPRWRPQAIVFEIVTNKKFDMIIMLFIGFNMLTMTLDH	1625
AAEL023266-RH      DQKKYYNAMKKMGSKKPLKAIPRPRWRPQAIVFEIVTNKKFDMIIMLFIGFNMLTMTLDH	1617
AAEL023266-RF      DQKKYYNAMKKMGSKKPLKAIPRPRWRPQAIVFEIVTNKKFDMIIMLFIGFNMLTMTLDH	1573
AAEL023266-RC      DQKKYYNAMKKMGSKKPLKAIPRPRWRPQAIVFEIVTNKKFDMIIMLFIGFNMLTMTLDH	1641
AAEL023266-RD      DQKKYYNAMKKMGSKKPLKAIPRPRWRPQAIVFEIVTNKKFDMIIMLFIGFNMLTMTLDH	1592
AAEL023266-RI      DQKKYYNAMKKMGSKKPLKAIPRPRWRPQAIVFEIVTNKKFDMIIMLFIGFNMLTMTLDH	1593
AAEL023266-RM      DQKKYYNAMKKMGSKKPLKAIPRPRWRPQAIVFEIVTNKKFDMIIMLFIGFNMLTMTLDH	1607
AAEL023266-RA      DQKKYYNAMKKMGSKKPLKAIPRPRWRPQAIVFEIVTNKKFDMIIMLFIGFNMLTMTLDH	1616
AAEL023266-RB      DQKKYYNAMKKMGSKKPLKAIPRPRWRPQAIVFEIVTNKKFDMIIMLFIGFNMLTMTLDH	1618
AAEL023266-RL      DQKKYYNAMKKMGSKKPLKAIPRPRWRPQAIVFEIVTNKKFDMIIMLFIGFNMLTMTLDH	1648
AAEL023266-RJ      DQKKYYNAMKKMGSKKPLKAIPRPRWRPQAIVFEIVTNKKFDMIIMLFIGFNMLTMTLDH	1641
AAEL023266-RK      DQKKYYNAMKKMGSKKPLKAIPRPRWRPQAIVFEIVTNKKFDMIIMLFIGFNMLTMTLDH	1641
                   *************************************:****:*******:**:*****:

MDOA002080-RB      YDASEAYNNVLDKLNGIFVVIFSGECLLKIFALRYHYFKEPWNLFDVVVVILSILGLVLS	1688
AAEL023266-RG      YKQTDTFSAVLDYLNMIFICIFSSECLMKIFALRYHYFIEPWNLFDFVVVILSILGLVLS	1688
AAEL023266-RE      YKQTDTFSAVLDYLNMIFICIFSSECLMKIFALRYHYFIEPWNLFDFVVVILSILGLVLS	1685
AAEL023266-RH      YKQTDTFSAVLDYLNMIFICIFSSECLMKIFALRYHYFIEPWNLFDFVVVILSILGLVLS	1677
AAEL023266-RF      YKQTDTFSAVLDYLNMIFICIFSSECLMKIFALRYHYFIEPWNLFDFVVVILSILGLVLS	1633
AAEL023266-RC      YKQTDTFSAVLDYLNMIFICIFSSECLMKIFALRYHYFIEPWNLFDFVVVILSILGLVLS	1701
AAEL023266-RD      YKQTDTFSAVLDYLNMIFICIFSSECLMKIFALRYHYFIEPWNLFDFVVVILSILGLVLS	1652
AAEL023266-RI      YKQTDTFSAVLDYLNMIFICIFSSECLMKIFALRYHYFIEPWNLFDFVVVILSILGLVLS	1653
AAEL023266-RM      YKQTDTFSAVLDYLNMIFICIFSSECLMKIFALRYHYFIEPWNLFDFVVVILSILGLVLS	1667
AAEL023266-RA      YKQTDTFSAVLDYLNMIFICIFSSECLMKIFALRYHYFIEPWNLFDFVVVILSILGLVLS	1676
AAEL023266-RB      YKQTDTFSAVLDYLNMIFICIFSSECLMKIFALRYHYFIEPWNLFDFVVVILSILGLVLS	1678
AAEL023266-RL      YKQTDTFSAVLDYLNMIFICIFSSECLMKIFALRYHYFIEPWNLFDFVVVILSILGLVLS	1708
AAEL023266-RJ      YKQTDTFSAVLDYLNMIFICIFSSECLMKIFALRYHYFIEPWNLFDFVVVILSILGLVLS	1701
AAEL023266-RK      YKQTDTFSAVLDYLNMIFICIFSSECLMKIFALRYHYFIEPWNLFDFVVVILSILGLVLS	1701
                   *. ::::. *** ** **: ***.***:********** *******.*************

MDOA002080-RB      DIIEKYFVSPTLLRVVRVAKVGRVLRLVKGAKGIRTLLFALAMSLPALFNICLLLFLVMF	1748
AAEL023266-RG      DLIEKYFVSPTLLRVVRVAKVGRVLRLVKGAKGIRTLLFALAMSLPALFNICLLLFLVMF	1748
AAEL023266-RE      DLIEKYFVSPTLLRVVRVAKVGRVLRLVKGAKGIRTLLFALAMSLPALFNICLLLFLVMF	1745
AAEL023266-RH      DLIEKYFVSPTLLRVVRVAKVGRVLRLVKGAKGIRTLLFALAMSLPALFNICLLLFLVMF	1737
AAEL023266-RF      DLIEKYFVSPTLLRVVRVAKVGRVLRLVKGAKGIRTLLFALAMSLPALFNICLLLFLVMF	1693
AAEL023266-RC      DLIEKYFVSPTLLRVVRVAKVGRVLRLVKGAKGIRTLLFALAMSLPALFNICLLLFLVMF	1761
AAEL023266-RD      DLIEKYFVSPTLLRVVRVAKVGRVLRLVKGAKGIRTLLFALAMSLPALFNICLLLFLVMF	1712
AAEL023266-RI      DLIEKYFVSPTLLRVVRVAKVGRVLRLVKGAKGIRTLLFALAMSLPALFNICLLLFLVMF	1713
AAEL023266-RM      DLIEKYFVSPTLLRVVRVAKVGRVLRLVKGAKGIRTLLFALAMSLPALFNICLLLFLVMF	1727
AAEL023266-RA      DLIEKYFVSPTLLRVVRVAKVGRVLRLVKGAKGIRTLLFALAMSLPALFNICLLLFLVMF	1736
AAEL023266-RB      DLIEKYFVSPTLLRVVRVAKVGRVLRLVKGAKGIRTLLFALAMSLPALFNICLLLFLVMF	1738
AAEL023266-RL      DLIEKYFVSPTLLRVVRVAKVGRVLRLVKGAKGIRTLLFALAMSLPALFNICLLLFLVMF	1768
AAEL023266-RJ      DLIEKYFVSPTLLRVVRVAKVGRVLRLVKGAKGIRTLLFALAMSLPALFNICLLLFLVMF	1761
AAEL023266-RK      DLIEKYFVSPTLLRVVRVAKVGRVLRLVKGAKGIRTLLFALAMSLPALFNICLLLFLVMF	1761
                   *:**********************************************************

MDOA002080-RB      IFAIFGMSFFMHVKEKSGINAVYNFKTFGQSMILLFQMSTSAGWDGVLDAIINEEDCDPP	1808
AAEL023266-RG      IFAIFGMSFFMHVKDKSGLDDVYNFKTFGQSMILLFQMSTSAGWDGVLDGIINEDECLPP	1808
AAEL023266-RE      IFAIFGMSFFMHVKDKSGLDDVYNFKTFGQSMILLFQMSTSAGWDGVLDGIINEDECLPP	1805
AAEL023266-RH      IFAIFGMSFFMHVKDKSGLDDVYNFKTFGQSMILLFQMSTSAGWDGVLDGIINEDECLPP	1797
AAEL023266-RF      IFAIFGMSFFMHVKDKSGLDDVYNFKTFGQSMILLFQMSTSAGWDGVLDGIINEDECLPP	1753
AAEL023266-RC      IFAIFGMSFFMHVKDKSGLDDVYNFKTFGQSMILLFQMSTSAGWDGVLDGIINEDECLPP	1821
AAEL023266-RD      IFAIFGMSFFMHVKDKSGLDDVYNFKTFGQSMILLFQMSTSAGWDGVLDGIINEDECLPP	1772
AAEL023266-RI      IFAIFGMSFFMHVKDKSGLDDVYNFKTFGQSMILLFQMSTSAGWDGVLDGIINEDECLPP	1773
AAEL023266-RM      IFAIFGMSFFMHVKDKSGLDDVYNFKTFGQSMILLFQMSTSAGWDGVLDGIINEDECLPP	1787
AAEL023266-RA      IFAIFGMSFFMHVKDKSGLDDVYNFKTFGQSMILLFQMSTSAGWDGVLDGIINEDECLPP	1796
AAEL023266-RB      IFAIFGMSFFMHVKDKSGLDDVYNFKTFGQSMILLFQMSTSAGWDGVLDGIINEDECLPP	1798
AAEL023266-RL      IFAIFGMSFFMHVKDKSGLDDVYNFKTFGQSMILLFQMSTSAGWDGVLDGIINEDECLPP	1828
AAEL023266-RJ      IFAIFGMSFFMHVKDKSGLDDVYNFKTFGQSMILLFQMSTSAGWDGVLDGIINEDECLPP	1821
AAEL023266-RK      IFAIFGMSFFMHVKDKSGLDDVYNFKTFGQSMILLFQMSTSAGWDGVLDGIINEDECLPP	1821
                   **************:***:: ****************************.****::* **

MDOA002080-RB      DNDKGYPGNCGSATVGITFLLSYLVISFLIVINMYIAVILENYSQATEDVQEGLTDDDYD	1868
AAEL023266-RG      DNDKGYPGNCGSATIGITYLLAYLVISFLIVINMYIAVILENYSQATEDVQEGLTDDDYD	1868
AAEL023266-RE      DNDKGYPGNCGSATIGITYLLAYLVISFLIVINMYIAVILENYSQATEDVQEGLTDDDYD	1865
AAEL023266-RH      DNDKGYPGNCGSATIGITYLLAYLVISFLIVINMYIAVILENYSQATEDVQEGLTDDDYD	1857
AAEL023266-RF      DNDKGYPGNCGSATIGITYLLAYLVISFLIVINMYIAVILENYSQATEDVQEGLTDDDYD	1813
AAEL023266-RC      DNDKGYPGNCGSATIGITYLLAYLVISFLIVINMYIAVILENYSQATEDVQEGLTDDDYD	1881
AAEL023266-RD      DNDKGYPGNCGSATIGITYLLAYLVISFLIVINMYIAVILENYSQATEDVQEGLTDDDYD	1832
AAEL023266-RI      DNDKGYPGNCGSATIGITYLLAYLVISFLIVINMYIAVILENYSQATEDVQEGLTDDDYD	1833
AAEL023266-RM      DNDKGYPGNCGSATIGITYLLAYLVISFLIVINMYIAVILENYSQATEDVQEGLTDDDYD	1847
AAEL023266-RA      DNDKGYPGNCGSATIGITYLLAYLVISFLIVINMYIAVILENYSQATEDVQEGLTDDDYD	1856
AAEL023266-RB      DNDKGYPGNCGSATIGITYLLAYLVISFLIVINMYIAVILENYSQATEDVQEGLTDDDYD	1858
AAEL023266-RL      DNDKGYPGNCGSATIGITYLLAYLVISFLIVINMYIAVILENYSQATEDVQEGLTDDDYD	1888
AAEL023266-RJ      DNDKGYPGNCGSATIGITYLLAYLVISFLIVINMYIAVILENYSQATEDVQEGLTDDDYD	1881
AAEL023266-RK      DNDKGYPGNCGSATIGITYLLAYLVISFLIVINMYIAVILENYSQATEDVQEGLTDDDYD	1881
                   **************:***:**:**************************************

MDOA002080-RB      MYYEIWQQFDPEGTQYIRYDQLSEFLDVLEPPLQIHKPNKYKIISMDMPICRGDMMYCVD	1928
AAEL023266-RG      MYYEIWQQFDPDGTQYIRYDQLSDFLDVLEPPLQIHKPNKYKIISMDIPICRGDMMFCVD	1928
AAEL023266-RE      MYYEIWQQFDPDGTQYIRYDQLSDFLDVLEPPLQIHKPNKYKIISMDIPICRGDMMFCVD	1925
AAEL023266-RH      MYYEIWQQFDPDGTQYIRYDQLSDFLDVLEPPLQIHKPNKYKIISMDIPICRGDMMFCVD	1917
AAEL023266-RF      MYYEIWQQFDPDGTQYIRYDQLSDFLDVLEPPLQIHKPNKYKIISMDIPICRGDMMFCVD	1873
AAEL023266-RC      MYYEIWQQFDPDGTQYIRYDQLSDFLDVLEPPLQIHKPNKYKIISMDIPICRGDMMFCVD	1941
AAEL023266-RD      MYYEIWQQFDPDGTQYIRYDQLSDFLDVLEPPLQIHKPNKYKIISMDIPICRGDMMFCVD	1892
AAEL023266-RI      MYYEIWQQFDPDGTQYIRYDQLSDFLDVLEPPLQIHKPNKYKIISMDIPICRGDMMFCVD	1893
AAEL023266-RM      MYYEIWQQFDPDGTQYIRYDQLSDFLDVLEPPLQIHKPNKYKIISMDIPICRGDMMFCVD	1907
AAEL023266-RA      MYYEIWQQFDPDGTQYIRYDQLSDFLDVLEPPLQIHKPNKYKIISMDIPICRGDMMFCVD	1916
AAEL023266-RB      MYYEIWQQFDPDGTQYIRYDQLSDFLDVLEPPLQIHKPNKYKIISMDIPICRGDMMFCVD	1918
AAEL023266-RL      MYYEIWQQFDPDGTQYIRYDQLSDFLDVLEPPLQIHKPNKYKIISMDIPICRGDMMFCVD	1948
AAEL023266-RJ      MYYEIWQQFDPDGTQYIRYDQLSDFLDVLEPPLQIHKPNKYKIISMDIPICRGDMMFCVD	1941
AAEL023266-RK      MYYEIWQQFDPDGTQYIRYDQLSDFLDVLEPPLQIHKPNKYKIISMDIPICRGDMMFCVD	1941
                   ***********:***********:***********************:********:***

MDOA002080-RB      ILDALTKDFFARKGNPIEETGEIGEIAARPDTEGYDPVSSTLWRQREEYCAKLIQNAWRR	1988
AAEL023266-RG      ILDALTKDFFARKGNPIEETAELGEVQARPDEVGYEPVSSTLWRQREEYCARVIQHAWRK	1988
AAEL023266-RE      ILDALTKDFFARKGNPIEETAELGEVQARPDEVGYEPVSSTLWRQREEYCARVIQHAWRK	1985
AAEL023266-RH      ILDALTKDFFARKGNPIEETAELGEVQARPDEVGYEPVSSTLWRQREEYCARVIQHAWRK	1977
AAEL023266-RF      ILDALTKDFFARKGNPIEETAELGEVQARPDEVGYEPVSSTLWRQREEYCARVIQHAWRK	1933
AAEL023266-RC      ILDALTKDFFARKGNPIEETAELGEVQARPDEVGYEPVSSTLWRQREEYCARVIQHAWRK	2001
AAEL023266-RD      ILDALTKDFFARKGNPIEETAELGEVQARPDEVGYEPVSSTLWRQREEYCARVIQHAWRK	1952
AAEL023266-RI      ILDALTKDFFARKGNPIEETAELGEVQARPDEVGYEPVSSTLWRQREEYCARVIQHAWRK	1953
AAEL023266-RM      ILDALTKDFFARKGNPIEETAELGEVQARPDEVGYEPVSSTLWRQREEYCARVIQHAWRK	1967
AAEL023266-RA      ILDALTKDFFARKGNPIEETAELGEVQARPDEVGYEPVSSTLWRQREEYCARVIQHAWRK	1976
AAEL023266-RB      ILDALTKDFFARKGNPIEETAELGEVQARPDEVGYEPVSSTLWRQREEYCARVIQHAWRK	1978
AAEL023266-RL      ILDALTKDFFARKGNPIEETAELGEVQARPDEVGYEPVSSTLWRQREEYCARVIQHAWRK	2008
AAEL023266-RJ      ILDALTKDFFARKGNPIEETAELGEVQARPDEVGYEPVSSTLWRQREEYCARVIQHAWRK	2001
AAEL023266-RK      ILDALTKDFFARKGNPIEETAELGEVQARPDEVGYEPVSSTLWRQREEYCARVIQHAWRK	2001
                   ********************.*:**: ****  **:***************::**:***:

MDOA002080-RB      YKNGPPQEGDEG----EAAGGEDGAEGGEGEGGSGGGGGGGDDGGSATGATAAAAGATSP	2044
AAEL023266-RG      HKERQAGGGGGDDTDADACDNDDGDDGGGGAGDGGSAGGGGVT-SPGVGSGSIVGGGTTP	2047
AAEL023266-RE      HKERQAGGGGGDDTDADACDNDDGDDGGGGAGDGGSAGGGGVT-SPGVGSGSIVGGGTTP	2044
AAEL023266-RH      HKERQAGGGGGDDTDADACDNDDGDDGGGGAGDGGSAGGGGVT-SPGVGSGSIVGGGTTP	2036
AAEL023266-RF      HKERQAGGGGGDDTDADACDNDDGDDGGGGAGDGGSAGGGGVT-SPGVGSGSIVGGGTTP	1992
AAEL023266-RC      HKERQAGGGGGDDTDADACDNDDGDDGGGGAGDGGSAGGGGVT-SPGVGSGSIVGGGTTP	2060
AAEL023266-RD      HKERQAGGGGGDDTDADACDNDDGDDGGGGAGDGGSAGGGGVT-SPGVGSGSIVGGGTTP	2011
AAEL023266-RI      HKERQAGGGGGDDTDADACDNDDGDDGGGGAGDGGSAGGGGVT-SPGVGSGSIVGGGTTP	2012
AAEL023266-RM      HKERQAGGGGGDDTDADACDNDDGDDGGGGAGDGGSAGGGGVT-SPGVGSGSIVGGGTTP	2026
AAEL023266-RA      HKERQAGGGGGDDTDADACDNDDGDDGGGGAGDGGSAGGGGVT-SPGVGSGSIVGGGTTP	2035
AAEL023266-RB      HKERQAGGGGGDDTDADACDNDDGDDGGGGAGDGGSAGGGGVT-SPGVGSGSIVGGGTTP	2037
AAEL023266-RL      HKERQAGGGGGDDTDADACDNDDGDDGGGGAGDGGSAGGGGVT-SPGVGSGSIVGGGTTP	2067
AAEL023266-RJ      HKERQAGGGGGDDTDADACDNDDGDDGGGGAGDGGSAGGGGVT-SPGVGSGSIVGGGTTP	2060
AAEL023266-RK      HKERQAGGGGGDDTDADACDNDDGDDGGGGAGDGGSAGGGGVT-SPGVGSGSIVGGGTTP	2060
                   :*:     *. .    :*...:** :** * *..*..****   . ..*: : ..*.*:*

MDOA002080-RB      SDPDAGEA--DGASVGGPLSPGCV---SGGSNGRQTAVLVESDGFVTKNGHKVVIHSRSP	2099
AAEL023266-RG      GSGGGGSQANLGIVVEHNLSPKESPDGNNDPQGRQTAVLVESDGFVTKNGHRVVIHSRSP	2107
AAEL023266-RE      GSGGGGSQANLGIVVEHNLSPKESPDGNNDPQGRQTAVLVESDGFVTKNGHRVVIHSRSP	2104
AAEL023266-RH      GSGGGGSQANLGIVVEHNLSPKESPDGNNDPQGRQTAVLVESDGFVTKNGHRVVIHSRSP	2096
AAEL023266-RF      GSGGGGSQANLGIVVEHNLSPKESPDGNNDPQGRQTAVLVESDGFVTKNGHRVVIHSRSP	2052
AAEL023266-RC      GSGGGGSQANLGIVVEHNLSPKESPDGNNDPQGRQTAVLVESDGFVTKNGHRVVIHSRSP	2120
AAEL023266-RD      GSGGGGSQANLGIVVEHNLSPKESPDGNNDPQGRQTAVLVESDGFVTKNGHRVVIHSRSP	2071
AAEL023266-RI      GSGGGGSQANLGIVVEHNLSPKESPDGNNDPQGRQTAVLVESDGFVTKNGHRVVIHSRSP	2072
AAEL023266-RM      GSGGGGSQANLGIVVEHNLSPKESPDGNNDPQGRQTAVLVESDGFVTKNGHRVVIHSRSP	2086
AAEL023266-RA      GSGGGGSQANLGIVVEHNLSPKESPDGNNDPQGRQTAVLVESDGFVTKNGHRVVIHSRSP	2095
AAEL023266-RB      GSGGGGSQANLGIVVEHNLSPKESPDGNNDPQGRQTAVLVESDGFVTKNGHRVVIHSRSP	2097
AAEL023266-RL      GSGGGGSQANLGIVVEHNLSPKESPDGNNDPQGRQTAVLVESDGFVTKNGHRVVIHSRSP	2127
AAEL023266-RJ      GSGGGGSQANLGIVVEHNLSPKESPDGNNDPQGRQTAVLVESDGFVTKNGHRVVIHSRSP	2120
AAEL023266-RK      GSGGGGSQANLGIVVEHNLSPKESPDGNNDPQGRQTAVLVESDGFVTKNGHRVVIHSRSP	2120
                   .. ..*.    *  *   ***      ... :*******************:********

MDOA002080-RB      SITSRTADV	2108
AAEL023266-RG      SITSRSADV	2116
AAEL023266-RE      SITSRSADV	2113
AAEL023266-RH      SITSRSADV	2105
AAEL023266-RF      SITSRSADV	2061
AAEL023266-RC      SITSRSADV	2129
AAEL023266-RD      SITSRSADV	2080
AAEL023266-RI      SITSRSADV	2081
AAEL023266-RM      SITSRSADV	2095
AAEL023266-RA      SITSRSADV	2104
AAEL023266-RB      SITSRSADV	2106
AAEL023266-RL      SITSRSADV	2136
AAEL023266-RJ      SITSRSADV	2129
AAEL023266-RK      SITSRSADV	2129
                   *****:***
```
